# Gram-positive bacteria secrete RNA aptamers to activate human STING for IL-1β release

**DOI:** 10.1101/2021.07.21.453173

**Authors:** Shivalee N. Duduskar, Mohamed Ghait, Martin Westermann, Huijuan Guo, Anuradha Ramoji, Christoph Sponholz, Bianca Göhrig, Tony Bruns, Ute Neugebauer, Jurgen Popp, Lorena P N Tuchscherr, Bettina Löffler, Nico Ueberschaar, Christine Beemelmanns, Mervyn Singer, Michael Bauer, Sachin D. Deshmukh

## Abstract

Molecular mechanisms through which Gram-positive bacteria induce the canonical inflammasome are poorly understood. Here, we studied the effects of Group B streptococci (GBS) and *Staphylococcus aureus* (SA) on inflammasome activation in human macrophages. Dinucleotide binding small RNA aptamers released by SA and GBS were shown to trigger increased IL-1β generation by inflammasomes. The stimulator of interferon genes-STING as a central mediator of innate immune responses has been identified as the key target of pathogenic RNA. Multi-lamellar lipid bodies (MLBs) produced by SA function as vehicles for the RNA aptamers. Notably, expression of RNA aptamers is controlled by an accessory gene regulator quorum sensing system of the bacteria. These findings have been translated to patients with Gram-positive sepsis showing hallmarks of MLB-RNA-mediated inflammasome activation. Together our findings may provide a new perspective for the pathogenicity of Gram-positive bacterial infection in man.

## INTRODUCTION

Antimicrobial activity triggered by exposure to microbes follows activation of germ line-encoded pattern recognition receptors (PRRs) upon detection of pathogen-associated molecular patterns (PAMPs) or self-encoded damage-associated molecular patterns (DAMPs) (1). Inflammasomes are multimeric complexes assembled by PRRs in the cytosol of host cells (2). NLRP3 is the best characterized inflammasome; it consists of three major components: a cytoplasmic sensor NLRP3, an adaptor ASC (apoptosis-associated speck-like protein with CARD domain), and an interleukin-1β converting enzyme pro-caspase-1 (3). NLRP3 and ASC together promote activation and cleavage of pro-caspase-1 which subsequently cleaves and matures pro-IL-1β and pro-IL-18 (3, 4). Microbial ligands such as lipopolysaccharide (LPS) and nucleic acids are potent activators of PRRs at both the cell surface and within the cytosol of macrophages and dendritic cells (4). Much progress has been made in understanding how DNA and RNA trigger host pro-inflammatory immune responses. However, how these nucleic acids (in particular RNA) are sensed and thus activate the inflammasome remains unclear.

The endoplasmic reticulum (ER) resident transmembrane protein, stimulator of interferon genes (STING) plays a crucial role in the innate immune response against pathogen infections (5). On recognition of the cytosolic DNA, cyclic GMP-AMP synthase (cGAS) catalyses the synthesis of cGAMP which, in turn, binds to and activates STING (6). Activated STING exits the ER to the Golgi apparatus, activating TBK1 which uses STING as a scaffold to phosphorylate and activate the transcriptional factor IRF3. Upon activation STING translocates to the lysosome, resulting in a drop of cytoplasmic K^+^ due to lysosomal-mediated cell death. This subsequent fall in cytoplasmic K^+^ triggers the NLRP3 inflammasome, activating caspase-1 to release IL-1β (7, 8). A recent study reported that inflammasome activation in human cells is initiated by STING upstream of NLRP3 (9). STING-dependent interferon responses are induced by bacteria derived second messenger, cyclic dinucleotides (CDN) (10), bacterially derived second messenger molecules (11, 12). CDN are delivered into the cytosol by bacterial pathogens (13, 14) that bind to STING (15) for release of interferons. The existence of CdnP enzyme anchored in extracellular cell wall was recently shown to have phosphodiesterase activity towards cyclic-di-AMP in GBS. In the absence of CdnP the host overproduced type-I IFN leading to bacterial clearance (16). Bacterial CDN bind and activate STING for interferon responses, but it remains unclear how STING could still initiate NLRP3 responses for IL-1β release.

*S. aureus* and Group-B streptococci (GBS) are the most common opportunistic bacteria causing adult and neonatal sepsis, respectively (17, 18). The presence of pathogenicity-related genes including protein toxins and bacterial RNA are involved in induction of IL-1β following infection by Gram-positive bacteria (19, 20). Over 90% of clinical isolates of *S. aureus* produce the golden carotenoid pigment, staphyloxanthin. Production of staphyloxanthin is key to *S. aureus* virulence; bacteria lacking the capacity for staphyloxanthin synthesis are non-pathogenic and fail to induce host cell death (21, 22, 23). Similar observations were made for granadaene, a structurally bio-similar lipid toxin present in many clinical isolates of GBS causing sepsis and endocarditis. Notably, strains lacking granadaene are incapable of inducing cell death and translocation across the placenta; nevertheless stabilization of lipids was necessary for granadaene (24). Expression of numerous virulence factors is necessary for infections caused by *S. aureus* with the coordinated action of several regulators including two component systems:, transcriptional regulatory proteins and regulatory RNAIII (25). These regulatory networks are controlled by multiple trans-acting modulators including proteins, secondary metabolites, small peptides and RNAs. The ability of RNA to regulate these virulence factors in *S. aureus* has been widely studied (26). The 5’ untranslated regions of mRNA and small non-coding RNA are the known RNA-based regulators within *S.* aureus (27). Small trans-acting non-coding RNAs, referred to as small RNAs (sRNAs), play a key role in RNA-dependent regulation that enables *S. aureus* to express virulence genes during infection (27). How small RNAs are involved in inflammasome activation for release of IL-1β has yet to be studied.

Moreover, bacterial RNA is also a potent activator of cytokines and can induce the cytosolic inflammasome through the NLRP3 scaffold secreting IL-1β (20, 28, 29). Immune activating bacterial RNA contains mRNA, tRNA and three different sizes of rRNA (23s, 16s and 5s). Only mRNA is sensed by murine bone marrow-derived macrophages (BMDMs) while all other types are sensed by human macrophages (30). Different structural moieties of bacterial RNA (including µRNA) can induce the inflammasome via caspase-1 (30, 31). The RNA binding to small molecule ligands are cis-regulatory RNAs commonly found in the 5‘ untranslated region of mRNA (33). How RNA, in combination with CDN, activates the inflammasome has not yet been studied.

How the two major components, namely, lipid toxins and RNA, activate the inflammasome in bacterial infections to release IL-1β is not well understood. Moreover, how PAMPs get delivered to cytosolic receptors is unknown for Gram-positive bacteria. One such delivery system involving outer membrane extracellular vesicles (OMV) from Gram-negative bacteria can activate the non-canonical inflammasome (32, 34). Release of extracellular vesicles (EVs) from Gram-positive bacteria has been previously reported (35) yet their exact molecular structure and role in pathogenicity, including inflammasome activation remain poorly understood.

In this current study, we sought to explore how Gram-positive bacteria deliver RNA to activate cytosolic inflammasome in human macrophages. We have structurally characterized Gram-positive bacterial vesicles (multi lamellar lipid bodies MLBs) which show alternatively structured lipid assemblies. MLBs enriched in RNA aptamers activate the canonical inflammasome pathway via STING for release of IL-1β. In SA, expression of MLB specific

µRNA aptamers is controlled by an accessory gene regulator quorum sensing system. We also characterized the functional role of the staphyloxanthin type of lipids in delivery of PAMPs to cytosolic inflammasome receptors. Taken together, this study highlights the importance of STING-specific CDN binding to RNA aptamer delivered by Gram-positive bacteria to activate the inflammasome in relevance to human sepsis.

## RESULTS

### *S. aureus* and GBS induces the canonical inflammasome via STING and bacterial RNA

Highly pathogenic bacteria such as *S. aureus* and GBS evoke a pronounced NLRP3-ASC-caspase-1 dependent inflammasome activation in macrophages; however, this only occurs upon infection with live but not heat-killed bacteria (38). Varied stimuli including microbial products such as muramyl dipeptide, pore-forming toxins, and bacterial and viral RNA can activate the NLRP3 inflammasome (39). Nucleic acids within the cytoplasm can activate the cGAS-STING pathway leading to induction of interferon (IFN) and anti-viral immunity (5, 6). Considering the importance of IFN production via the cGAS-STING pathway, we investigated production of caspase-1 dependent cytokines through STING.

Human primary blood-derived macrophages and differentiated THP1 macrophage cells which activate both canonical and non-canonical inflammasome pathways (37) were used to examine activation by Gram-positive bacteria. Comparison was made against a genetically deleted panel of known signalling molecules within THP1 macrophages. Consistent with previous literature on non-canonical inflammasome activation by cytoplasmic LPS, we found that cell death was caspase-4 dependent but caspase-1 independent **(Supplemental Fig. 1A, 1B)**. On the other hand, nigericin, a potent inducer of the canonical inflammasome (38) was caspase-1 dependent but caspase-4 independent **(Supplemental Fig. 1C)**. This underlines the usability of our CRISPR cell lines.

Of note, subsequent experiments with SA and GBS infections led to classical NLRP3 inflammasome activation. The release of mature IL-1β was solely dependent on caspase-1 through an adaptor ASC, but independent of caspase-4 **(Fig. 1A, 1B)**. Moreover, activation of caspase-1 was independent of cGAS but STING dependent **(Fig. 1C, 1D)**. STING deficient macrophages infected with GBS and SA showed reduction in IL-1β secreation and caspase-1 activation when compared with cGAS deficient and wild type macrophages **(Fig. 1E)**. The effect of STING deficiency was retrieved by transgenic expression of a wild type variant of STING^R232^ **(Fig. 1F)**. On the other hand, inflammasome-independent cytokines such as TNF- α showed no differences in the absence of either STING or cGAS **(Supplemental Fig. 1D)**. Taking these results further we utilized a recently discovered, highly potent and selective small molecule antagonist (H-151) of STING protein (40). IL-1β production in human primary blood derived macrophages was abrogated by 80% following infection with GBS and *S. aureus* when pre-treated with H-151 **(Fig. 1G, 1H)**. In contrast to this IL-1β response, H- 151 pre-treatment had no effect on TNF-α production **(Supplemental Fig. 1E, 1F)**. These inflammasome activation findings contrast with previous findings for interferon production following bacterial infection that often requires both cGAS and STING (6, 16). As the significance of cGAS-STING in interferon production is well established, we queried which stimuli produced by Gram-positive bacteria could activate STING in the process of IL-1β production. As anticipated, cytosolic RNA stimulation resulted in strong caspase-1 inflammasome activation with release of IL-1β **(Fig. 1I)**. To confirm that RNA can act as a potent activator of the canonical inflammasome pathway and IL-1β release, cells were treated with RNase A and then infected with SA. Consistent with prior literature (20), RNase A treated cells showed significantly reduced IL-1β secretion **(Fig. 1J)**. Cells treated with DNase I and heat inactivated RNase A were used as controls and these were unable to show differences in IL-1β secretion **(Fig. 1K, 1L)**. The above results generate a new paradigm where by Gram-positive bacterial RNA can activate the canonical inflammasome via STING.

**Figure. 1.**
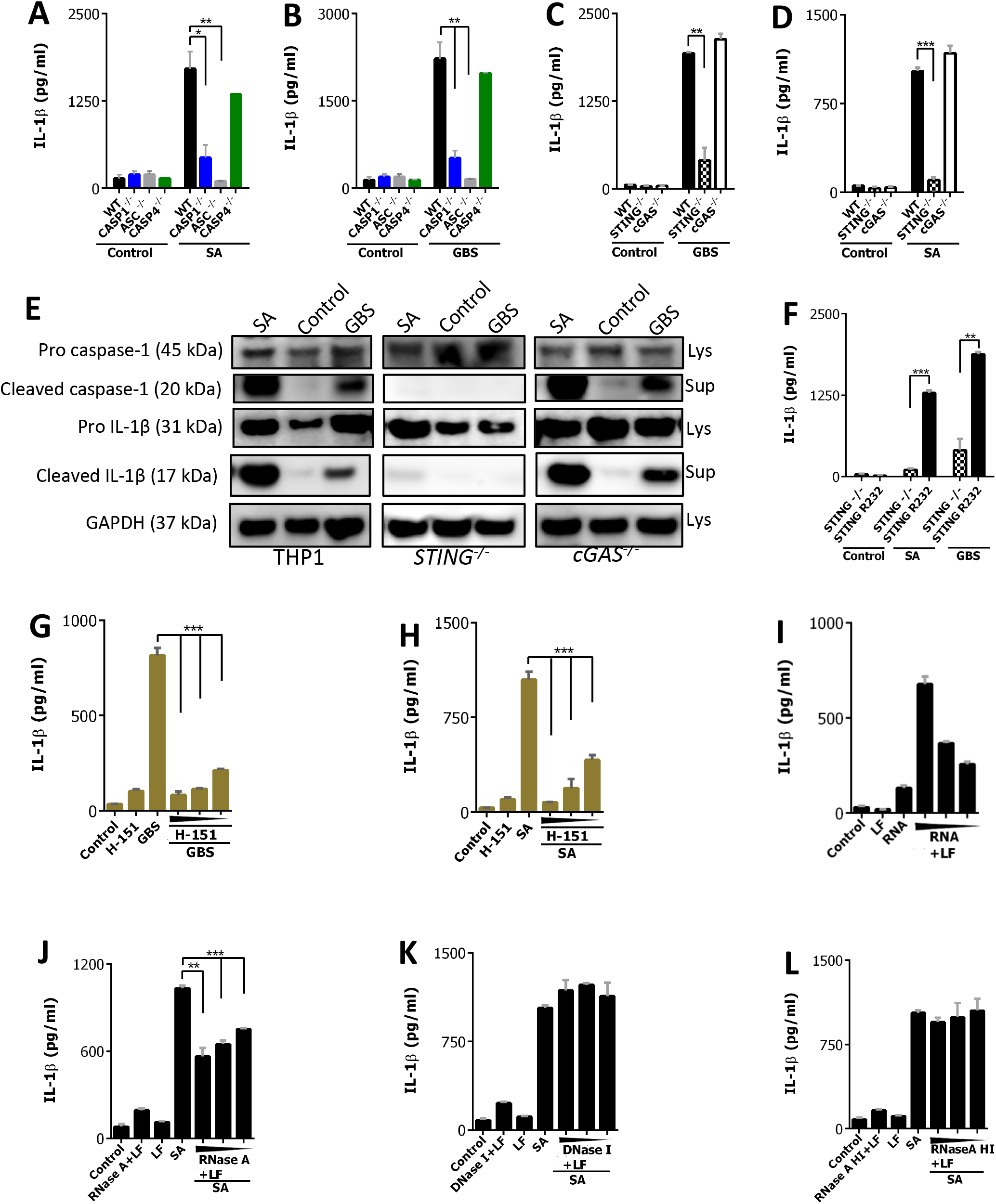
Gram-positive bacteria and bacterial RNA induce the canonical inflammasome pathway via STING. **(A-D)** THP1 (black), *CASP1^-/-^* (blue), *ASC^-/-^* (grey), *CASP4^-/-^* (green), *cGAS^-/-^* (white) and *STING^-/-^* (pattern), MΦs were infected with SA (MOI 10) and GBS (MOI 20), IL-1β production was measured in cell supernetant. **(E)** THP1, *STING^-/-^*, and *cGAS^-/-^* MΦs were infected with SA (MOI 10) and GBS (MOI 20), cleaved IL-1β (P17) and caspase-1 (P20) were detected in cell supernetants (Sup) and pro-IL- 1β, pro-caspase-1 and GAPDH in cell lysate by immunoblotting. **(F)** *STING^-/-^* (pattern), *STING^-/-^* expressing STING^R232^ (Black) MΦs were infected with SA (MOI 10) and GBS (MOI 20), IL-1β production was measured in cell supernetant. **(G,H)** Human blood derived MΦs were pre-treated with increasing concentrations of STING inhibitor (H-151) (100 µM, 50 µM, 10 µM) for 1 h and then infected with SA (MOI 10) and GBS (MOI 20) respectively. IL-1β production was measured in cell supernetant. **(I)** IL-1β production in wild type THP1 MΦs following transfection with increasing concentrations of SA RNA (RNA+LF) (5 µg/ml, 2.5 µg/ml, 0.1 µg/ml). **(J,K,L)** THP1 MΦs were pre-treated with increasing concentrations of cytosolic RNase A (RNase A+LF) (100 ng/ml, 50ng/ml, 10 ng/ml) followed by infection with SA. IL-1β production was measured in cell supernetant. Heat inactivated RNase A (100 ng/ml, 50ng/ml, 10 ng/ml) **(K)** and DNase I (100 ng/ml, 50ng/ml, 10 ng/ml) **(L)** were used as a control in similar experimental setup. Data shown are mean ±SD (n=3), representative of at least three independent experiments. Asterisks indicate statistically significant differences (∗p < 0.05, ∗∗p < 0.01 and ∗∗∗p < 0.001).

### *S. aureus* and GBS secrete multi-lamellar lipid bodies (MLBs) that contain staphyloxanthin type of lipids and RNA

*S. aureus* and GBS mainly follow an extracellular or phagosomal lifestyle. However, activation of STING and NLRP3 by bacterial RNA requires cytoplasmic delivery of PAMPs. Extracellular vesicles (EV) such as outer membrane vesicles from Gram-negative bacteria deliver LPS into the cytosol (34, 41). Heterogeneous EV-like structures have been observed in Gram-positive bacteria; these have been reported as membrane vesicles (MVs) but their formation and ultrastructure remain unknown (42). Hence, we sought to isolate and characterize these membrane vesicles from Gram-positive bacteria and determine their ability to induce the STING-dependent canonical inflammasome. We found that *S. aureus* and GBS showed multiple lobular protrusions 50-100 nm in size through the cell wall when subjected to scanning electron microscopy (SEM) **(Fig. 2A, 2B)**. Occasionally, these structures were seen to form assemblies of several 100 nm sizes, filling the intracellular space **(Fig. 2C)**.

**Figure. 2.**
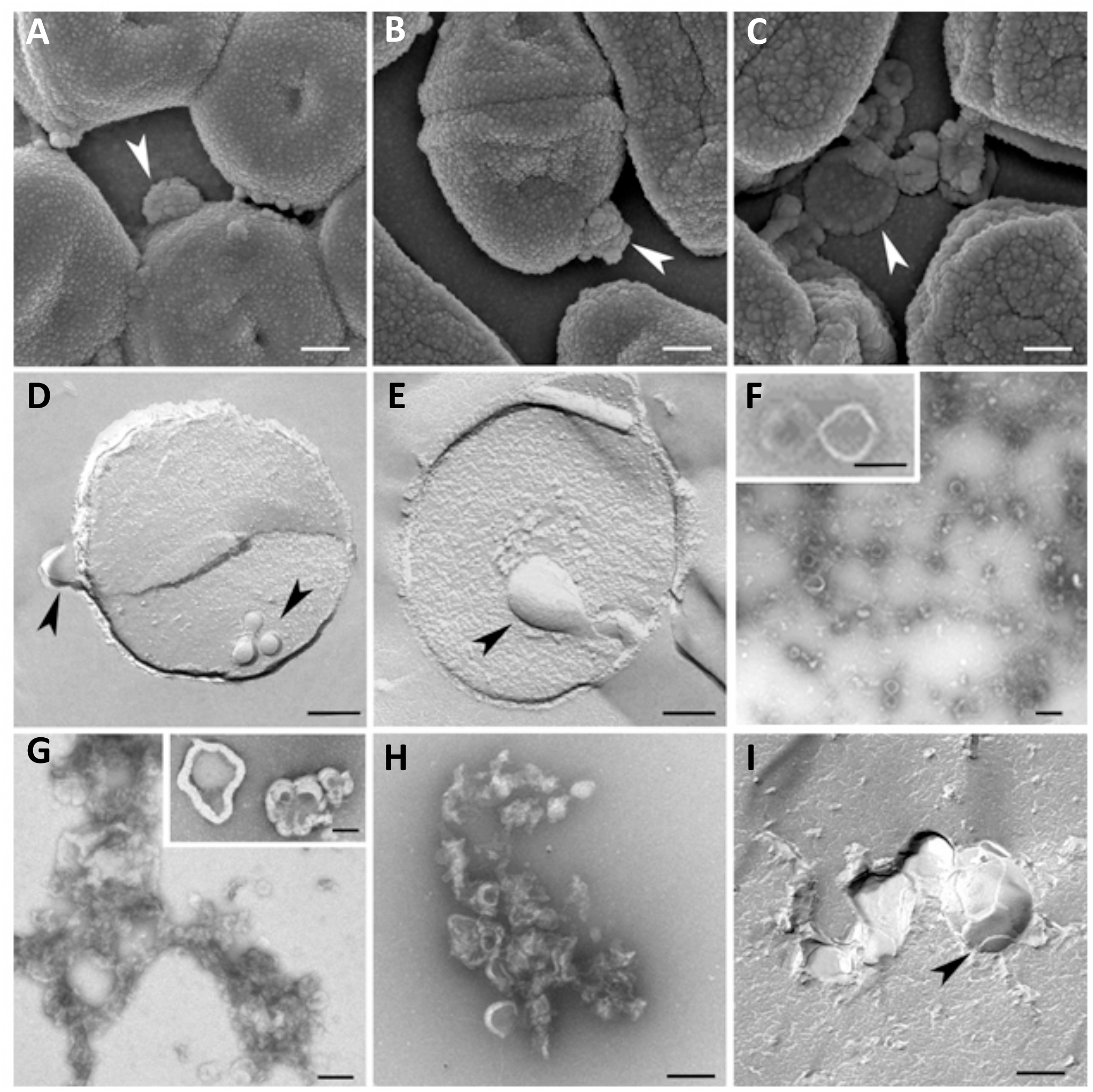
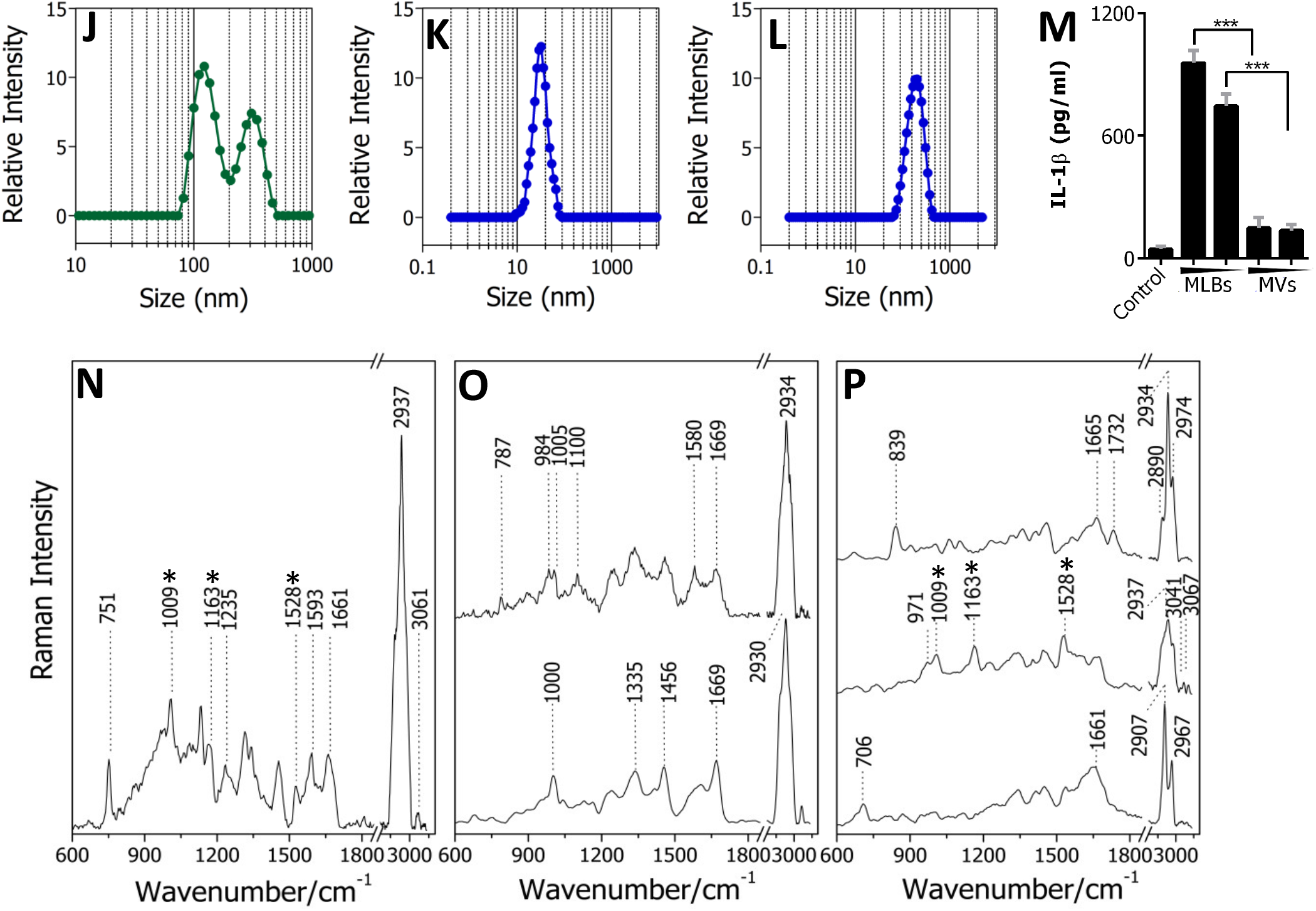
*S. aureus* and GBS secrete multi-lamellar lipid bodies (MLBs) that contain staphyloxanthin type of lipids and RNA. **(A,B)** Scanning electron microscopic (SEM) images of *S. aureus* **(A)** and GBS **(B)**. The arrowhead indicates the multi-lobular secretion from the membrane. **(C)** Magnified SEM image of GBS showing large aggregated assemblies of MLBs. **(D)** Freeze fracture TEM image of *S. aureus*. The arrowhead indicates cytoplasmic accumulation and secretion of lipid bodies. **(E)** Freeze fracture TEM image of GBS; the arrowhead indicates bacterial membrane imbibing the cytoplasmic accumulated lipid bodies. **(F)** Negative staining TEM images of purified membrane vesicles (MVs) from GBS. The inset image shows a magnified ultrastructure of MVs. **(G,H)** Negative staining TEM images of purified Multi lamellar lipid bodies (MLBs) from GBS **(G)** and *S. aureus* **(H)**. The inset image shows a magnified ultrastructure of MLBs. **(I)** Freeze fracture TEM image of GBS culture supernatant purified MLBs. The arrowhead indicates multi lamellar lipid bodies (MLBs). Scale bar in image (a, b c =200 nm, d, e, f, g, h, i and inset = 100 nm). **(J)** Size distribution of vesicles isolated from *S. aureus* (MLBs+MVs) by dynamic light scattering. The x-axis is set to logarithmic scale. **(K,L)** DLS analysis representing size of MVs **(K)** and MLBs **(L)** isolated from *S. aureus*. The x-axis is set to logarithmic scale. **(M)** IL-1β production was measured in wild type THP1 MΦs following stimulation with MLBs (10 µg/ml, 5 µg/ml) and MVs (10 µg/ml, 5 µg/ml) isolated from SA. Data shown are mean ±SD (n=3), representative of at least three independent experiments. Asterisks indicate statistically significant differences (∗∗∗p < 0.001). **(N)** Raman spectra of *S. aureus*. The prominent Raman peaks at 1009, 1163 and 1528 cm^-1^ corresponds to staphyloxanthin type of lipids are marked with dotted lines. **(O,P)** Raman spectra of MLBs isolated from *S. aureus*. The prominent Raman peaks corresponding to staphyloxanthin type of lipids are marked with star. For positive control refer (Figure 6 A,B) Raman spectra of HPLC purified staphyloxanthin type of lipids.

To explore the formation of these vesicles further we subjected *S. aureus* and GBS to freeze-fracture electron microscopy (FEM). Similar to SEM, FEM revealed accumulated membranous structures both inside and outside the bacterial cell wall **(Fig. 2D, 2E).** Negative-staining TEM of vesicles isolated from GBS and *S. aureus* showed multiple membrane vesicle (MV)-like structures of 50 nm size. Upon closer inspection, some of these structures reveled to 100-400 nm size multi-lobular assemblies. We fractionated and purified both these structures and, on electron microscopy, noted 50 nm uni-lamellar MVs which were abundant in lower density fractions **(Fig. 2F)** whereas several 100 nm lipid-rich multi lamellar structures were abundant in the higher density fractions **(Fig. 2G-I)**. This structure and size heterogeneity was in contrast to *E. coli* OMVs that were uni-lamellar, spherical and dispersed **(Supplemental Fig. 2A)**. Similar preparations from *S. aureus* when observed under TEM showed alternatively structured lamellar assemblies been secreted **(Fig. 2H)**. Due to their structural and morphological characteristics we named these Gram-positive secretions ‘multi- lamellar lipid bodies’ (MLBs). FEM electron microscopy analysis noted alternating sheet-like structures **(Fig. 2I)**. Hence, our analysis demonstrates Gram-positive bacteria secrete MLBs in addition to well-characterized MVs.

To further characterize MLBs and MVs secreted by Gram-positive bacteria, we separated MLBs and MVs using density gradient ultracentrifugation followed by quality measurements. We subjected vesicle mixture (MLBs+MVs), purified MVs and MLBs to Dynamic Light Scattering (DLS). Similar to electron microscopy, DLS revealed two partly overlapping populations in the vesicle mixture (MVs+MLBs), one 50-100 nm and the other around 400 nm **(Fig. 2J)**. The size distribution of purified SA MVs was ∼50-100 nm **(Fig. 2K)** and, ∼400 nm for purified MLBs **(Fig. 2L)**. Thus *S. aureus* secretes heterogeneous populations of vesicles that differ in size and density. The immunological relevance of these vesicles was determined upon stimulation of human macrophages with increasing concentrations of MVs and MLBs isolated from *S. aureus*. To our surprise MLBs activated the inflammasome with abundant release of IL-1β **(Fig. 2M)** while MVs failed to activate the inflammasome for IL- 1β release **(Fig. 2M)**. However, MVs and MLBs could both induce TLR-dependent TNF-α production **(Supplemental Fig. 2B)**.

As MLBs and MVs showed different stimulation patterns we investigated the lipid composition of these vesicles. Both contained abundant amounts of lysyl-dipalmitoyl phosphatidylglycerol (lysyl-PG), one of the membrane lipids of *S.* aureus (43). On detailed analysis of the MS fragmentation pattern, diagnostic fragment ion and corresponding intensity showed the abundant species of lysyl PG were in the range of *m/z* 800-900 in MVs, MLBs including the mixture of MLBs and MVs (MLB+MV) of *S. aureus* **(Supplemental Fig. 2C)**.

Staphyloxanthin biosynthetic pathway products play an important role in innate immune activation (21, 22, 23). We thus postulated that MLBs and MVs may differ in their abundance of the staphyloxanthin type of lipids. The prominent lipid identified was in the range of *m/z* 400-500. After detailed analysis of their Shimadzu LCMS data and comparison of the retention time and MS spectra, we could detect the presence of 4,4’-diaponeurosporenoic acid, the precursor of the staphyloxanthin biosynthetic pathway. The relative abundant species of 4,4’-diaponeurosporenoic acid, was detected at *m/z* 432-433 in MLBs, and in a mixture of MVs and MLBs (MV+MLB) and in *S. aureus* bacteria **(Supplemental Fig. 2D)**. However, of note MVs did not indicate the presence of 4,4’-diaponeurosporenoic acid at *m/z* 433. This finding highlights our electron microscopic observation that MLBs are higher order structures formed in the presence of polyunsaturated lipids that appear to arise from the staphyloxanthin biosynthetic pathway **(Fig. 2G-I, Supplemental Fig. 2D).** Moreover, the LC-MS profile also identified small molecules such as c-di-AMP (CDN) and its derivative pApA in *S. aureus* MLBs. The retention time of c-di-AMP and pApA detected in *S. aureus* MLBs was around 2.73 min and 2.75 min respectively when compared to synthetic ligands **(Supplemental Fig. 2E, Supplemental Table 3)**.

We performed Raman spectroscopic analysis of *S. aureus* MLBs to understand its biochemical cargo. MLB Raman spectra were carefully sorted as per dominating biomolecules using vibrational signatures deduced from the literature **(Supplemental Table 1).** Average Raman spectra were generated for carbohydrates, proteins, lipids and nucleic acids **(Fig. 2N-P)**. Nucleic acids and lipids were more prominent than carbohydrates **(Supplemental Table 1)**. To identify the type of lipid present within the MLBs, we analysed spectral assignments and found unsaturated fatty acids and long-chain lipids as seen by Raman vibrations at 2974, 2967, 1732, 1669, 1528 and 1163 cm^-1^. The presence of prominent vibrations at 1009, 1163 and 1528 cm^-1^ indicate microbial polylene-type lipids within the MLBs. The ribose type of nucleic acid vibration was seen by 1580, 1335 and 787 cm^-1^ Raman bands. Detailed Raman assignments are provided in **(Supplemental Table 1)**. Following the prominent detection of lipids by LC-MS, electron microscopy and Raman spectroscopy, we subjected MLB preparations to thermotropic analysis, performing electron microscopy at higher temperatures. Consistent with our previous observations, temperature elevation from 25°C to 37°C produced an aqueous type of MLB texture **(Supplemental Fig. 2F)**. Similar thermodynamic properties of MLBs were seen during Dynamic Light Scattering (DLS) analysis performed at 15°C to 55°C. The reduction in size at higher temperatures suggests thermochemical liquefaction or phase transition of lipids present within the MLBs **(Supplement Fig. 2G)**. The above overall data indicate that Gram-positive bacterial secretions have at least two types of vesicles that differ in density, size and lipid content. Importantly, they differ in their immunological function of inducing the inflammasome in human macrophages and this may bestow a further immune-stimulatory capacity to bacteria during infection.

### MLBs from Gram-positive bacteria activate the STING dependent NLRP3 canonical inflammasome pathway

To dissect the role of MLBs we postulated that MLBs could activate the inflammasome when secreted by bacteria. Similar to THP1 macrophages, release of IL-1β and cell death were seen when human blood-derived primary macrophages were stimulated with GBS and SA MLBs **(Fig. 3A, 3B)**.

**Figure. 3.**
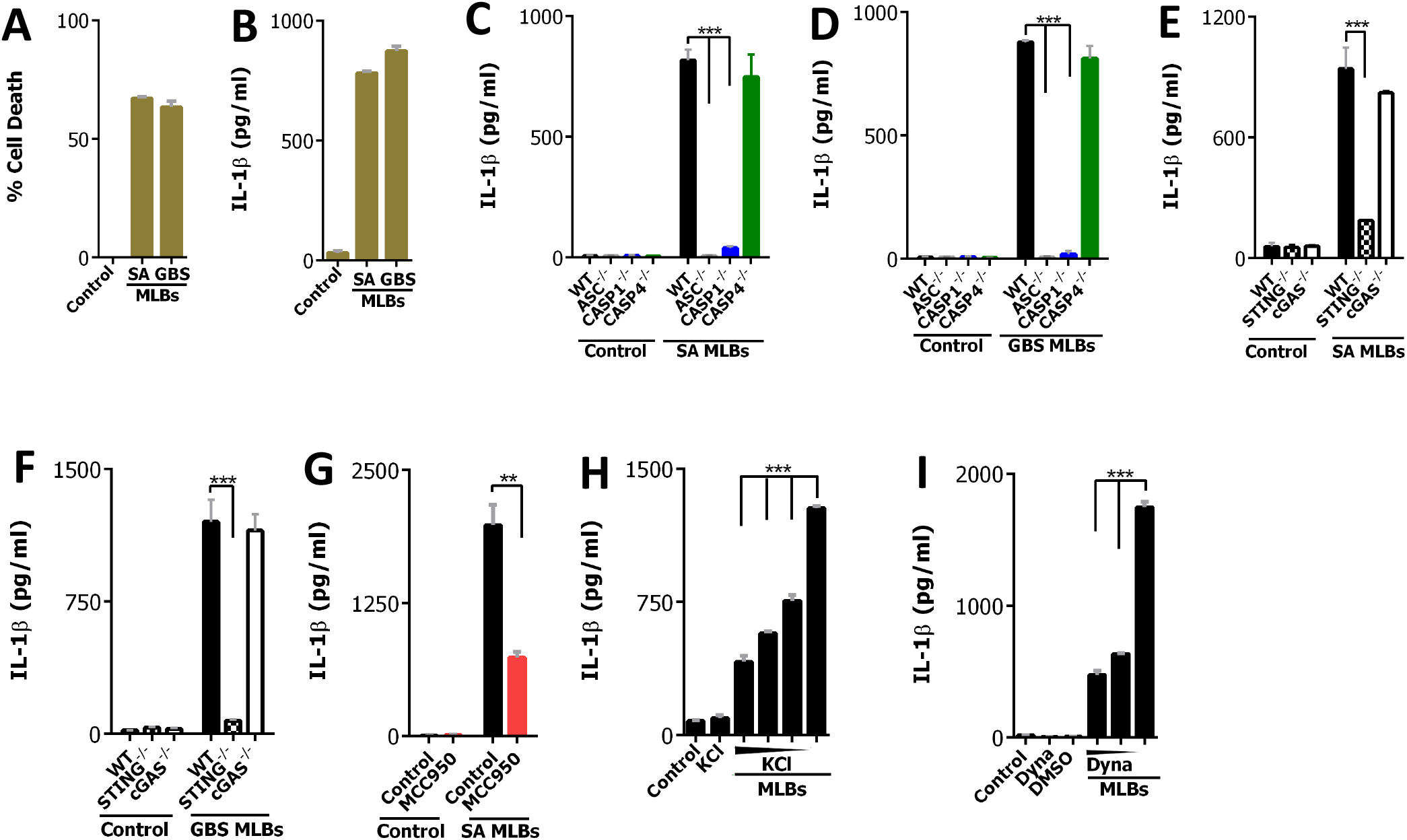
MLBs from Gram-positive bacteria activate the STING dependent NLRP3 canonical inflammasome pathway. **(A,B)** Cell death and IL-1β production measured in human blood-derived MΦs following stimulation with GBS and *S. aureus* MLBs (10 µg/ml) respectively. **(C,D)** THP1 (black), *ASC^-/-^* (grey), *CASP1^-/-^* (blue), *CASP4^-/-^* (green) MΦs show IL-1β production following stimulations with *S. aureus* and GBS MLBs (10 µg/ml) respectively. **(E,F)** THP1 (black), *STING^-/-^* (pattern) *cGAS^-/-^* (white) MΦs show IL-1β production when stimulated with *S. aureus* and GBS MLBs (10 µg/ml) respectively. **(G)** Wild type THP1 MΦs (black) were pre-treated with and without MCC950 inhibitor (red) (5 µM) for 1 h and stimulated with SA MLBs; IL-1β was measured in the supernatant. **(H)** Wild type THP1 MΦs (black) were pre-treated with increasing concentrations of KCl (60 mM, 45 mM, 75 mM) and stimulated with GBS MLBs (10 µg/ml). IL-1β production was measured in cell supernatant. **(I)** Wild type THP1 MΦs were pre-treated with increasing concentrations of Dynasore inhibitor for 1 h and stimulated with GBS MLBs. IL-1β production was measured in cell supernatant. Data shown are mean ±SD (n=3), representative of at least three independent experiments. Asterisks indicate statistically significant differences (∗p < 0.05, ∗∗p < 0.01 and ∗∗∗p < 0.001).

We purified MLBs from Gram-positive bacteria (SA and GBS) and stimulated macrophages replete or deficient in caspase-1 or caspase-4 to examine canonical inflammasome activation. Similar to bacterial infection, IL-1β responses were reduced in caspase-1 deficient cells whereas responses remained unchanged in caspase-4 deficient cells. This is consistent with the notion that Gram-positive bacteria lack LPS and hence are caspase-4 independent. Similarly, IL-1β responses were completely abrogated in adaptor ASC-deficient cells when stimulated with MLBs **(Fig. 3C, 3D)**. Similar to studies with bacterial infections, MLBs isolated from Gram-positive bacteria induced release of IL-1β that was STING-dependent but cGAS-independent **(Fig. 3E, 3F)**. MLBs also showed NLRP3 dependency when tested with the potent NLRP3 inhibitor MCC950 (44) **(Fig. 3G)**. As most activators of NLRP3 are K^+^ efflux-dependent, we determined the significance of this efflux on MLB-mediated NLRP3 activation. Indeed, blocking K^+^ efflux prevented inflammasome activation when stimulated with MLBs, and when using nigericin as a positive control **(Fig. 3H, Supplemental Fig. 3A, 3B).** As a control to Gram-positive MLBs, Gram-negative outer membrane vesicles (OMVs) are taken up by cells via an endocytic pathway with LPS escaping the early endocytic compartments to activate caspase-11 (32) **(Supplemental Fig. 3C)**. Consistent with these findings, MLBs are taken up by cells via dynasore-sensitive dynamin-dependent endocytic pathways **(Fig. 3I)**.

### RNA aptamers present in bacterial small non-coding RNA activates STING to induce the IL-1β response

Cytoplasmic bacterial nucleic acids activate the caspase-1 dependent canonical pathway (20, 30). As our MLB analysis showed entrapped RNA, with the use of lipofectamine (LF) we delivered purified MLB RNA alone to cells either replete or deplete of caspase-4 or, caspase- 1 and subsequently measured IL-1β and cell death responses. Cytoplasmic RNA (RNA+LF) showed a caspase-1 dependency for LDH response as compared to wild-type and caspase-4 deficient cells. Consistent responses were found for IL-1β **(Fig. 4A, 4B)**. NLRP3 was also required for cytosolic sensing of MLB RNA, since RNA-induced IL-1β secretion was profoundly reduced when inhibited with MCC950 **(Fig. 4C)**. When MLB RNA was combined with LPS and delivered to the cytoplasm, cell death and IL-1β responses became both caspase-4 and caspase-1 dependent **(Supplemental Fig. 1A, 1B)**. This set-up underlines the robustness of RNA as a PAMP for activation of the canonical inflammasome pathway. Cytokine production can be abrogated when enzymatically treated with RNase A (20). Experimentally translating cleavage of RNA ligands by RNase A in MLB-stimulated cells resulted in >80% reduction in cytokine production **(Fig. 4D).** At the same time cells treated with heat inactivated RNase A and DNase I and showed no difference in the release of IL-1β **(Fig. 4E, 4F)**. Similar observations were found for SA infections **(Fig. 1J, 1K, 1I)**. Taken together, these findings indicate that Gram-positive bacteria and MLBs activate the canonical NLRP3 pathway via STING without activating cGAS using bacterial RNA.

**Figure. 4.**
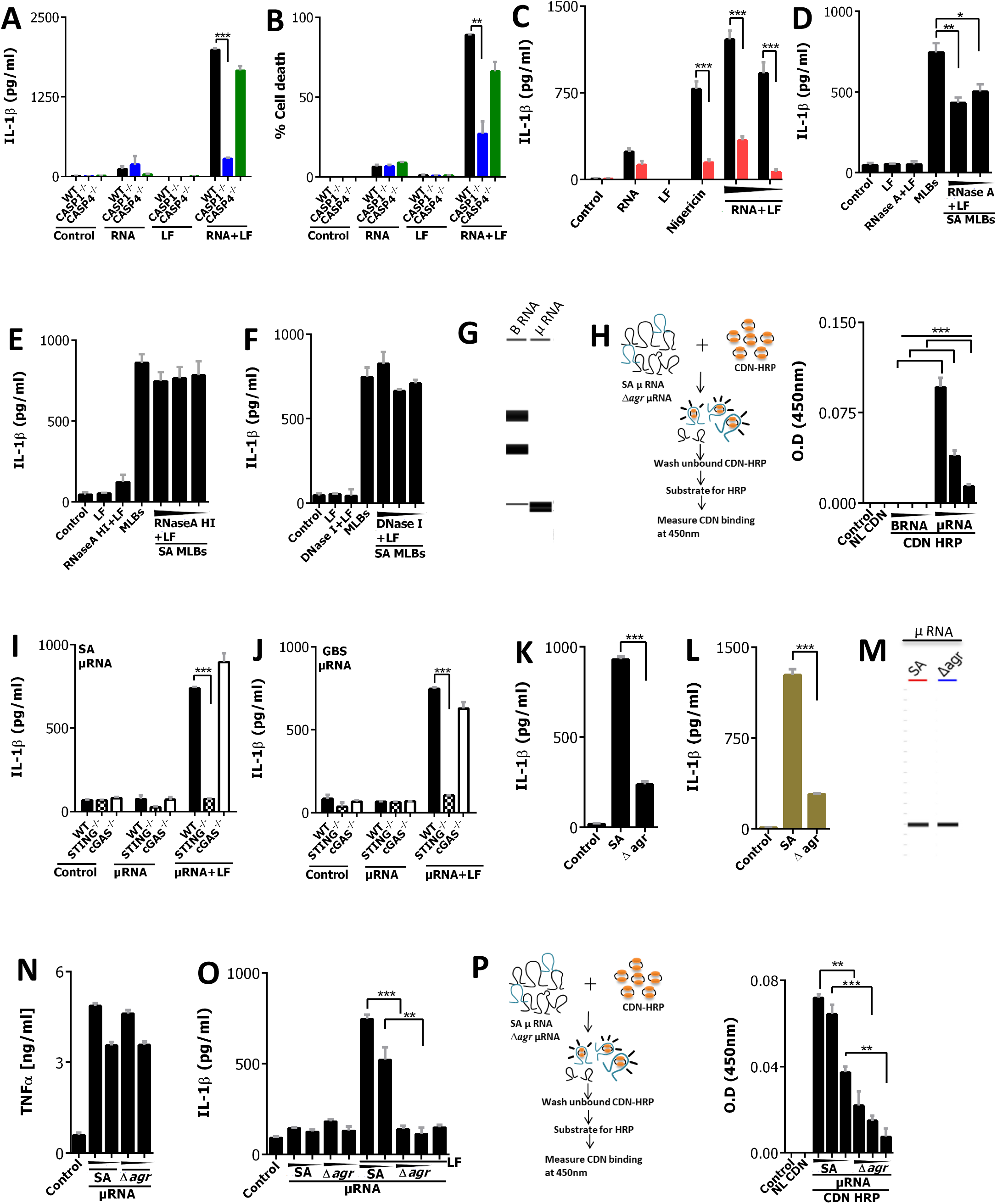
RNA aptamers present in bacterial small non-coding RNA activates STING for IL-1β response. **(A,B)** THP1 (black), *CASP1^-/-^* (blue), *CASP4^-/-^* (green) MΦs were stimulated with MLB RNA, cytosolic MLB RNA (RNA+LF) (5 µg/ml). IL-1β and cell death were measured in supernetant. **(C)** THP1 (black) MΦs were pre-treated with increasing concentrations of MCC950 inhibitor (red) (5 µM, 10 µM) and stimulated with *S. aureus* RNA, lipofectamine, nigericin (6.7 µM), cytoplasmic RNA (RNA+LF) (5 µg/ml, 2.5 µg/ml), IL-1β was determined in cell supernetant. **(D,E,F)** THP1 MΦs were pre-treated with increasing concentrations of cytosolic RNase A (RNase A+LF) (100 ng/ml, 50ng/ml, 10 ng/ml) followed by stimulation with SA MLBs, IL- 1β was determined. Heat inactivated RNase A (100 ng/ml, 50ng/ml, 10 ng/ml) **(E)** and DNase I (100 ng/ml, 50ng/ml, 10 ng/ml) **(F)** were used as a control in similar experimental setup. **(G)** RNA capillary gel electrophoresis of two types of purified GBS RNA (Big-BRNA (>200 nucleotides), micro-µRNA (<200 nucleotides)) (1 µg) run on QIAxcel advanced system. Please note lower most band in B RNA sample is a capillary electrophoresis marker. **(H)** Optical density at 450 nm measured in binding assay of GBS µRNA, B RNA (B RNA) (4 µg, 2 µg, 1 µg) with HRP-labelled CDN (c-di-AMP) (16 µM). Non-labelled CDN (NL c-di- AMP) was used as a control. **(I,J)** THP1 (black), *STING^-/-^* (pattern) and *cGAS^-/-^* (white) MΦs were stimulated with GBS **(I)** and SA **(J)** µRNA (µRNA), cytosolic µRNA (µRNA+LF) (5 µg/ml) and IL-1β production measured. **(K)** IL-1β production was measured in THP1 MΦs upon infection with *S. aureus* (MOI 10) and Δ*agr* mutant (MOI 10) bacteria. **(L)** IL-1β production measured in human blood-derived MΦs when infected with *S. aureus* (MOI 10) and Δ*agr* mutant (MOI 10) bacteria. **(M)** RNA capillary electrophoresis gel of µRNA (1 µg) from *S. aureus* and Δ*agr* mutant bacteria run on QIAxcel advanced system. **(N)** TNF-α production measured in THP1 MΦs when stimulated with increasing concentration of *S. aureus* and Δ*agr* mutant µRNA (5 µg/ml, 2.5 µg/ml). **(O)** IL-1β production measured in THP1 (black) MΦs when stimulated with µRNA (5 µg/ml, 2.5 µg/ml) from *S. aureus* and Δ*agr* mutant on the surface (SA and Δ*agr)* and into the cytosol (SA+LF, Δ*agr*+LF) and only LF. **(P)** Optical density at 450 nm measured in binding assay of bacterial µRNA (4 µg, 2 µg, 1 µg) from *S. aureus* and Δ*agr* mutant with HRP labelled CDN (c-di-AMP) (16 µM). Non-labelled CDN (NL c-di-AMP) was used as a control. Data shown are mean ±SD (n=3), representative of at least three independent experiments. Asterisks indicate statistically significant differences (∗p < 0.05, ∗∗p < 0.01 and ∗∗∗p < 0.001).

Although no link is known between RNA and STING (45), we postulated that bacteria deliver RNA aptamers to the cytosol to sequester CDN ligands to STING for downstream activation of IL-1β. As several types of RNA (30) activate the inflammasome pathway, we investigated which RNA packed within the MLBs could activate the canonical inflammasome pathway via STING. To this end we isolated and purified different sizes of RNA from GBS, e.g. big RNA that contains >200 nucleotides and small non-coding µRNA that contains 15-200 nucleotides. Analysis by Qiagen QIAxcel capillary gel electrophoresis confirmed the size and purity of the RNA species **(Fig. 4G)**.

STING is a well-studied receptor for CDN (6, 15, 16). Hence, small RNA aptamers may play a role in sequestering CDN to STING by a yet to be discovered mechanism. One such possibility arises through small RNA aptamers which may bind to small molecule metabolites such as CDN. To evaluate this possibility we tested GBS µRNA and big RNA for CDN-binding capacity in an assay using horse radish peroxidase labelled c-di-AMP as depicted in **Fig. 4H**. Consistent with our hypothesis, we found bacterial µRNA fraction carries CDN-binding capacity as a significant quantity of µRNA was bound to CDN whereas big RNA failed to bind **(Fig. 4H)**. To further dissect the role of µRNA, we activated the NLRP3 inflammasome by delivering purified µRNA species to genetically STING, cGAS-deficient or replete THP1 macrophages. To our surprise, we found the inflammasome was activated in a STING-dependent, cGAS-independent manner for µRNA species (15-200 nucleotides) from GBS and SA **(Fig. 4I, 4J)**, resembling the activation pattern of bacterial infections and MLBs. Non-cytoplasmic activation by all RNA types resulted in no release of IL-1β **(Fig. 4I, 4J)**. Consistent with these results, µRNA isolated from GBS MLBs also activated the inflammasome in a STING-dependent, cGAS-independent manner **(Supplemental Fig. 4A)**.

The above results indicate that Gram-positive bacteria may express specialized µRNA aptamers with a strong binding capacity to CDN. CDN are known signal transducers of quorum-sensing mechanisms in bacteria (16, 25) and production of CDNs is vital for the bacterial life cycle (46). In SA a well-studied quorum sensing system is regulated by the accessory gene regulator (agr) locus. The AGR quorum sensing system plays a central role in virulence regulation and pathogenesis. Mutation in the AGR locus shows no growth defects but leads to loss of various virulence-related genes causing a pleiotropic effect on bacterial pathogenicity (47, 25). A majority of clinical isolates produce high levels of transcripts dependent on the AGR locus. Hence, we postulated that expression of CDN aptamers may directly or indirectly be governed by the AGR locus. To evaluate this possibility further, we tested SA and isogenic mutant Δ*agr* bacteria for the inflammasome activation pathway. Consistent with our hypothesis, the IL-1β response was abrogated when infected with Δ*agr* in THP1 macrophages and human blood derived macrophages as compared to SA **(Fig. 4K, 4L)**. These results were consistent with the previous literature, where it shows that *S. aureus* associated pore forming toxins enhance the process of inflammasome activation. We postulated the abrogated IL-1β response in Δ*agr* infection was due to a lack of CDN aptamer expression in mutant bacteria. To further test this hypothesis we isolated µRNA from Δ*agr* and SA and found equal production of µRNA when analysed on Qiaexcel capillary gel electrophoresis **(Fig. 4M)**. Bacterial RNA, when given to the cell surface, induced TLR-dependent TNF-α response. When Δ*agr and* SA isolated µRNA were tested for TLR-dependent TNF-α production, Δ*agr* -µRNA showed an equal and comparable TNF-α response to SA µRNA **(Fig. 4N)**. However, when the same µRNA was given to the cytosol, Δ*agr* µRNA showed a pronounced reduction in IL-1β production compared to SA µRNA **(Fig. 4O)**. This further indicates that Δ*agr* µRNA may lack CDN-binding aptamers and consequently lacks an ability to stimulate IL-1β production. To evaluate this hypothesis further, we tested µRNA from SA and Δ*agr* for CDN-binding capacity in an assay using horse radish peroxidase-labelled c-di-AMP as shown in **Fig. 4P.** µRNA from SA showed more CDN-binding capacity than µRNA from mutant Δ*agr* bacteria **(Fig. 4P)**. To independently confirm the comparative binding capacity of SA and Δ*agr* µRNA to CDN, we subjected CDN-bound RNA fractions to LC-MS analysis. The increasing concentration of SA µRNA purified upon binding to CDN showed the prominent presence of bound CDN at aretention time of 3.31 min **(Supplemental Fig. 4B (II)).** However, at the same time Δ*agr* µRNA fractions showed no detectable bound CDN **(Supplemental Fig. 4B (III))**. Of note the SA and Δ*agr* µRNA used in this analysis did not show any residual CDN **(Supplemental Fig. 4B (I)).** Collectively, SA µRNA but not the Δ*agr* µRNA have the CDN binding capacity and hence can be concluded that Δ*agr* µRNA lack CDN-binding aptamers. Moreover, owing to its lack of CDN-binding aptamers, after external addition of CDN to Δ*agr*, µRNA did not recover its IL-1β producing ability **(Supplemental Fig. 4C).** All the above results indicate an essential role of the CDN-binding aptamer in IL-1β production, and for the AGR locus of *S. aureus* for expression of CDN-binding aptamers.

### Differentially abundant µRNA species are present in *S. aureus* MLBs

Microbial extracellular vesicles contain small RNA (48), which may regulate the host immune system upon cytoplasmic delivery (48, 49). Our analysis raises the possibility that CDN-binding RNA aptamers are enriched in MLBs which, in turn, activates STING within the host cytosol. Hence, to characterize RNA species from MLBs, µRNA isolated from MLBs (SA- MLB) underwent deep RNA sequencing and were compared to *S. aureus* (SA) and Δ*agr S. aureus* (SA-Δ*agr*) µRNA. For sequence analysis, obtained RNA seq reads were mapped on *S. aureus* reference genome **(Supplemental Result SR2).** We found 186 genomic regions showing differentially abundant RNA seq reads on the *S. aureus* N315 genome **(Fig. 5A)**. Similar results were obtained for *S. aureus* NCTC 8325 (data not shown). 85, 8 and 17 genomic regions were unique to SA, SA-Δ*agr* and SA-MLB respectively. However, 41, 8 and 5 genomic regions were common to SA, SA-Δ*agr*, and SA-MLB, respectively. 22 genomic regions were abundantly present in all three and were mainly comprised of 6S RNA, tmRNA and 4.5S RNA and reads arriving from a noncoding pseudo gene **(Supplemental Fig. 5A)**. One such read (named as HP) was commonly abundant in all three libraries and was used as a control in our further analysis. Collectively, our NGS analysis raises the possibility of a unique set of µRNA species enriched within SA-MLB **(Fig. 5A, Supplemental Fig. 5A)**. Further, scrutinization revealed several differentially abundant genomic locations covering known sRNA from *S. aureus* **(Supplemental Fig. 5A, 5B, Supplemental Result SR2)**. Interestingly, reads for Rsa C, Sbr C, RNAIII, and WAN01CC66-rc were abundantly present in SA and SA-MLB and absent in SA-Δ*agr* **(Supplemental Fig. 5B)**. All other sRNAs were differentially abundant in reads of SA, SA-MLB and SA-Δ*agr* **(Supplemental Fig. 5B, 5C, Supplemental Table 4).** Interestingly, our analysis demonstrate that SA MLBs contain distinct parts of sRNA species that are differentially enriched from the originating bacteria **(Supplemental Results SR2, Supplemental Fig. 5D)** indicating the possibility of specific processing and packaging of sRNA species within MLBs of *S. aureus*.

**Fig. 5.**
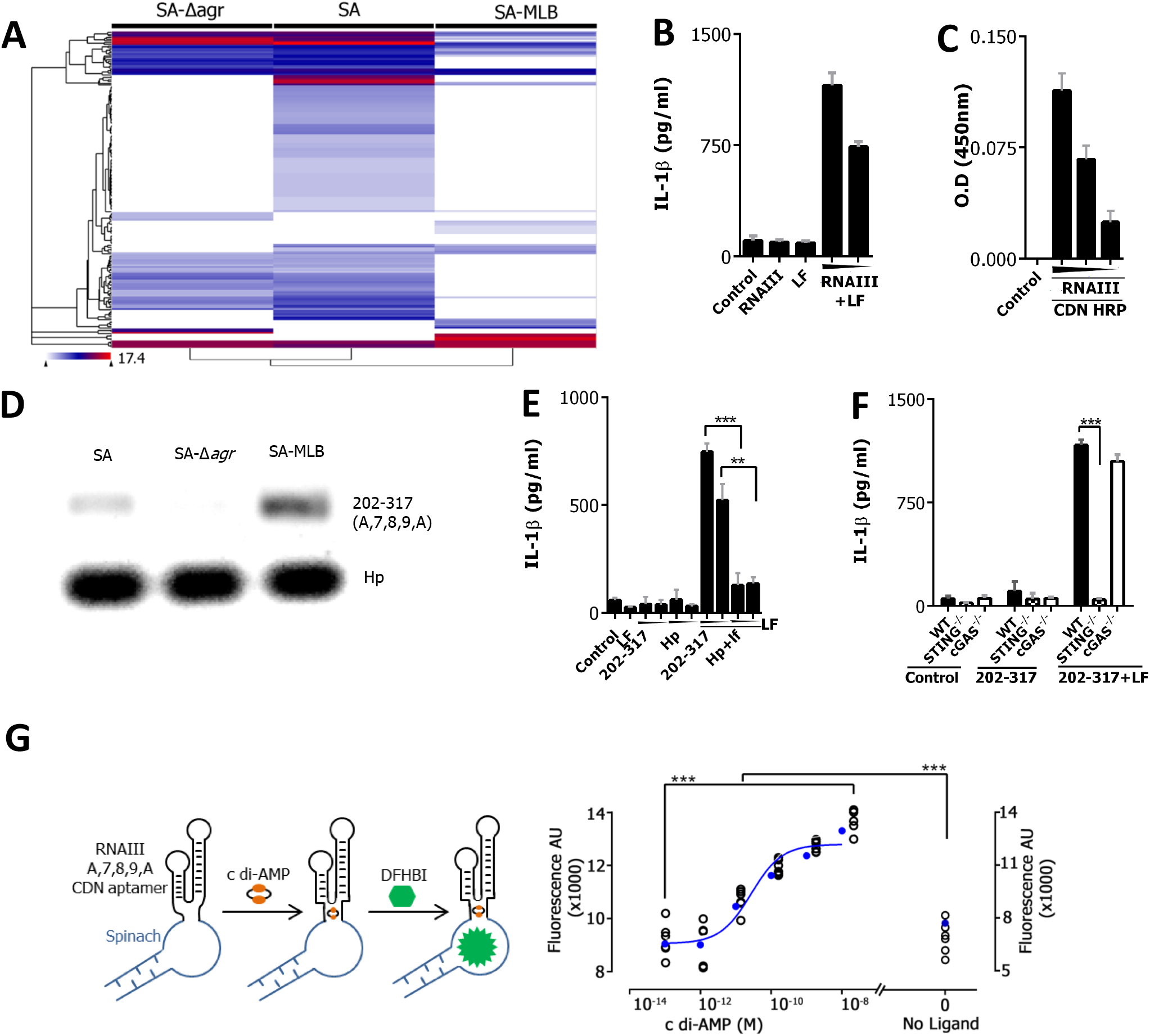
Differentially abundant µRNA species are present in *S. aureus* MLBs and analysis of RNAIII with A,7,8,9,A stem loop domain present in MLBs for the inflammasome activation. **(A)** Heat map of differentially abundant µRNA sequence reads obtained through deep sequencing in SA, SA-MLB and SA-Δ*agr*. The reads were mapped on *S. aureus* subsp. *aureus* N315 (accession number: NC_002745.2) corresponding to 186 differentially abundant genomic regions. **(B)** IL-1β production measured in THP1 (black) MΦ following stimulations with *in-vitro*- transcribed cytosolic RNAIII (RNAIII+LF) (5 µg/ml, 2.5 µg/ml), surface RNAIII and LF. **(C)** Optical density at 450 nm measured in binding assay of *in-vitro*-transcribed RNAIII (4 µg, 2 µg, 1 µg) with cyclic-di-AMP (16 µM). **(D)** Semi quantitative PCR analysis of RNAIII central domain A,7,8,9,A (202-317 nt) in SA, SA-Δ*agr* and SA-MLB. HP RNA was used as a control. **(E)** IL-1β production measured in THP1 (black) MΦ following stimulations with *in-vitro*- transcribed RNAIII central domain A,7,8,9,A (202-317 nt) (5 µg/ml, 2.5 µg/ml) on surface and into the cytosol (202-317+LF), HP RNA (HP+LF) and LF. **(F)** THP1 (black), *STING^-/ -^*(pattern) and *cGAS^-/ -^*(white) MΦ were stimulated with *in-vitro*- transcribed RNAIII central domain A,7,8,9,A (202-317 nt) (5 µg/ml, 2.5 µg/ml) on surface and into the cytosol (202-317+LF), and IL-1β production measured. **(G)** Analysis of *in vitro-*transcribed RNAIII central domain A,7,8,9,A (202-317 nt) spinach binding to cyclic-di-AMP at 37°C in presence of DFHBI (10 µM) and different concentrations of cyclic-di-AMP (0.1 pM-10 nM). Data from replicates (black) and mean (blue) are shown. Background fluorescence was subtracted from all data points. Data shown are mean ±SD (n=3), representative of at least three independent experiments. Asterisks indicate statistically significant differences (∗p < 0.05, ∗∗p < 0.01 and ∗∗∗p < 0.001).

### A specific fragment of RNAIII with A,7,8,9,A stem loop domain present in *S. aureus* MLBs activate the inflammasome pathway

RNAIII is a known effector molecule of the quorum sensing system and controls expression of various virulence genes in *S. aureus* (56, 57). We further characterized the key regulatory sRNA, RNAIII from *S. aureus*. Reads of SA, SA-MLBs and SA-Δ*agr* µRNA showed that RNAIII was abundantly mapped in SA and SA-MLBs, however were SA-Δ*agr* dependent. For validation of the role of RNAIII in activation of the inflammasome, we *in-vitro*- transcribed RNAIII and delivered it in increasing concentrations cytosolically to THP1 macrophages. Surprisingly, RNAIII activated the inflammasome for the release of IL-1β **(Fig. 5B)**. Surface stimulations of RNAIII and LF did not induce the NLRP3 inflammasome pathway **(Fig. 5B)**. To access this further, we checked the CDN binding capacity of RNAIII in an assay using horse radish peroxidase labelled c-di-AMP. Similar to SA µRNA, RNAIII could bind to c-di-AMP **(Fig. 5C)** Additionally, mass spectrometry based binding assay further confirmed binding of c-di-AMP to RNAIII **(Supplemental Fig. 5E),** indicating the possibility that RNAIII may contain a CDN-binding aptamer.

RNAIII has a complex secondary structure of 514 nucleotides and comprises 14 hairpin domains (58, 59). Three long range interactions present on the RNAIII brings the 3‘and 5’ends in close proximity. The central domain contains the branched structure of stem loop 7, 8 and 9 which is enclosed by helix A (56, 57). To further analyse the RNAIII in detail, we mapped the RNA seq reads of SA and SA-MLBs on *S. aureus* RNAIII 514 bp genomic locus **(Supplemental Fig. 5F)**. Our analysis revealed that hairpins 1-6 and 10-14 were present in SA (blue) and SA-MLBs (pink) µRNA **(Supplemental Fig 5F)**. Some reads present in the 1- 6 and 10-14 hairpin regions were overlapping in SA and SA-MLB **(Supplemental Fig. 5F)**. The SA-MLB reads showed the unique presence of the central domain A,7,8,9,A hairpins and some stretches in the 3’regions corresponding to 1-380 nt, and 5’regions corresponding to 202-514 nt. The overlaid reads also confirmed that the central domain A,7,8,9,A corresponding to 202-317 nt is exclusively present within SA-MLBs **(Supplemental Fig. 5F)**. We amplified the central domain A,7,8,9,A from an independently prepared cDNA of the SA, SA-Δ*agr* and SA-MLB µRNA. Comparable to the sequencing results, the central domain (A,7,8,9,A) was present in SA-MLBs, absent in SA-Δ*agr*, and slightly present in SA **(Fig. 5D)**. HP, the sRNA expressed in all three µRNA libraries was used as a control **(Fig. 5D)**.

Following this, we *in-vitro*-transcribed 3’and 5’regions corresponding to 1-380 nt and 202- 317 nt, respectively with the overlapping central domain of RNAIII according to the reads within SA-MLB. These *in-vitro*-transcribed RNA when given cytosolically to THP1 macrophages could induce the inflammasome with release of IL-1β **(Supplemental Fig. 5G)**. As the central domain of A,7,8,9,A hairpins corresponding to 202-317 nt were specifically present in SA-MLBs we *in-vitro*-transcribed it with control sRNA HP and stimulated cytosolically on THP1 macrophages. Interestingly, the central domain (202-317 nt) robustly activated the inflammasome leading to IL-1β release **(Fig. 5E)** similar to RNAIII (1-514 nt) **(Fig. 5B),** the control sRNA HP however did not induce IL-1β production **(Fig. 5E)**. Moreover, resembling the activation pattern of bacterial µRNA, central domain (202-317 nt) could release IL-1β in a cGAS-independent, STING-dependent manner **(Fig. 5F)**. This indicates that the A,7,8,9,A hairpins of RNAIII present within SA-MLBs are immuno stimulatory. To characterize RNAIII further we *in silico* folded RNAIII and different parts of RNAIII present within the SA-MLBs using RNA fold web server-Vienna RNA Package (60). The full RNAIII structure (1-514 nt) resembled the predicted structure of 14 hairpins. The central domain of A,7,8,9,A loops structurally resembles known CDN binding aptamers (61). A similar central domain structure was seen in different parts of the RNAIII, such as 1-380, 202-514 and 202-317. This suggest that A,7,8,9,A hairpins of the central domain may be important for RNAIII to activate the inflammasome. Complimenting our mapped RNA seq reads on RNAIII **(Supplemental Fig. 5F)**, we *in silico* simulated mutations around nucleotides 208-217 corresponding to RNAIII stem A. Specific mutations at 212, 213, and 214 were vulnerable for the formation of the central domain. Nucleotides from 1-213 and 213-514 of RNAIII failed to assemble a similar secondary structure to the A,7,8,9,A central domain **(Supplemental Fig. 5H)**. Hence, we *in-vitro*-transcribed RNAIII from 1-213 and 213-514 nt which failed to show the formation of the central domain. The *in-vitro-*transcribed 1-213 and 213-514 nt RNA when given cytosolically failed to induce release of IL-1β in caontrast to complete (1–514) RNAIII **(Supplemental Fig. 5I)**. These results suggests that the formation of the branched central domain of 202-317 nucleotides with hairpin A,7,8,9,A is crucial for CDN binding and release of IL-1β. Moreover this above mutational analysis raises the possibility that the structure of the RNAIII which can bind to CDN is more important than the sequence. To this end we used spinach aptamer with conditionally fluorescent molecule difluoro-4-hydroxybenzylidene imidazolinone (DFHBI) (62) as depicted in **Fig. 5G** to independently confirm the structure specific CDN binding capacity of hairpin A,7,8,9,A of RNAIII. The fluorescence of spinach aptamer depends on the formation of a second stem loop (62). Hence, we replaced the second stem loop with the aptamer domain A,7,8,9,A of RNAIII. Consistent with previous results, the A,7,8,9,A central domain linked to spinach aptamer exhibited strong fluorescence activation in a concentration-dependent manner from 0.1 to 10 nM c-di-AMP **(Fig. 5G)**.

To consolidate our findings of a dependency on STING, we performed a STING dimerization assay following activation by RNAIII, SA µRNA or Δ*agr* µRNA. Consistent with inflammasome activation, Δ*agr* µRNA showed reduced STING dimerization compared to SA µRNA and RNAIII **(Supplemental Fig. 5J)**. Blocking K^+^ efflux reduced the induction of NLRP3 activation **(Fig. 3H)**. Hence, we investigated the trafficking of STING to the lysosome (LAMP-1+ structures) as an indicator of lysosome-dependent NLRP3 activation following STING dimerization. SA µRNA was transfected into human macrophages and stained with anti-LAMP-1 and anti-STING antibody to determine lysosomal trafficking of STING. Upon activation with µRNA, STING concentrated in small vesicle-like punctate structures compared to a dispersed pattern in control preparations **(Supplemental Fig. 5K(I))**. LAMP-1 co-localization with STING was pronounced in these punctate STING structures upon stimulation with SA µRNA as compared with the non-stimulated control **(Supplemental Fig. 5K(I))**. As shown previously, due to a lack of CDN aptamer the IL-1β responses were abrogated by Δ*agr* µRNA. Hence we postulated and confirmed that Δ*agr* µRNA was also abrogated in co-localizing STING to the lysosome compared to SA µRNA **(Supplemental Fig. 5K(I))**. Consistent with these results GBS and SA bacterial infections also led to higher co-localization of STING within lysosomes **(Supplemental Fig. 5K(II))**. These data indicate that CDN binding RNA aptamers present in the µRNA fraction can sequester CDN ligands to STING, thus acting as potent inducers of the inflammasome during Gram-positive bacterial infection **(Supplemental Fig. 5L)**.

### Staphyloxanthin type of lipids in MLBs can target RNA PAMPs to the cytoplasm for activation of the inflammasome

Taking advantage of the robust activation of STING by µRNA we investigated how MLB RNA reaches the cytoplasm to serve as a PAMP during inflammasome activation. In *S. aureus*, multiple pore forming toxins and hemolysins have shown to evolve NLRP3 inflammasome activation. Especially α-hemolysin is shown to release IL-1β in caspase-1 dependent manner in both human and mouse monocytic cells (8). Similarly, GBS pigment/ lipid toxin was also shown to induce membrane permeabilization which causes the efflux of intracellular potassium activating the NLRP3 inflammasome via caspase-1 (24). Hemolytic strains of GBS have been reported to induce NLRP3 inflammasome for the release of IL-1β in murine dendritic cells and macrophages (38, 20). Using Raman spectroscopy, electron microscopy and LC-MS analysis we identified the presence of staphyloxanthin biosynthetic pathway precursors and derivatives within MLBs. We then queried whether these polyunsaturated lipids from MLB had the capacity to deliver RNA aptamers for cytoplasmic inflammasome activation.

Polyene biosynthesis pathway is a major contributor to the synthesis of staphyloxanthin and granadaene by *S. aureus* and GBS, respectively (22, 23, 63). Both these pathways contain many structurally similar polyunsaturated lipids. Granadaene has been implicated in inflammasome activation by GBS (64) though precise mechanisms are unknown. Accordingly, we postulated that similar products of the staphyloxanthin and granadaene biosynthetic pathways, such as 4,4’-diaponeurosporenoic acid, are present in their respective MLBs and can transfer MLB-embedded RNA PAMPs to the cytoplasm.

To this end we HPLC-purified staphyloxanthin (St2) and its pathway precursor 4,4’- diaponeurosporenoic acid (St1) from *S. aureus* **(Supplemental Fig. 6,7)**. The presence of St1 and St2 in HPLC fractions was confirmed by analysis of UV-Vis, MS and NMR spectra **(Supplemental Result 1, Supplemental Fig. 7)**. Having purified these compounds, we examined their Raman spectra and found bacterial carotenoid-specific strong vibrations at 975, 1014, 1168, 1210, 1294, 1451, 1528 and 1581 cm^1^ **(Fig. 6A, 6B, Supplemental Table 2)**. These Raman vibrations were conserved in MLBs isolated from *S. aureus* and GBS and were also present in Raman spectra of complete *S. aureus* and GBS bacteria **(Fig. 2N-P, Supplemental Table 1)**. Cryo-electron microscopy of purified lipid 4,4’-diaponeurosporenoic acid (St1) revealed micelle-like lamellar structures **(Fig. 6C)** resembling the structure of MLBs released from bacteria.

**Fig. 6.**
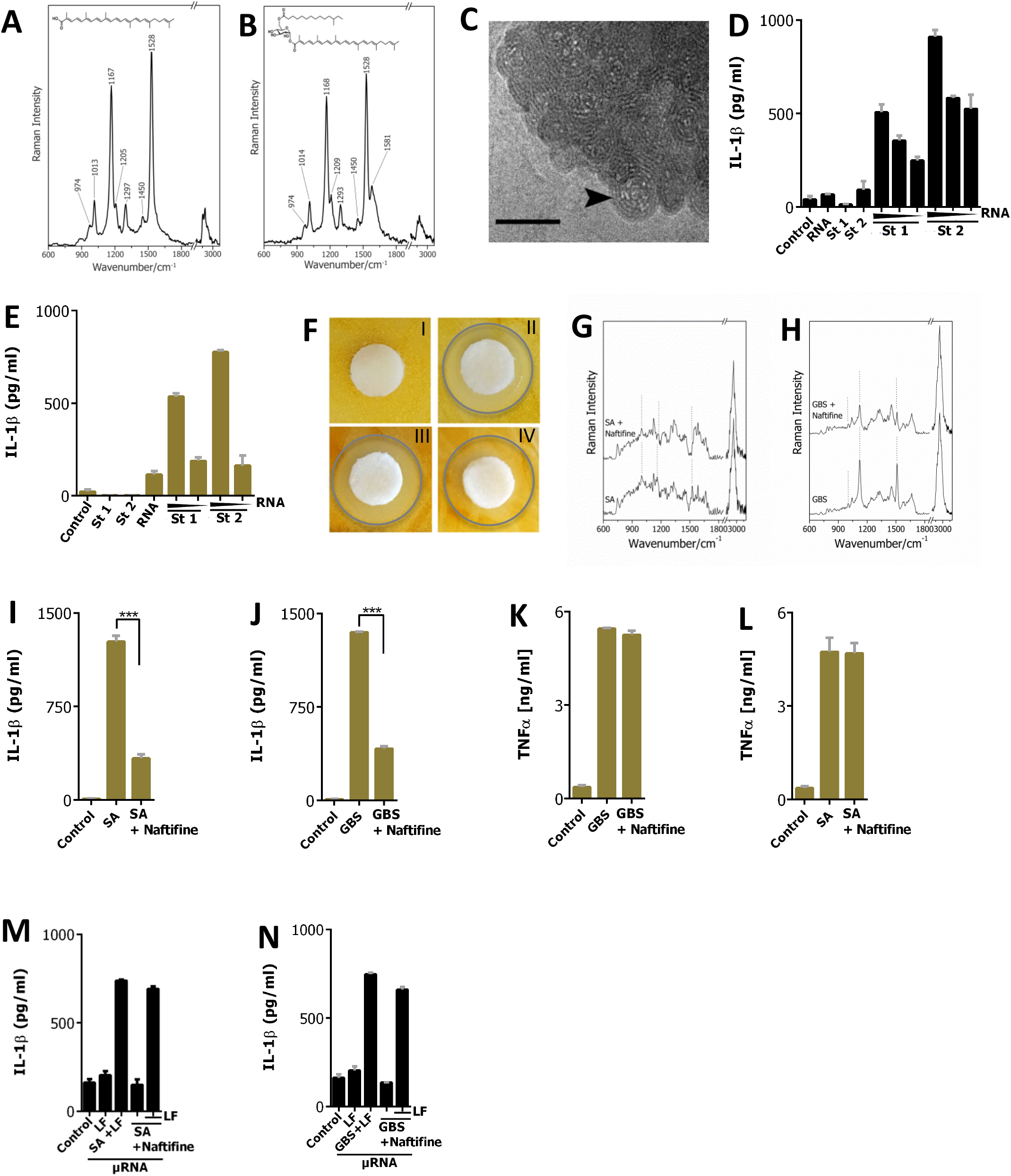
Staphyloxanthin type of lipids in MLBs can target RNA PAMPs to the cytoplasm to activate the inflammasome. **(A,B)** 4,4’-diaponeurosporenoic acid and staphyloxanthin were HPLC purified from SA followed by fingerprint LC-MS spectral analysis and confirmed by Raman spectra analysis. The prominent Raman peaks are marked with dotted lines. **(C)** Cryo TEM image of HPLC purified 4,4’-diaponeurosporenoic acid; the arrowhead indicates multi-lamellar lipid structures; the bar represents 100 nm. **(D)** IL-1β production in THP1 MΦs following stimulations with increasing concentrations of SA RNA, HPLC purified 4,4’-diaponeurosporenoic acid (St1), staphyloxanthin (St2) (0.125 µM), and 4,4’-diaponeurosporenoic acid (St1) (0.125 µM), staphyloxanthin (St2) micelles with RNA (St1+RNA, St2+RNA) (5 µg/ml, 2.5 µg/ml). **(E)** IL-1β production in human blood-derived MΦs when stimulated with 4,4’- diaponeurosporenoic acid (St1) (0.125 µM), staphyloxanthin (St2) (0.125 µM) and 4,4’- diaponeurosporenoic acid (St1+RNA) staphyloxanthin (St2+RNA) micelles with increasing concentrations of SA RNA (5 µg/ml, 2.5 µg/ml). **(F)** Inhibition of the staphyloxanthin biosynthetic pathway using the small molecular drug naftifine, **I**. Paper disc (white) with PBS as control showing no inhibition of biosynthetic pathway leading to golden yellow pigmentation over bacterial lawn **(II,III,IV)**. Increasing concentrations of naftifine (200 ng/ml, 100 ng/ml, 10 ng/ml) applied on paper disc (white) inhibiting the biosynthetic pathway leading to a zone of decolourization and opaque growth of GBS. **(G,H)** Raman spectra of **(G)** SA and SA treated with naftifine (SA+Naftifine) and **(H)** GBS and GBS treated naftifine (50 ng/ml). The fingerprint Raman peaks are marked by dotted lines. **(I,J)** Human blood-derived MΦs were infected **(I)** SA (MOI 10) and SA treated with naftifine (SA+Naftifine) and **(J)** GBS (MOI 20) and GBS treated with naftifine (GBS+Naftifine) (50 ng/ml). IL-1β was measured in supernatant. **(K,L)** Human blood-derived MΦs showing TNF-α production when infected with **(K)** GBS (MOI 10) and GBS treated with naftifine (GBS+Naftifine) and **(L)** SA (MOI 20) and SA treated with naftifine (SA+Naftifine). **(M,N)** Wild type THP1 MΦs (black) were stimulated on surface and into cytosol with naftifine treated SA and GBS µRNA (5 mg/ml) (SA (Naftifine)+LF) (GBS (Naftifine)+LF) respectively. IL-1β production was measured. Data shown are mean ±SD (n=3), representative of at least three independent experiments. Asterisks indicate statistically significant differences (∗p < 0.05, ∗∗p < 0.01 and ∗∗∗p < 0.001).

After empirical optimization of a sub-lethal purified lipid St1, St2 concentration, we examined whether St1 and St2 have the potential to deliver bacterial RNA to the cytosol. To this end St1 and St2 in the absence of stabilizers and when stimulated alone did not induce any cytoplasmic inflammasome activation **(Fig. 6D).** However RNA was embedded while rehydrating purified lipids St1, St2 and stimulated on THP1 macrophages. The St1 and St2 lipid alone or RNA alone did not activate the inflammasome while the combination of both induced IL-1β release in a concentration-dependent manner **(Fig. 6D)**. These results indicate that staphyloxanthin along with the pathway precursor 4,4’-diaponeurosporenoic acid could deliver RNA PAMPs to the cytosol to induce the inflammasome **(Fig. 6D)**. This may indicate synergism between these compounds and RNA within MLBs to activate the inflammasome. We further confirmed the capacity of staphyloxanthin to target PAMPs to cytosolic inflammasome activation using primary human blood-derived macrophages **(Fig. 6E)**. Collectively, the above data demonstrate the mechanism by which MLBs deliver PAMPs such as RNA to the cytosol with the assistance of staphyloxanthin-type lipids.

As seen in the chemical analysis, MLBs are enriched with staphyloxanthin and its precursor **(Fig. 2N-P, Supplemental Fig. 2D)**. Staphyloxanthin is an important virulence factor in SA. Bacteria lacking this carotenoid pigment grow normally but are rapidly killed by neutrophils and lack the ability to cause skin infection and abscess formation (21, 65). The biosynthetic pathway for staphyloxanthin starts with farnesyl diphosphate, a key intermediate in the isoprenoid biosynthetic pathway, and consists of six enzymes: 4,4′-diapophytoene synthase (CrtM), 4,4′-diapophytoene desaturase (CrtN), 4,4′-diaponeurosporene oxidase (CrtP), glycosyl transferase (CrtQ) and acyl transferase (CrtO) and the newly discovered 4,4′- diaponeurosporen-aldehyde dehydrogenase (AldH) (22, 23) **(Supplemental Fig. 6A)**. In *S. aureus*, five of the staphyloxanthin biosynthesis genes are arranged in a crtOPQMN operon. Hence, we inhibited 4,4′-diapophytoene synthase (CrtM) in SA using the small molecule inhibitor, naftifine (66). As expected, naftifine-treated SA turned colorless in contrast to the golden colour seen with untreated SA. Of note, similar treatment of GBS with naftifine rendered GBS colonies colorless without affecting bacterial growth **(Fig. 6F)**. Raman analysis of naftifine-treated bacteria showed a reduction in vibrations corresponding to the bacterial unsaturated lipids at wavenumber 1528, 1168, 1013 cm^-1^ and 1528, 1129, 1012 cm^-1^ for SA and GBS, respectively **(Fig. 6G, 6H)**. Inflammasome activation was reduced in naftifine- treated bacterial infection in primary human blood-derived macrophages **(Fig. 6I, 6J)** while the inflammatory response (e.g.TNFα) remained unchanged for naftifine-treated SA and GBS **(Fig. 6K, 6L)**. To gain further insights as to whether naftifine inhibition had any effect on the synthesis of µRNA aptamers, we isolated µRNA from GBS and SA treated with naftifine and stimulated to THP1 macrophages. A pronounced IL-1β secretion was seen with both treated and non-treated µRNA **(Fig. 6M, 6N)**. This shows that naftifine inhibition reduces production of staphaloxanthin biosynthetic pathway lipids. However, naftifine treatment had no effect on synthesis of RNA aptamers. Overall, the above results demonstrate an evolutionary interplay between lipid toxin and RNA PAMP to specifically activate the inflammasome to release IL-1 β.

### Gram-positive sepsis patients shows the hallmarks of MLB-mediated inflammasome activation

Organ damage and mortality in sepsis have been attributed to a deleterious and dysregulated host responses to infection (67). We investigated whether our observed immune activation pathway is conserved in clinical isolates arising from Gram-positive infections. *S. aureus* patient isolates (n=50) collected from different clinical departments were screened **(Supplemental Table 6)**. Those isolates collected from patients mainly showed an infection focus with intravascular catheter, pneumonia and wound infection **(Fig. 7A)**. Of all blood culture positive patients, 10% had sepsis (organ dysfunction) according to the most recent Sepsis-3 definition (67). When screened for the presence of RNAIII 16% of obtained bacterial patient isolates showed undetectable RNAIII. We thus isolated µRNA from RNAIII positive (P5, P11, P27) and negative (P14, P15, P36) *S. aureus* isolates. The isolated µRNA was delivered cytosolically in THP1 macrophages in equimolar concentrations. Consistent with our previous findings using *S. aureus* (LS1) and Δ*agr* strain the RNAIII negative clinical bacterial isolates (P14, P15, P36) did not activate the inflammasome pathway **(Fig. 7B)** while RNAIII positive bacterial isolates (P5, P11, P27) had a pronounced release of IL-1β **(Fig. 7B)**. These results indicate the presence of RNAIII in *S. aureus* is correlated to µRNA-dependent inflammasome activation and release of IL-1β during severe Gram-positive infections.

**Fig. 7.**
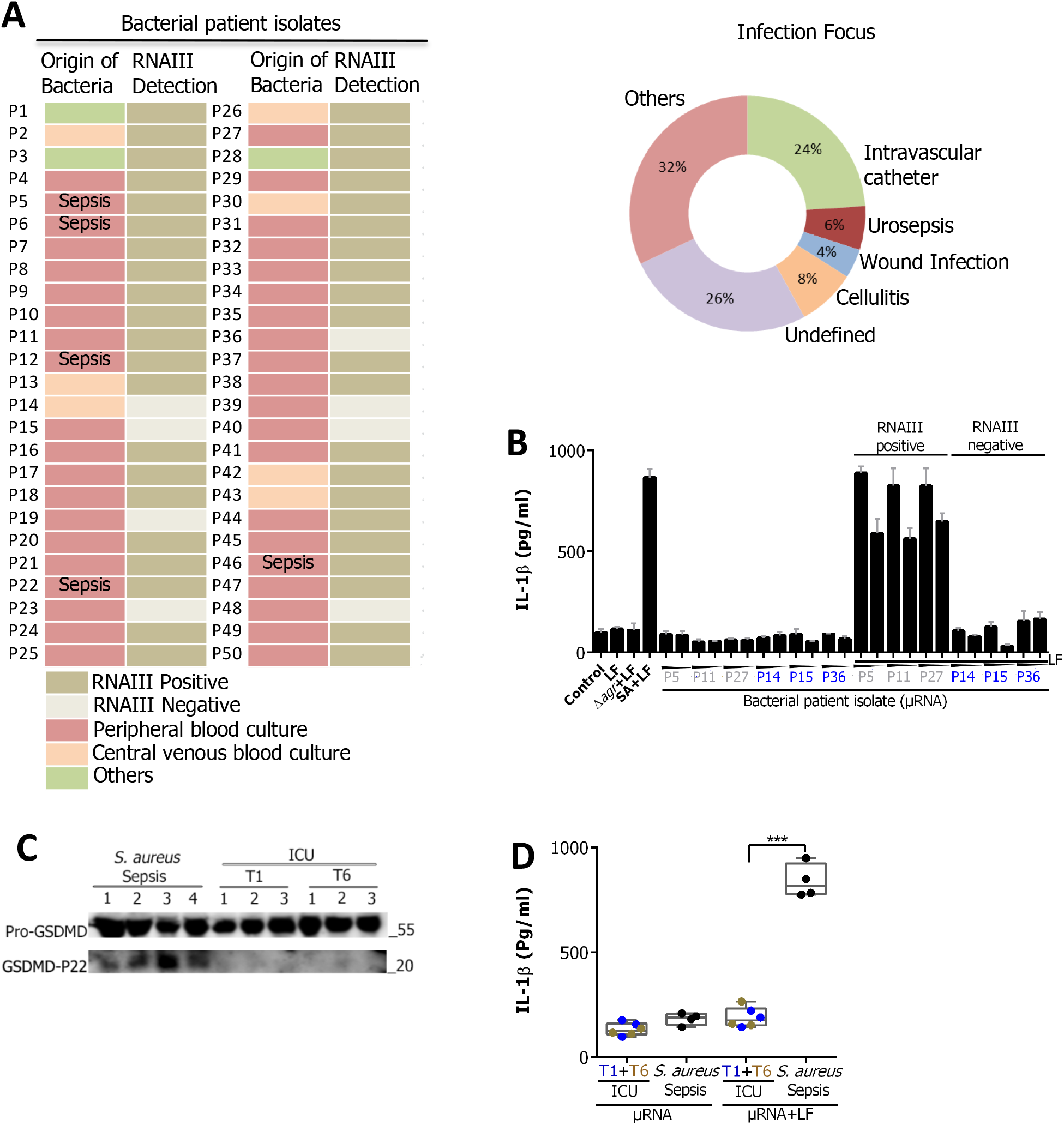
Gram-positive sepsis patients shows hallmarks of MLB-mediated inflammasome activation. **(A)** Diagrammatic representation of characteristics of *S. aureus* patient isolates (n=50) showing the presence of RNAIII (beige), absence of RNAIII (grey) and origin of bacteria. Pie diagram shows the infection focus of the patients. **(B)** THP1 MΦs (black) were stimulated with bacterial µRNA (5 µg/ml, 2.5 µg/ml) of patient isolates P5, P11, P27, P14, P15, P36 on the surface and cytosolically (P5+LF, P11+LF, P27+LF, P14+LF, P15+LF, P36+LF). IL-1β production was measured. **(C)** Immunoblots showing active GSDMD (22 kDa) in plasma of *S. aureus* sepsis patients (1- 4), ICU patients T1 (1-3), T6 (1-3). **(D)** THP1 MΦs were stimulated on the surface and cytosolically with µRNA (2.5 µg/ml) isolated from plasma of *S. aureus* sepsis patients (n=4) and ICU controls pre operation and post operation (n=3). IL-1β production was measured. Asterisks indicate statistically sign

During inflammasome activation GSDMD (55 kDa) is cleaved into N (30 kDa) and C (22 kDa) terminal subunits. The cleaved GSDMD triggers pyroptosis and a robust IL-1β production (2). To confirm inflammasome activation through GSDMD cleavage in an independent sepsis cohort we processed plasma from four *S. aureus* sepsis patients **(Supplemental Table 7)**. As controls, we collected plasma from non-septic patients undergoing cardiac surgery both pre-operatively (T1) and at 6 h post-operatively (T6). On western blot analysis, plasma from the sepsis patients showed two distinct bands at 55 and 22 kDa, indicating cleaved GSDMD, while the non-septic ICU controls did not show this GSDMD cleavage **(Fig. 7C)**. We subsequently isolated µRNA from the plasma of *S. aureus* sepsis patients and non-septic controls and cytosolically delivered this µRNA into THP1 macrophages to determine inflammasome activation. µRNA from *S. aureus* sepsis patients elicited a pronounced IL-1β response as compared to the cardiac surgery controls **(Fig. 7D)**. The surface stimulation of µRNA from patients and controls failed to release IL-1β **(Fig. 7D)**. These details indicate that plasma from septic patients with *S. aureus* infection may contain hallmark RNA aptamers that could activate the inflammasome pathway **(Fig. 7D)**. Taken together, our translational results suggest that *S. aureus* µRNA-dependent cytosolic inflammasome induction is an important feature of Gram-positive sepsis.

## DISCUSSION

Sepsis is a dysregulated host response to infection leading to life-threatening organ dysfunction (67). With more than 19 million cases annually, sepsis is a leading cause of death worldwide and a major socioeconomic burden (68, 69). Activation of cytosolic caspases leading to inflammasome activation is emerging as a key host defense strategy for clearance of bacteria. Under this evolutionary pressure both bacteria and host may develop new strategies for regulated inflammasome activation. GBS uses granadaene and RNA to activate the inflammasome (20, 64). How these PAMPs orchestrate activation and whether any upstream mechanisms are induced by bacteria, are still uncertain (69).

In this current study, we demonstrated that GBS and SA utilize µRNA aptamers for downstream activation of the canonical inflammasome pathway. The specialized µRNA aptamers with capacity to bind CDN when delivered to host cytoplasm, activate the STING- dependent canonical inflammasome to release IL-1β.

STING is an endoplasmic reticulum resident protein activated by second messenger CDN ligands for pronounced interferon responses (6, 16). Canonical inflammasome induction following activation of STING is known in literature. Moreover few mechanisms of how STING activation propagates to inflammasome activation are also shown (87, 88, 36, 70). For DNA activators of NLRP3, a cGAS and STING-dependent lysosomal pathway has been shown in human cells (36). Evidence is lacking for RNA activators and it remains uncertain how Gram-positive bacteria use these pathways during infection. Our data suggest that bacterial RNA PAMPs activate STING-dependent IL-1β responses on arrival in the cytoplasm. Small RNA (<200 nucleotides) which comprises CDN RNA aptamers may sequester CDN ligands to STING for NLRP3 activation, albeit bypassing cGAS **(Fig. 4I, 4J)**.

The main findings of our work are the delivery of CDN to host cells for activation of STING by Gram-positive bacteria through MLBs, and the involvement of RNA aptamers in this process. Consistent with this observation, we found that CDN binding RNA aptamer-mediated STING activation is a dominant pathway for canonical inflammasome activation during Gram-positive bacterial infection in human macrophages. The possibility of a redundant involvement of other cytosolic receptors cannot be ruled out as nucleic acid signalling shows cell type-and host species-type specificities. The co-localization of STING and LAMP1 upon Gram-positive bacterial infection or stimulation with CDN aptamers suggests that, upon activation by RNA aptamers, STING translocates to the lysosome. This coincides with a fall in cytoplasmic K^+^ levels and activation of the NLRP3 inflammasome. STING activation and downstream NLRP3-dependent inflammasome activation have been poorly studied to date. Our study provides important clues as to how Gram-positive bacteria deliver CDN to activate STING. The investigation of STING activation propagating to the NLRP3 merits further study in the light of the current findings (71).

The concept of inflammasome activation by cytosolic bacterial RNA raises the question as to how microbial RNA can gain access to this compartment. To activate cytosolic PRRs, specific PAMP delivery mechanisms are required. Here, we demonstrate that Gram-positive bacteria release at least two populations of vesicles named membrane vesicles (MVs) and multi- lamellar lipid bodies (MLBs). Similar vesicles have been reported for other organisms e.g. mycobacteria (72). A large heterogeneity of EVs secreted by Gram-positive bacteria has also been reported (32, 34). However, the current literature lacks both structural and functional analysis of these vesicles. GBS EVs have recently been implicated in causing premature birth in a mouse model (35). In this current study we structurally characterized MVs and MLBs secreted by GBS and SA and demonstrated differences in both size and lipid content. Biochemically, MLBs have abundant polyunsaturated lipids including those from the staphyloxanthin biosynthetic pathway. These lipids are highly hydrophobic and, consequently, may contribute to the overall hydrophobicity of MLBs. Structurally, they contain concentric shells of lipids with RNA PAMPs packed within. We also found that Gram-positive MLBs show phase transition properties **(Supplemental Fig. 2F, 2G)** providing specific biophysical properties to Gram-positive vesicles. MLBs replicate an important and similar function as live bacteria in inducing the inflammasome. Consistent with this we report that MLBs can activate the canonical pyroptotic cell death pathway and is dependent on caspase-1, ASC and NLRP3 **(Fig. 3C, 3D)**. While understanding the activation pattern of MLBs, how the bacterial PAMPs get packed within the MLBs remains unclear.

Packaging of distinct RNA in the vesicles is a known general phenomenon (48, 49). We identified enrichment of less than 200 nucleotide µRNA in the SA MLBs. The RNA content of the MLBs differs remarkably to the originating bacteria when analysed through deep sequencing **(Fig. 5A, Supplemental Fig. 5A-D)**. This may relate to active mechanisms within bacteria that process, sort and pack µRNA into the MLBs (48, 49). Moreover, RNA sequencing showed levels of sbr-C, Rsa-C, RNAIII and WAN01CC66-rc were *AGR* dependent and entirely expressed in SA and MLBs **(Supplemental Fig. 5D)**. A well-characterized RNAIII in *S. aureus* acts via trans-acting factors as it has the possibility to form a secondary structure for the binding of proteins (56, 58). A RNAIII central domain comprising hairpin 7 has a potential recognition site for trans-acting factors due to the conserved sequence of UCCCAA which makes it of particular interest (59). Our analysis also demonstrates a selective enrichment of central domain A,7,8,9,A of RNAIII within MLBs. Detailed analysis revealed that the structure formed by the central domain A,7,8,9,A of RNAIII is essential for activation of the inflammasome pathway **(Supplement Fig. 5F, 5H, 5I)**. Mutations in the central domain hampers the structure and fails to activate the inflammasome **(Supplemental Fig. 5H, 5I)**. Moreover, the central domain of RNAIII when fused with spinach aptamer showed CDN-binding capacity with varying concentrations of RNA and CDN. Altogether, MLB-mediated release of RNA aptamers within the cytosol to activate the inflammasome may be an important conserved mechanism within host-pathogen interactions.

Bacterial EVs are taken up by immune cells through a clathrin-mediated endocytic pathway that results in activation of caspase-11 (34). Gram-positive MLBs also follow a similar pathway for cellular entry. Their entry is dynamin-dependent as inhibition of dynamin leads to the loss of inflammasome induction by MLBs **(Fig. 3I)**. EVs may thus carry specific molecular components that help to transfer entrapped PAMPs to the cytosol. MLBs were enriched with polyunsaturated lipids **(Supplemental Fig. 2D, 6A)**. These lipids are produced by the staphyloxanthin and granadaene biosynthesis pathway in SA and GBS, respectively (22, 23, 65). Staphyloxanthin and its pathway intermediate, 4,4’-diaponeurosporenoic acid have strong capacities for transferring PAMPs to the cytosol **(Fig. 6D, 6E)**. This indicates potential new mechanisms by which these bacterial metabolites participate in bacterial pathogenesis. Interestingly, inhibition of crtM, a key enzyme in staphyloxanthin biosynthesis, reduced the ability of SA to activate the inflammasome without affecting the bacterial capacity to produce RNA aptamers **(Fig. 6I, 6J, 6M, 6N)**. These findings reveal the possibility of developing targeting drugs or chemical probes to control MLB-mediated effects during Gram-positive sepsis.

During the course of our investigation staphyloxanthin and 4,4’-diaponeurosporenoic acid from SA were HPLC-purified and characteristic Raman vibrational fingerprint spectra were determined for these molecules. Our determined Raman spectra of staphyloxanthin and 4,4’- diaponeurosporenoic acid molecules add to the current knowledge of microbial pigments and their properties. Staphyloxanthin types of lipids from SA were found to be pathogenic in combination with PAMPs within MLBs.

On the other hand, the same MLB lipids when embedded with PAMPs showed pronounced induction of the canonical inflammasome. These findings complement and advance current knowledge where pathogenic bacteria may induce over-activation of inflammatory processes through production of specific lipids and RNA, whereas absence of staphyloxanthin (47) or RNAIII (57) may render *S. aureus* an asymptomatic colonizer (21, 66). In a similar hypothesis, the balance of IL-1β and Type-I interferon responses could regulate a successful host defense against *S. pyrogenesis* (73). Our studies in combination with others (16, 74) suggest cGAS-STING pathway is a central pathway regulated by pathogenic Gram-positive bacteria that causes the imbalance in IL-1β and Type-I interferon responses. Consistent with this, staphyloxanthin type lipid biosynthetic pathways have evolved in many Gram-positive organisms. Strikingly, in the case of *S. aureus*, loss of its staphyloxanthin producing ability leads to bacterial clearance by lymphocytes (21). Corroborating these observations, the presence of RNAIII was detected in isolates from *S. aureus* infected patients. µRNA from the RNAIII positive patient isolates could profoundly activate the cytosolic inflammasome pathway in comparison with RNAIII negative patient isolates **(Fig. 7B)**. These results highlight the conserved nature of our findings in bacteria originating from clinical settings. Moreover, it may also indicate an association between the presence of RNAIII and inflammasome activation in *S. aureus* patients. To study these findings further we found µRNA from plasma of *S. aureus* sepsis patients activated the inflammasome with abundant release of IL-1β, whereas µRNA isolated from control patient plasma failed to activate the inflammasome **(Fig. 7D)**. Translating these findings further, the same plasma from *S. aureus* sepsis patients showed-hallmarks of the activated inflammasome in terms of cleaved GSDMD-P22 whereas, control patients failed to show an activated inflammasome and cleaved GSDMD **(Fig. 7C)**. Hence, all the above findings may provide a new perspective for the pathogenicity of Gram-positive bacterial infection in man.

## MATERIALS AND METHODS

### Bacterial strains and growth conditions

GBS wild type strain NEM 316, *S. aureus* wild type strain LS1 and isogenic mutant Δ*agr* were grown overnight in THY Medium (Todd-Hewitt Broth with 2% yeast extract) at 37°C. The overnight cultures were further diluted and grown untill the late logarithmic phase. *S. aureus* patient isolates and *E. coli* patient isolates were similarly grown in THY Medium and LB broth, respectively at 37°C 160 rpm.

### Purification and characterization of bacterial MLBs and OMVs

MLBs were purified from GBS, SA and OMVs were purified from *E. coli*, with some modifications (75, 76). In brief, GBS and SA were grown in THY at 37°C 160 rpm. The culture was centrifuged at 10,000 rpm for 15 min at 4°C. The bacteria-free supernatant was further filtered through a 0.22 µm filter. Bacterial vesicles were isolated from the supernatant by ultracentrifugation at 400,000 g for 2 h at 4°C in Beckman (TLA-100.3 rotor). The pelleted vesicular fraction was then washed with sterile PBS for 1 h. For purification of MLBs and MVs, vesicular fraction was further purified by opti-prep gradient from 45 to 15% made in 10 mm HEPES, 0.85% NaCl followed by 200,000 g, 2 h ultracentrifugation and different fractions were collected and washed. MLBs were enriched at a density of approximately >1.20 g/ml and showed fingerprint Raman vibrations at 1163 and 1528 cm^-1^ for SA, and 1129 and 1524 cm^-1^ for GBS. Each preparation of MLBs underwent a quality check by (i) Agar plating, (ii) electron microscopy, (iii) DLS (Dynamic Light Scattering) and (iv) Raman spectroscopy. Despite of the above mentioned quality measurements we cannot exclude the possibility that some MVs may appear within the MLB fraction or *vice versa*. The protein contents of the isolated MLBs were determined by a Pierce BCA protein assay kit (Thermo Scientific), according to the manufacturer’s instructions.

### Raman measurement

Isolated MLBs were washed with Millipore water and smeared onto a calcium fluoride slide. For the bacterial measurement SA and GBS were streaked on agar plates and incubated overnight. A single colony was smeared onto a calcium fluoride slide. Raman spectra were recorded with an upright micro-Raman set-up (CRM 300, WITec GmbH, Germany) equipped with a 600 g/mm grating and a deep depletion CCD camera (DU401A BV-532, ANDOR, 1024 x 127 pixels) cooled down to -60°C. Samples were excited through a Nikon 100x objective (NA 0.8) using an excitation wavelength of 532 nm Nd-YAG laser (20mW of laser power and exposure time of 1s per spectrum). Single Raman spectra were collected from 300 different sampling positions either in time series mode or single accumulation.

Prior to analysis Raman spectra exhibiting only a fluorescence background were removed and spectral regions, finger print (600 to 1800 cm^-1^) and C-H stretching (2750 to 3100 cm^-1^) of Raman spectra were further used for analysis. Data pre-processing and statistical analysis were carried out. Raman spectra were background corrected using the sensitive nonlinear iterative peak (SNIP) clipping algorithm and vector normalized. Pre-processed Raman spectra were carefully sorted as per dominating biomolecules allowing the vibrational signatures and average Raman spectra of proteins, lipids and nucleic acids to be then generated **(Supplemental Table 1)**.

### Electron microscopy

For negative staining transmission electron microscopy (TEM) carbon-coated EM-grids (400 meshes, Quantifoil, Großlöbichau, Germany) were hydrophilized by glow discharging at low pressure in air. 20 μl of MLB or OMV solution were adsorbed onto the hydrophilic grids for 2 min. The grids were washed twice on drops of distilled water and stained on a drop of 2% uranyl acetate in distilled water.

For freeze-fracture transmission electron microscopy (FEM) aliquots of cell suspension or isolated lipid fractions were enclosed between two 0.1-mm-thick copper profiles as used for the sandwich double-replica technique. The sandwiches were physically fixed by rapid plunge freezing in a liquid ethane/propane mixture, cooled by liquid nitrogen. Freeze-fracturing was performed at -150°C in a BAF400T freeze-fracture unit (BAL-TEC, Lichtenstein) using a double-replica stage. The fractured samples were shadowed with 2 nm Pt/C (platinum/carbon) at an angle of 35°, followed by perpendicular evaporation of a 15-20 nm thick carbon layer. The evaporation of Pt/C was controlled by a thin-layer quartz crystal monitor; the thickness of the carbon layer was controlled optically. The obtained freeze-fracture replicas were transferred to a “cleaning solution” (commercial sodium hypochlorite containing 12% active Cl2) for 30 min at 45°C. Then, the replicas were washed four times in distilled water and transferred onto Formvar-coated grids for examination under a transmission electron microscope. The negative staining and freeze-fracture samples were both imaged in a Zeiss EM902A electron microscope (Carl Zeiss AG, Oberkochen, Germany) operated at 80 kV accelerating voltage, and images were recorded with a 1k (1024 x 1024) FastScan-CCD-camera (CCD-camera and acquisition software EMMANU4 v 4.00.9.17, TVIPS, Munich, Germany).

For scanning electron microscopy (SEM) aliquots of cell suspension were fixed in 2.5% glutaraldehyde in cacodylic acid buffer (100 mM, pH 7.2) for 1 h. In order to avoid lipid extraction, dehydration by ethanol series and critical point drying steps were omitted and drops of glutaraldehyde fixed cell suspensions were air-dried on glass cover slips. The glass cover slips were mounted on aluminium sample holders and gold sputter coated (layer thickness 20 nm) in a Compact Coating Unit CCU-010 (Savematic GmbH, Bad Ragaz, Switzerland). The samples were examined in a Zeiss LEO 1530 SEM (Carl Zeiss AG, Oberkochen, Germany) at 8 kV acceleration voltages and a working distance of 3 mm using an InLense secondary electron detector.

### Dynamic light scattering

Size distribution and varying diameters of the isolated MLBs and MVs were measured using Zeta sizer nano series. The diluted MLBs and MVs samples of SA were measured at 25°C for size. Continuous measurement from 15°C to 60°C with the interval of 5°C was performed for the phase transition experiment.

### LC-MS of staphyloxanthin type of lipids in MVs and MLBs

The derivatives of staphyloxanthin were detected with LC-MS from MVs, MLBs and SA bacteria. Due to the light sensitivity of staphyloxanthin derivatives, all the extraction processes are performed in the dark condition or under the protection of aluminum foil. Bacterial pellet of SA grown until the late logarithmic phase was first extracted by 50 mL EtOH, and ultrasonicated for 20 min until the pellet was nearly colorless and all the pigments were extracted in EtOH. After centrifugation at 8000 rpm for 10 min, the EtOH extract was collected, filtrated, and finally concentrated under reduced vacuum until completely dried. The EtOH extract was further separated by solvent extraction with CHCl3: MeOH: H2O 2:2:1 (v/v/v) and thorough mixing by vortex. After centrifugation, two layers were clearly separated and the bottom layer was collected as a lipid extract and concentrated under reduced vacuum. The lipid extract was suspended into MeCN to a concentration of 1 mg/mL. After centrifugation, the clear supernatant was submitted to Shimadzu UHPLC-MS with the linear gradient: 0-1 min, 10% B; 1-7 min, 10%-100% B; 7-10 min, 100% B (A: dd H2O with 0.1% formic acid; B: MeCN with 0.1% formic acid) with a flow rate of 0.7 mL/min. 5 μL sample solution was injected into UHPLC-MS for analysis.

MVs and MLBs were prepared from the above protocol and suspended into 25 µL H2O, and then added into 25 µL MeOH, and ultra sonicated for 10 min. After centrifugation at 13000 rpm for 10 min at room temperature, the supernatant was transferred into brown HPLC vials and directly submitted to Shimadzu UHPLC-MS using the same procedure as described above. In the meantime, metabolite analysis was carried out on Thermo QExactive Plus HESI-HRMS equipped with a Luna Omega C18 column (100 x 2.1 mm, particle size 1.6 µm, pore diameter 100 Å, Phenomenex) preceded by a Security Guard™ ULTRA guard cartridge (2 x 2.1 mm, Phenomenex) with the linear gradient: 0-0.5 min, 70% B; 0.5-17 min, 70%-97% B; 17-22 min, 97% B (A: dd H2O with 0.1% formic acid; B: MeCN with 0.1% formic acid) with a flow rate of 0.3 mL/min. The column oven was set to 40 °C. 5 µl of the sample were submitted for analysis and metabolite separation was followed by a data-dependent MS/MS analysis in positive (MS^1^ and MS^2^) ionization mode. The gas flow rates were set to 35 and 10 for the sheath and auxiliary gases, respectively. The capillary and probe heater temperatures were 340°C and 200°C, respectively. The spray voltages were 4 kV for the positive ionization modes. S-lens RF level was set to 50. MS^1^ had the resolving power set to 70,000 FWHM at *m*/*z* 200, scan range to *m*/*z* 150 – 2,000; injection time to 100 ms; and AGC to 3e6. The ten most intense ions were selected for MS^2^ with a scan rate of 12 Hz. Resolving power was 17,500 FWHM at *m/z* 200, AGC target was 1e5, and injection time was 50 ms. Fragmentations were performed at 28 NCE (normalized collision energy). Data analysis was performed with Thermo XCalibur software (Thermo Scientific).

### LC-MS of c-di-AMP and it’s derivatives in MLBs

Ultra-high performance liquid chromatography coupled with high resolution mass spectrometry was carried out using a THERMO (Bremen, Germany) UltiMate HPG-3400 RS binary pump, WPS-3000 auto sampler set to 10°C and equipped with a 25 µL injection syringe and a 100 µL sample loop. The column was kept at 25°C within the TCC-3200 column compartment. A PHENOMENEX^®^ (Aschaffenburg, Germany) Hydro-RP (80 Å pore size, 150 × 2 mm; 4 µm particle size) Chromatography column was used **(Suplemental Table 3)** at a constant flow rate of 0.4 mL/min. Eluent A was water, with 2% acetonitrile and 0.1% formic acid. Eluent B was pure acetonitrile. For each sample 5 µL were injected prior to dilution with 10 µL UPLC-grade water.

Mass spectra were recorded with THERMO QExactive plus an Orbitrap mass spectrometer coupled to a heated electrospray source (HESI). Column flow was switched at 0.5 min from waste to the MS and at 11.5 min again back to the waste, to prevent source contamination. For monitoring two full scan modes were selected with the following parameters. Polarity: positive; scan range: 100 to 1500 *m*/*z*; resolution: 70,000; AGC target: 3 × 10^6^; maximum IT: 200 ms. General settings: sheath gas flow rate: 60; auxiliary gas flow rate 20; sweep gas flow rate: 5; spray voltage: 3.0 kV; capillary temperature: 360°C; S-lens RF level: 50; auxiliary gas heater temperature: 400°C; acquisition time frame: 0.5 - 11.5 min. For negative mode all values were kept instead at spray voltage set to 3.3 kV.

Presence of pApA was confirmed based on their [M−2H+Na]^−^ and c-di-AMP based on [M−H]^−^; respectively using their exact masses and a 10 ppm mass window. Presence was further verified by at least two of additionally occurring ions ([M−2H+Na]^−^; [M−H]^−^; [M+H]^+^; [M+Na]^+^; [M−H+2Na]^+^ ) within a retention time range of ± 0.2 min comparing to the authentic standards. Peak detection and integration were carried out using the THERMO Xcalibur^TM^ 3.0.63 software.

### Purification of 4,4’-diaponeurosporenoic acid and staphyloxanthin from *S. aureus*

NMR measurements were performed on a Bruker AVANCE III 600 MHz spectrometer, equipped with a Bruker Cryoplatform. Chemical shifts are reported in parts per million (ppm) relative to the solvent residual peak of CDCl3 (^1^H: 7.26 ppm, singlet; ^13^C: 77.16 ppm, triplet). LC-ESI-HRMS measurements were carried out on an Accela UPLC system (Thermo Scientific) coupled with a Accucore C18 column (100 x 2.1 mm, particle size 2.6 µm) combined with a Q-Exactive mass spectrometer (Thermo Scientific) equipped with an electrospray ion (ESI) source. UHPLC-MS measurements were performed on a Shimadzu LCMS-2020 system equipped with single quadruple mass spectrometer using a Phenomenex Kinetex C18 column (50 x 2.1 mm, particle size 1.7 μm, pore diameter 100 Å). The column oven was set to 40°C; the scan range of MS was set to m/z 150 to 2,000 with a scan speed of 10,000 u/s and event time of 0.25 s under positive and negative mode. DL temperature was set to 250°C with an interface temperature of 350°C and a heat block of 400°C. The nebulizing gas flow was set to 1.5 L/min and dry gas flow to 15 L/min. Semi-preparative HPLC was performed on a Shimadzu HPLC system using a Phenomenex Luna Phenyl Hexyl 250 x 10 mm column (particle size 5 μm, pore diameter 100 Å). Solid phase extraction was carried out using Chromabond SiOH cartridges filled with 2 g of unmodified silica gel (Macherey-Nagel, Germany). Chemicals: Methanol (VWR, Germany); water for analytical and preparative HPLC (Millipore, Germany), formic acid (Carl Roth, Germany); acetonitrile (VWR as LC- MS grade), media ingredients (Carl Roth, Germany).

SA was cultured overnight in THY medium (37°C, 160 rpm). The cell pellet was separated from the supernatant by centrifugation at 4000 rpm at 4°C. The cell pellet was extracted twice using 100 mL ethanol at 37°C for 2 h. The EtOH extract was combined and concentrated under reduced pressure. The dried extract was partitioned using 100 mL 1.7 M NaCl and 100 mL ethyl acetate three times. The EtOAc phase was combined, dried over MgSO4 and concentrated under reduced pressure conditions. In total, 189 mg crude extract was obtained from 10 L liquid culture. The crude extract was separated using a SiOH cartridge (2 g, twice) and eluted by a step gradient of cyclohexane and EtOAc (100% cyclohexane, 5:1, 2:1, 1:1, 1:2, 100% EtOAc and 100% MeOH). The fraction resulting in an orange band was collected from the elution of cyclohexane: EtOAc 5:1 (v/v) (Fraction 2, 36.22 mg) and 1:1 (v/v) (Fraction 5, 4.58 mg).

Fr. 2 was further separated on RP-C18 F254s preparative TLC plates (Merck) using a methanol-acetonitrile (1:1, v/v) mixture. The pigment band with an Rf value of 0.61 was scratched from the preparative TLC plates, extracted by 1 x 3 mL EtOAc, and concentrated under reduced pressure to yield Fr. 2.1, (4.29 mg). Fr. 2.1 was subjected to purification by semi-preparative HPLC on Phenomenex Phenyl-Hexyl column 250 x 10 mm, 100 Å. The fraction containing 4,4’-diaponeurosporenoic acid (**1**, 0.79 mg, *tR* = 13.92 min) was obtained using a gradient of 0-5 min, 90% MeCN/10% H2O containing 0.1% formic acid; 5-10 min, 90%-100% MeCN; 10-20 min, 100% MeCN, under a flow rate of 2.0 mL/min. Similarly, Fr. 5 was further separated with a pigment band with an Rf value of 0.55 scratched from the preparative TLC plates and extracted three times with 1 mL EtOAc. The EtOAc extract of the pigment band (Fr. 5.1, 1.6 mg) was concentrated under reduced pressure and subjected to purification by semi-preparative HPLC on a Phenomenex Phenyl-Hexyl column 250 x 10 mm, 100 Å. Staphyloxanthin (2, 0.1 mg, *tR* = 19.54 min) was obtained by semi-preparative HPLC under a gradient of 0-5 min, 90% MeCN/10% H2O containing 0.1% formic acid; 5-10 min, 90%-100% MeCN; 10-20 min, 100% MeCN, under a flow rate of 2.0 mL/min.

The sample was analysed using a Shimadzu UHPLC-MS (gradient: 0–1 min, 70% B; 1-5 min, 70%-100% B; 5-7 min, 100% B (A: dd H2O with 0.1% formic acid; B: MeCN with 0.1% formic acid) with a flow rate of 1.0 mL/min. 1 μL sample solution was injected into the UHPLC-MS for analysis.

### RNA extraction and quality control

RNA was extracted from GBS, SA and MLBs from GBS and SA, respectively using a Qiagen RNA extraction kit with some modifications. Briefly, samples were homogenized in Qiazol (Qiagen) for 1 min with a minute of cool-down step using a homogenizer. The total RNA from each sample was collected into two fractions using two column methods. Big RNA (BRNA), which contains more than 200 nucleotides, was isolated using total RNA extraction columns with 1 volume of 70% ethanol followed by washes as described in the Qiagen total RNA extraction kit instructions. Flow through from the BRNA was collected and mixed with

1.5 volume of 100% ethanol to isolate µRNA which contains RNA shorter than 200 nucleotides. This was followed by 65°C warm buffer washes and elution as mentioned in the kit. The quantity of extracted RNA for the individual fractions ranged from 0.5 µg to 3 µg. The size distribution of the BRNA and µRNA was measured on a bioanalyser (QIAxcel Advanced systems, Qiagen) following the manufacturer’s instructions.

### RNA sequencing for wild type *S. aureus* (SA), Δ*agr* mutant and SA MLB

µRNA was isolated from SA, SA-MLB and SA-Δ*agr* as mentioned in the RNA extraction protocol. The respective isolated µRNA was quantified with a A260/A280 ratio more than 1.8. RNA integrity was subsequently assessed using Qubit RNA HS assay (ThermoFisher scientific) with a Qubit fluorometer. Dephosphorylation was performed on the respective µRNA with addition of 5µg to a Quick dephosphorylation kit (New England BioLabs). Dephosphorylated µRNA was again purified using µRNA isolation columns. The eluted µRNA was quantified again using Qubit RNA HS assay. Libraries were prepared using smRNA-seq kit (TAKARA) with the input of 1 µg µRNA for all three samples with positive and negative controls. For the library preparation the manufacturer’s instructions were followed. These included polyadenylation and, cDNA synthesis following PCR and clean-up. cDNA libraries obtained were quantified using a Qubit DNA HS assay with a Qubit fluorometer. Each library prepared yielded aprroximately 18-20 ng/µl of cDNA. The pooled library was sequenced using Illumina dual sided 250 base pair sequencing. Approximately 3 million reads of SA, SA-Δ*agr* and SA-MLB µRNA, respectively were obtained covering the entire µRNA region. Sequenced reads for SA, Δ*agr* and SA-MLB were mapped on reference sequences of *Staphylococcus aureus* subsp. aureus N315 (accession number: NC_002745.2) and *Staphylococcus aureus* subsp. aureus NCTC 8325 (accession number: NC_007795.1) and were analysed using SeqMan Pro ArrayStar and GenVision Pro followed by identification of at least 2 fold differentially abundant reads. The reads obtained from rRNA, antisense and coding regions were eliminated.

### RNAIII cloning

RNAIII which is 514 bp long was PCR amplified on the genome of *S. aureus* (LS1) with primers containing either HindIII or EcoRI restriction sites on the 3’ or 5’ ends of RNAIII respectively **(Supplemental Table 5)**. The digested PCR product of RNAIII was cloned into pET28a vector. Ligated products were transformed into *E. coli* competent cells. The transformed bacteria resistant to kanamycin were selected. Colonies from the agar plate were subjected to plasmid extraction using a Macherey Nagel Plasmid Isolation Kit, according to the manufacturer’s instructions. To confirm the accuracy of the constructs, the recombinant plasmids were sequenced.

### *In-vitro*-transcription

Templates for the *in-vitro*-transcription were amplified by PCR from a RNAIII clone using primers mentioned in **Supplemental Table 5**. *In-vitro-*transcription was carried out using the Hi-Scribe RNA transcription kit (New England BioLabs). Briefly, 10 mM NTPs, 1× reaction buffer, 800 ng DNA template and 2 μl HiScribe™ T7 polymerase in 20 μl RNase-free water with RNase inhibitor was incubated at 37°C for 2h. 2U of DNase treatment was given for 20 min. The RNA transcripts were further precipitated in 100% ethanol with sodium acetate for 16 h at -20°C. The precipitated RNA was further purified using RNA purification columns following the manufacturer’s instructions (Qiagen). The RNA columns were washed twice with warm buffer heated at 65°C. Quality control of IVT RNA was performed using QIAxcel capillary electrophoresis fragment analysis protocol to ensure lack of poor transcription template contamination and degraded RNA. Assurance of RNA purity and lack of contaminating DNA was further affirmed in the cellular assays using WT and cGAS deficient THP1 macrophages which can differentiate DNA dependent responses.

### Human blood and tissue sampling

Samples were collected in a clinical cohort study performed on the multidisciplinary intensive care unit (ICU) of Jena University Hospital (77). All patients admitted to the ICU were screened within 2h of admission for evidence of a systemic inflammatory response syndrome (SIRS) resulting from possible or proven infection. When sepsis was diagnosed, all adult patients with organ dysfunction according to the old criteria for sepsis (Sepsis-2) were eligible for study inclusion. All samples are thus in accordance with the new Sepsis-3 definition (67).

Blood samples were collected within 24 h after clinical diagnosis. After approval by the local ethics committee (IDs: 2160-11/07, 2712-12/09 and 3824-11/12), all patients or legal surrogates gave informed consent for genetic analyses, blood collection and data evaluation. Peripheral blood mononuclear cells (PBMCs) were isolated from heparinized peripheral blood from consenting healthy volunteers. PBMCs were isolated with density gradient centrifugation using Bicol separating solution and subjected to short erythrocyte lysis. Macrophages were differentiated from the isolated PBMCs with recombinant human M-CSF (10 ng/ml). On day 4, the cytokine-supplemented medium was refreshed. Macrophages were plated onto fresh 96 well plates in complete medium and then stimulated.

### Cell culture

THP1 wild type, *CASP4^-/-^*, *CASP1^-/-^*, *STING^-/^*^-^, *cGAS^-/-^*, *ASC^-/-^* and STING^R232^ expressing *STING^-/-^* cells were cultivated in RPMI Medium 1640 supplemented with L-glutamine, sodium pyruvate, 10% (v/v) FCS (life technologies) and 100 U/ml of penicillin streptomycin. Cells were stimulated in fresh medium as indicated. THP1 *CASP4^-/-^*, *CASP1^-/-^*, *ASC^-/-^* cells with caspase-4, caspase-1 and ASC gene deletion were reported previously and genotypically and phenotypically were confirmed by genome sequencing and Western blot (36). THP1 *STING^-/-^*, *cGAS^-/-^*, STING^R232^ expressing *STING^-/-^* were purchased from Invivogen. The deletion of STING and cGAS gene was confirmed with Western blot and sequencing (data not shown). Cells were differentiated into MΦ overnight using 100 ng/ml phorbol 12- myristate 13 acetate (PMA) then washed with PBS and used for experiments. All cell lines and respective gene targeted clones were tested as free from mycoplasma.

### Cell stimulation

THP1 wild type, *CASP4^-/-^*, *CASP1^-/-^*, *STING^-/-^*, *cGAS^-/-^*, *ASC^-/-^* and STING^R232^ MΦ cells were used to access the cytosolic inflammasome and cell death responses. Unless otherwise indicated cells were primed with 400 ng/ml of Pam3CSK4 (Invivogen) when LPS was the second stimulus or with 100 ng/ml LPS (Invivogen) for 3 h.

For MLB stimulation: THP1 wild type, *CASP4^-/-^*, *CASP1^-/-^*, *ASC^-/-^*, *STING^-/-^*, and *cGAS^-/-^* MΦ were washed with PBS and stimulated with 10 µg/ml MLBs. For control purposes cells were stimulated with cytosolic 1 µg LPS using lipofectamine or nigericin (6.7 µM) (InvivoGen, tlrl-nig). 1% triton X100 was used as a lysis control in cell death assays (37). Cytosolic RNA with lipid: THP1 MΦ were stimulated with RNA (5, 2.5, 0.1 µg/ml) micelled with 0.125 µM of isolated lipid. In experiments with lipid stimulations, the respective solvent control was used which was then normalized. IVT RNA stimulations: THP1 wild type, *STING^-/-^* and *cGAS^-/-^* MΦ were stimulated with *in-vitro*-transcribed RNA (5, 2.5 µg/ml) using lipofectamine. For bacterial infections: cells were infected for 1 h with bacteria grown to a late log phase *S. aureus*, mutant Δ*agr* with MOI 10 and GBS with MOI 20 unless mentioned otherwise. The medium was replaced with gentamicin (100 µg/ml) and, penicillin (250 UI/ml)/streptomycin (250 mg/ml) containing medium after 1 h of infection. For naftifine (Medice) inhibited bacterial infections, bacteria were grown overnight in medium with empirically-optimized concentrations of naftifine (50 µM). The pelleted SA and GBS were characterized for staphyloxanthin biosynthetic pathway inhibition by Raman spectroscopy and used for infection with MOI 10 and MOI 20 respectively. µRNA was isolated from the naftifine-inhibited bacteria and used for stimulations tests. Experiments with small molecule inhibitors: THP1 MΦ were pre-treated with MCC950 (5 µM, 10 µM) (44), Dynasore (150, 100 µM) (Enzo), KCL (60 mM, 45 mM, 75 mM), H-151 (100 µM, 50 µM, 10 µM) (ProbeChem) for 1 h prior to the stimulations. Experiments with RNase A, DNase I: THP1 MΦ were pre-treated cytosolically (+LF) with increasing concentrations of RNase A, DNase I (100 ng/ml, 50 ng/ml, 10 ng/ml) for 2 h washed and then infected with SA and MLBs. Heat inactivated RNase A was used as a control for these experiments. Supernatants were collected after 16 hr for IL-1β production measurement.

### ELISA and LDH

IL-1β and TNF-α production were measured with ELISA (R&D systems) and cell death was measured with LDH (TAKARA clonetech). This was performed in the supernatant collected after 16 h of stimulation. Relative LDH release was calculated as LDH release (% cell death) =100*((measurement-unstimulated cells)/ (lysis control-unstimulated cells)).

### Western blot and STING dimerization assay

STING dimerization was assayed under semi-native conditions. Five hundred thousand THP1 MΦ after stimulation with µRNA of *S. aureus* (SA), Δ*agr* and *in-vitro*-transcribed RNAIII (3 µg ml^-1^, 1 µg ml^-1^) were lysed in 30 µl of 1X cell lysis buffer (Cell Signaling) supplemented with 1X protease cocktail inhibitor (Roche) and 1X sample buffer. Whole cell lysate were sonicated for 7 min with at intensity before loading onto gel without heating. Separation was done using 4-12% SDS-PAGE gel electrophoresis were each gel was run initially for 10 min at 70 V, and then at 120 V for 1 h 30 min.

GSDMD cleavage in human plasma of *S. aureus* sepsis patients and ICU controls was assayed through Western blot. 1 µg plasma with 1X sample buffer was heated at 95°C for 5 min before loading on the gel. Separation was performed by 4-12% SDS-PAGE gel electrophoresis with each gel run at 70 V.

Proteins were blotted onto PVDF membrane, blocked in bovine serum albumin for anti-GSDMD and in skim–milk for anti-STING, respectively with indicated primary and secondary antibodies. Chemiluminescent signals were recorded with a CCD camera. Antibodies used: STING (R&D systems), GAPDH (Thermo Fisher), GSDMD (Proteintech).

### Binding assay for µRNA and CDN

4 µg µRNA or 4 µg Big RNA was prepared in buffer (20 mM tris-HCL pH 8.4, 50 mM KCl 0.1 mM MgCl2) were heated at 80°C for 10 min (reaction A). 16 µM biotin-c-di-AMP and streptavidin-HRP was prepared in buffer (20 mM tris-HCL pH 8.4, 50 mM KCl 0.1 mM MgCl2) and heated at 37°C for 30 min (reaction B). Reaction B was then added to reaction A and stepwise incubated for 37°C for 10 min followed by 30°C for 10 min, 25°C for 10 min and, finally, 4°C for 5 min.

To the above reaction 300 µl cold water and 1.5 V 100% ethanol were added and eluted in 50 µl water through a µRNA extraction column according to the manufacturer’s instructions. Following this, the eluted RNA was diluted with TMB substrate (1:1) in a 96 well plate. The reaction was stopped with 2N H2SO4 and absorbance measured at 450 nm. Diagrammatic representation of the method is shown in **Fig. 4H**. A similar procedure was performed for µRNA from SA and mutant Δ*agr* and *in-vitro-*transcribed RNAIII as depicted in **Fig. 4P**. **LC-MS for binding of µRNA and CDN** 4 µg µRNA of SA, Δ*agr* and *in-vitro*-transcribed RNAIII were prepared in buffer (20 mM tris-HCL pH 8.4, 50 mM KCl 0.1 mM MgCl2) heated at 80°C for 10 min (reaction A). 16 µM biotin c-di-AMP and streptavidin-HRP was prepared in buffer (20 mM tris-HCL pH 8.4, 50 mM KCl 0.1 mM MgCl2) and heated at 37°C for 30 min (reaction B). Reaction B was then added to reaction A and stepwise incubated for 37°C for 10 min then 30°C for 10 min, 25°C for 10 min and, finally, 4°C for 5 min. To the above reaction 300 µl cold water and 1.5 volume of 100% ethanol were added and eluted in 50 µl water through a RNA extraction column according to the manufacturer’s instructions. Following this the eluted RNA was diluted 1:100, 1:50 and 1:10 with RNase free water. The diluted RNA was subjected to ultra-high performance liquid chromatography coupled with high resolution mass spectrometry using THERMO (Bremen, Germany). The detailed procedure of LC-MS is described in the section entitled “LC-MS of c-di-AMP and derivatives in MLBs”. One of the three independent experiments is shown.

### Spinach fluorescence assay

The small molecule binding aptamer used in the study had been previously characterized (62). The spinach DNA primers were amplified by PCR up to 25 cycles with DNA polymerase. The resulting DNA was gel purified confirmed by sequencing and subsequently *in-vitro-* transcribed using T7 RNA polymerase. The RNA was further precipitated and column-purified with warm wash buffer and dissolved in spinach reaction buffer (40 mM HEPES, 125 mM KCL, 3 mM MgCl2). The RNA samples were heated at 70°C for 3 min, and then cooled down at room temperature over 5 min. The concentration of RNA was adjusted to 100 nM for each measurement. The concentration of CDN (c-di-AMP) was 0.1 pM-10 nM and DFHBI (3,5-Difluoro-4-hydroxybenzylidene) was added to a final concentration of 10 µM. Samples were incubated at 37°C in a black 96 well microtiter plate until equilibrium was reached. Measured were then made with a Tecan infinite plate reader at excitation 460 nm and emission 500 nm. Background fluorescence was subtracted and data were normalized. Diagrammatic representation is shown in **Fig. 5G**.

### Confocal microscopy

For visualization of STING and LAMP-1 co-localization following transfection with µRNA (1 µg ml^-1^) from SA, mutant Δ*agr* bacteria and infections with GBS and *S. aureus* wild type, 300,000 cells on coverslips were fixed with ice-cold methanol for 15 min and permeabilized with saponin based permebilization buffer (Invitrogen). Cells were stained with antibodies directed against STING (R&D systems) and LAMP-1 (Abcam). Secondary antibodies included Alexa Flour^®^ 488 and Alexa flour^®^ 647 were used. Images were acquired using a Zeiss LSM 780 confocal microscope using a 63X lens. Quantification of the co-localization (%) was calculated by ((number of co-localized yellow speckles / number of STING (red) speckles) *100).

### Inhibition of staphyloxanthin biosynthetic pathway

For inhibition of staphyloxanthin an overnight culture of GBS was inoculated onto the plate as a lawn culture. After drying, increasing concentrations of naftifine (200 ng/ml, 100 ng/ml, 10 ng/ml) disc filter paper (6 mm diameter, Whatmann filter paper no. 2) were added. A paper disc with PBS was used as a control. The plates were incubated overnight at 37°C and the zone of decolorization analysed.

### Statistical analysis

Data were analysed using Graph Pad Prism software. If indicated, data were analysed for statistically significant differences using paired Students’s *t-*test for two conditions or groups. The quantification of the microscopic STING+LAMP1 structures undertaken using a Mann Whitney test. P values of *< 0.05, **p < 0.01, ***p<0.001,and <0.0001 were considered significant.

## Supporting information

Supplemental data and will be used to link to the file on the preprint site

Supplemental figures and will be used for the link to the file on the preprint site

## Acknowledgements

The authors thank Prof. Viet Hornung for providing some of the CRISPR cell lines used in this study. We thank Prof. Ingrid Hilger for making available the zetasizer nano series instrument. We thank Prof. Vijay A.K. Rathinam and Prof. Reinhard Wetzker for critical assessment of the manuscript. We thank the DFG for funding of the UHPLC-Q-Exactive plus mass spectrometer system within the CRC 1127 (ChemBioSys). This work was mainly supported by the Federal Ministry of Education and Research, Germany (FKZ: 01EO1502, FKZ: 01EO1002) and partly by the DFG-funded Collaborative Research Centre PolyTarget (SFB 1278, Project C03).

## Authors contributions

SND performed or assisted in majority of experiments and analysed data; MG, BG assisted with the experiments and data analysis; SND, SDD wrote the manuscript; MW performed electron microscopy and data analysis; HG, NU and CB performed HPLC/MS/NMR experiments and data analysis; AR performed Raman spectroscopy and data analysis; UN and JP provided resources and technical help in Raman spectroscopy; MB, CS and TB provided human specimens, clinical data; BL and LPNT provided patient isolates, bacterial mutants and clinical data; MB, MS helped in experimental design, clinical data analysis and writing of the manuscript; SDD and MB obtained funding; SDD conceived, designed, and supervised this study, interpreted data, provided guidance with experimental design. All authors critically revised the manuscript for important intellectual content.

## Disclosures

All authors declare no conflict of interest.

## Notes

### Competing Interest Statement

The authors have declared no competing interest.

