## Supplemental data and will be used to link to the file on the preprint site for "Gram-positive bacteria secrete RNA aptamers to activate human STING for IL-1β release"

### SUPPLEMENTARY FIGURE LEGENDS

#### Supplemental Fig. 1: Activation of canonical and non-canonical inflammasome pathway with LPS and RNA (related to Fig. 1)

**(A, B)** THP1 (black), *CASP1^-/-^* (blue), *CASP4^-/-^* (green) MΦs were stimulated with LPS (1 µg), MLB RNA, cytosolic LPS (LPS+LF), cytosolic MLB RNA (5µg/ml) and LPS (RNA+LPS+LF). Cell death and IL-1β were measured.

**(C)** THP1 (black), *CASP4^-/-^* (green), *CASP1^-/-^* (blue), *STING^-/-^* (pattern), *cGAS^-/-^* (blank), *ASC^-/-^* (grey) MΦs were stimulated with nigericin (6.7 µM) IL-1β production was measured.

**(D)** THP1 (black), *cGAS^-/ -^*(white) and *STING^-/-^* (pattern) MΦs were infected with *S. aureus* (MOI 20) and GBS (MOI 10), TNF-α was measured

**(E, F)** Human blood derived MΦs were pretreated with increasing concentrations of STING inhibitor (100 µM, 50 µM, 10µM) for 1 h and then infected with SA (MOI 10) and GBS (MOI 20) respectively. TNF-α production was measured.

#### Supplemental Fig. 2: Characterization of MVs and MLBs (related to Fig. 2)

**(A)** Negative staining TEM image of purified outer membrane vesicles (OMV) from *E.coli*. The bar represents 100 nm.

**(B)** TNF-α production was measured in THP1 MΦs following stimulation with MLBs and MVs isolated from SA.

**(C)** Lysyl-PG mass spectra detected in (a) vesicle mixture (MLBs+MVs). (b) Membrane vesicles (MVs). (c) Multi lamellar vesicular bodies (MLBs).

**(D)** Comparative LCMS analysis on the presence of 4,4’-diaponeurosporenoic acid (St1) showing in EIC chromatogram of m/z 433.3 at t_R_=7.625 min (a) SA lipid extract (b) MeOH extract of vesicle mixture (MLBs+MVs) (c) MeoH extract of multi lamellar vesicular bodies (MLBs) (d) MeOH extract of membrane vesicles (MVs).

**(E)** Mass spectra of positive control c-di-AMP (blue) (I) & pApA (green) (II) mass spectra of c-di-AMP and its derivative pApA in *S. aureus* MLBs (III).

**(F)** Negative staining TEM image of purified GBS MLBs when subjected to higher temperature. Bar represents 100 nm.

**(G)** DLS measurement of MLBs showing size with increasing temperature (15°C-50°C).

#### Supplemental Fig. 3: Activation of canonical inflammasome pathway with nigericin (related to Fig. 3)

**(A)** THP1 MΦs were pre-treated with increasing concentrations of KCL (60 mM, 45 mM, 75 µM) and stimulated with nigericin (6.7 µM). IL-1β was measured.

**(B)** THP1 (black), *STING^-/-^* (grey) MΦs treated with increasing concentrations (10 µM-60 µM) of nigericin. IL-1β was measured.

**(C)** IL-1β production measured when THP1 MΦ were pre-treated with indicated inhibitor for 1 h and stimulated with OMVs.

#### Supplemental Fig. 4: MLB µRNA activates inflammasome via STING (related to Fig. 4)

**(A)** THP1 (black), *STING^-/-^* (pattern), *cGAS^-/-^* (white) MΦs were stimulated with GBS MLBs µRNA (µRNA) (5 µg/ml), cytosolic µRNA (µRNA+LF) and IL-1β production was measured.

**(B)** Mass spectra of c-di-AMP detected in I) µRNA of *S. aureus* (SA), *∆agr*. II) Different concentration of SA µRNA bound CDN fractions. III) Different concentration of ∆*agr* µRNA bound CDN fractions.

**(C)** IL-1β production was measured in THP1 (black) MΦs when stimulated with µRNA from *S. aureus* (SA) and *∆agr* mutant cytosolically (SA+LF, *∆agr*+LF) and with CDN (SA_CDN_+LF, *∆agr*_CDN_+LF), LF.

#### Supplemental Fig. 5: RNAIII and Differentially abundant µRNA in MLB (related to Fig. 5)

**(A)** Venn diagram obtained through deep sequencing in SA (blue), SA-MLB (white) and SA-∆*agr* (green), summarizing the overlap between differentially abundant reads mapped on *S. aureus* subsp. *aureus* N315 (accession number: NC_002745.2) corresponding to 186 differentially abundant genomic regions.

**(B)** Comparative percent abundance sequencing read profiles of different sRNA quantified from *S. aureus* MLB (SA-MLB) (white), *S. aureus* (SA) (blue), and *S. aureus* ∆*agr* (SA-∆*agr*) (green).

**(C)** Comparative percent abundance sequencing read profile of sbr-C, Rsa C, RNAIII and WAN01CC66-rc in wild type *S. aureus* (SA) (Blue) and *S. aureus* MLBs (SA-MLB) (white).

**(D)** Visualization of read coverage of genomic locations of Rsa C (upper), RNAIII (middle) and WAN01CC66-rc (lower) identified by deep sequencing on wild type *S. aureus* (SA), *S. aureus* ∆*agr* (SA-∆*agr*) and *S. aureus* MLBs (SA-MLB). The reads of each gene are shown in red colour.

**(E)** Mass spectra of c-di-AMP detected in different concentration of c-di-AMP bound RNAIII fractions.

**(F)** Visualization of differentially abundant sequencing reads of SA (blue) and SA-MLB (pink) and overlaid (maroon) mapped on 514 base RNAIII cloned genomic locus of *S. aureus* (LS1).

**(G)** IL-1β production measured in THP1 (black) MΦs following stimulations with *in-vitro*-transcribed RNAIII from 1-380 nt, 202-514 nt (5 µg/ml, 2.5 µg/ml) on surface and into the cytosol (1-380+LF, 202-514+LF) and LF.

**(H)** Secondary structure of RNAIII and different stretches of RNAIII from 1-380 nt, 202-514 nt, 202-317 nt, 213-514 nt and 1-213 nt. Central domain A,7,8,9,A in the structures is denoted in red colour. Structures were obtained using RNA fold web server at Vienna RNA Package (Gruber, Lorenz et al., 2008).

**(I)** THP1 MΦs (black) were stimulated with *in-vitro*-transcribed RNAIII from 1-514 nt, 1-213 and 213-514 nt (5 µg/ml, 2.5 µg/ml) on surface and into cytosol (1-514+LF, 1-213+LF, 213-514+LF) and LF. IL-1β production measured.

**(J)** Control THP1 MΦs incubated with µRNA from (SA), ∆*agr* and *in-vitro*-transcribed RNAIII (3 µg/ml, 1 µg/ml) were used in STING dimerization assay followed by semi-native gel electrophoresis and resolution of STING monomer (37 kDa) and dimer (72 kDa) were observed. GAPDH was used as a loading control.

**K (i))** THP1 MΦs were transfected with µRNA (1 µg/ml) from SA, ∆*agr* for 3 h. The cells were fixed and stained with anti-STING (red) and anti-LAMP-1 (green) specific antibodies. Percent co-localization of colocalized STING and LAMP1 was measured.

**(K (ii))** THP1 MΦs were infected with GBS (MOI 20) and SA (MOI 10) for 3 h. The cells were fixed and stained with anti-STING (red) and anti-LAMP-1 (green) specific antibodies. Percent co-localization of colocalized STING and LAMP1 was measured.

Data shown are mean ±SD (n=3), representative of at least three independent experiments. Asterisks indicate statistically significant differences (∗p < 0.05, ∗∗p < 0.01 and ∗∗∗p < 0.001).

**(L)** Overview of the inflammasome activation via Gram-positive bacterial RNA aptamers.

#### Supplemental Fig. 6: Staphyloxanthin biosynthesis pathway (related to Fig. 6)

**(A)** Bacterial biosynthetic pathway showing staphyloxanthin production in *S. aureus*.

#### Supplemental Fig. 7: Characterization of 4,4´-diaponeurosporenoic acid (St1) and staphyloxanthin (St2). (related to Fig. 2 and 6)

**(A)** UV spectra showing maximum absorption wavelenght of 4,4´-diaponeurosporenoic acid (St1).

**(B)** UV spectra showing maximum absorption wavelength of staphyloxanthin (St2).

**(C)** ESI- MS spectra of 4,4´-diaponeurosporenoic acid (St1) in a positive ion mode ([M+H]^+^ represents the protonated molecular ion).

**(D)** ESI- MS spectra of staphyloxanthin (St2) in a positive ion mode.

**(E)** ESI- MS spectra of staphyloxanthin (St2) negative ion mode.

([M]^+^• represents the molecular ion due to the high polyene conjugation; [M+Na]^+^ represent the sodium adduct ion; [M-H]- represents the deprotonated molecular ion; [M-H+FA]^-^ represents the deprotonated formic acid adduct ion.)

**(F)** ^1^HNMR spectrum (600 MHz, CDCl_3_) of staphyloxanthin (St2).

**(G)** ^1^H-^1^H COSY spectrum (600 MHz, CDCl_3_) of staphyloxanthin (St2), showing the coupled spins between proton.

**(H)** HSQC spectrum (600 MHz, CDCl_3_) of staphyloxanthin (St2), showing the proton-carbon single bond correlation.

**(I)** HMBC spectrum (600 MHz, CDCl3) of staphyloxanthin (St2), showing the correlations between carbons and protons that are separated by two-, three- or four bond.

**(J)** Structures of 4,4’-diaponeurosporenoic acid (St1) and **(K)** staphyloxanthin (St2).

### SUPPLEMENTARY TABLES

#### Supplemental Table 1: Raman vibrational assignments of MLB and staphyloxanthin (St2) and 4,4’-diaponeurosporenoic acid (St1)

| MLB | St1 | St2 | Vibrational assignments and reference |
| --- | --- | --- | --- |
| 706 |  |  | Ring breathing of ester like compounds (78) |
| 787 |  |  | O-P-O of nucleic acids (79) |
| 839 |  |  | C-O-C stretching of long chain lipids, fatty acids (80) |
| 971 | 975 | 975 | Phosphates monoester group (81, 80, 85) |
| 984 |  |  | C-C stretching of proteins (83) |
| 1000 |  |  | C-C aromatic ring stretching |
| 1005 |  |  | Phenylalanine (proteins) |
| 1009 | 1014 | 1014 | Aromatic ring C-O stretching, C-C stretching, O-C-H deformation (80) |
| 1100 |  |  | C-C stretching (82) |
| 1163 | 1168 | 1168 | C-C stretching of unsaturated compounds (80, 82) |
|  | 1206 | 1210 | CH_2_ deformation vibrations |
| 1239 |  |  | Amide III C-N stretching |
|  | 1298 | 1294 | Deformation vibration of ester and saturated fatty acid chains |
| 1335 |  |  | CH_3_CH_2_ deformation and nucleic acid vibrations (80) |
| 1345 |  |  | C-H deformation vibration |
| 1456 | 1451 | 1451 | CH_2_/CH_3_ deformation of lipids, proteins and carbohydrates (82) |
| 1459 |  |  | CH_2_ deformation of deoxyribose |
| 1528 | 1528 | 1528 | C=C stretching of unsaturated compounds (80) |
| 1580 |  | 1581 | Nucleic acid vibrations/asymetric stretching(COO−) (79) |
| 1661 |  |  | Amide I C=O stretching |
| 1669 |  |  | C=C stretching of Cholesterol ester (80, 82) |
| 1732 |  |  | C=O stretching of long chain lipids(unsaturated) (81) |
| 2856 | 2856 | 2856 | Symmetric stretching of CH_2_ lipids and fatty acids (82) |
| 2890 | 2890 | 2886 | Symmetric stretching of CH_3_ lipids and fatty acids (82) |
| 2907 |  |  | CH_3_ stretching |
| 2934 |  |  | C-H stretching |
| 2967 |  |  | Out-of-plane long chain end asymmetric CH_3_ stretching |
| 2974 |  |  | Asymmetric CH3 stretching of lipids, fatty acids cholesterol ester (82, 86) |
| 3041 |  |  | C-H stretching aromatic ring compounds (82) |
| 3067 |  |  | C-H stretching aromatic ring compounds (82) |

#### Supplemental Table 2: Characteristic fingerprint Raman vibrations of MLB and staphyloxanthin (St2) and 4,4´-diaponeurosporenoic acid (St1) from *S. aureus*

| Sample | Fingerprint Raman Vibrations cm-1 | | | |
| --- | --- | --- | --- | --- |
| *S. aureus* MLB | 1009 | 1163 | 1528 | **Fig. 2 N, 2O, 2P** |
| (St1) | 1013 | 1167 | 1528 | **Fig. 7A** |
| (St2) | 1014 | 1168 | 1528 | **Fig.7B** |

#### Supplemental Table 3: Gradient for UHPLC / HRMS measurement.

| **Time**  **[min]** | **Solvent B**  **[%]** |
| --- | --- |
| 0 | 0 |
| 0.2 | 0 |
| 8.0 | 100 |
| 11.0 | 100 |
| 11.1 | 0 |
| 12.0 | 0 |

| Supplemental Table 4: Differentially abundant sRNA species identified in our study | **sRNA Genes** | | **Start-end (size)** | **Flanking ends** | **Strand** | **Peak Height** | | |  | **Study** | |
| --- | --- | --- | --- | --- | --- | --- | --- | --- | --- | --- | --- |
|  |  |  |  |  |  | **∆*agr*** | **SA** | **SA MLBs** |  |  | |
| ***rsa* genes** | Rsa A | | 637111-637250  (139) | SA0543/SA0544 | \| <>< | 178,731 |  |  | 637112-637256 | **52** | |
|  | RsaB | | 11778009–1778068 (60) | SA1552/*fhs* | <>< | ND | ND | ND |  |  |  |
|  | Rsa C | | 679909-680452  (554) | SA0586/SA0587 | <>> |  |  | 74,703 | 679912-680136 |  |  |
|  |  |  |  |  |  |  |  | 162,989 | 680286-680594 |  |  |
|  |  |  |  |  |  |  | 87,701 |  | 680489-680602 |  |  |
|  | Rsa D | | 695867-696043  (177) | SA06007sa0601 | ><> | ND | ND | ND |  |  |  |
|  | Rsa E | | 975383-975482  (100) | SA0859/SA0860 | >>< | 268,097 | 684,07 | 74,703 | 975382-975493 |  |  |
|  | RsaF | | 975461–975564  (104) | SA0859/SA0860 | >>< | 268,097 | 684,067 | 74,703 | 975382-975493 |  |  |
|  | RsaG | | 254702–254895  (194) | *uhp*T/SA0215 | >>< | ND | ND | ND |  |  |  |
|  | RsaH | | 829511–829634  (124) | SA0724/SA0725 | <>> | 983,021 | 1490,914 | 129,033 | 829508-829675 |  |  |
|  | RsaI | | 2367918–2368061 (144) | SA2104/SA2105 | <<> | 625,559 | 1219,042 | 339,561 | 2367919-2368077 |  |  |
|  | RsaJ | | 2486806–2487092 (287) | *bio*D/SA2216 | <>< | ND | ND | ND |  |  |  |
|  | RsaK | | 216920–217128  (209) | *glc*A/SA0184 | >>< | ND | ND | ND |  |  |  |
|  | RsaX01 | | 43000–43600 | SA0035/SA0036 |  | ND | ND | ND |  |  |  |
|  | RsaX02 | | 81706–81855 | kdpC/SA0072 |  | ND | ND | ND |  |  |  |
|  | RsaX03 | | 95295–95689 | SA0084–SA0085 |  | ND | ND | ND |  |  |  |
|  | RsaX04 | | 149756-149850 | SA0129–SA0130 |  | ND | ND | ND |  |  |  |
|  | RsaX05 | | 470600-470900 | SA0410/ndhF |  | ND | ND | ND |  |  |  |
|  | RsaX06 | | 659823-659968 | SA0565/SA0566 |  | ND | ND | ND |  |  |  |
|  | RsaX07 | | 666122-666257 | SA0575/sarA |  | ND | ND | ND |  |  |  |
|  | RsaX08 | | 761087-761230 | SA0667/SA0668 |  | ND | ND | ND |  |  |  |
|  | RsaX09 | | 905650-905835 | SA0801/SA0802 |  | ND | ND | ND |  |  |  |
|  | RsaX10 | | 1138640-1138848 | SA1005/SA1006 |  | ND | ND | ND |  |  |  |
|  | RsaX11 | | 1175489-1175753 | SA1037/lsp HMM |  | ND | ND | ND |  |  |  |
|  | RsaX12 | | 1399044-1399209 | SA1224/lysC |  | 102,132 |  |  | 1381953-1382011 |  |  |
|  | RsaX13 | | 1421983-1422128 | odhA/arlS |  | ND | ND | ND |  |  |  |
|  | RsaX14 | | 1801062-1801380 | SA1570/SA1571 |  | ND | ND | ND |  |  |  |
|  | RsaX15 | | 1839300-1839600 | SA1602/trunc.SA |  | ND | ND | ND |  |  |  |
|  | RsaX16 | | 1914236-1914464 | SA1676/tnp |  | ND | ND | ND |  |  |  |
|  | RsaX17 | | 1992090-1992414 | SA1738/SAS056 |  |  | 149,091 |  | 1992614-1992746 |  |  |
|  | RsaX18 | | 2436200-2436800 | SA2168–SA2169 |  | 765,99 | 1280,432 | 190,154 | 2435837-2436684 |  |  |
|  |  |  |  |  |  |  |  | 196,945 | 2436600-2436872 |  |  |
|  | RsaX19 | | 2440668-2440850 | scrA/gltT |  | ND | ND | ND |  |  |  |
|  | RsaX20 | | 2473450-2473566 | SA2203/SA2204 |  | ND | ND | ND |  |  |  |
|  | RsaX21 | | 2544796-2545113 | SA2268/SA2269 |  | 2578,833 | 3244,931 | 543,297 | 2544417-2544836 |  |  |
|  |  |  |  |  |  |  |  | 54,33 | 2544740-2544833 |  |  |
|  | RsaX22 | | 2586551-2586821 | SA2301/SA2302 |  | ND | ND | ND |  |  |  |
|  | RsaX23 | | 2590928-2591046 | fbp/SA2305 |  | ND | ND | ND |  |  |  |
|  | RsaX24 | | 2649194-2649400 | isaA/SA2357 |  | ND | ND | ND |  |  |  |
|  | RsaX25 | | 2769241-2769725 | SA2457/icaR |  | ND | ND | ND |  |  |  |
| ***sau*** **genes** | Sau-02 | | 1006365-1006481  (117) |  | < | 268,097 | 929,629 |  | 1006365-1006481 | | **50** |
|  | Sau-13 | | 2715794-2715897  (104) |  | < | ND | ND | ND |  | |  |
|  | Sau-19 | | 2544997:2545071  (75) |  | > | ND | ND | ND |  | |  |
|  | Sau-24 | | 1225832:1225902  (70) |  | > | ND | ND | ND |  | |  |
|  | Sau-27 | | 1283539-1283629  (90) |  | > | ND | ND | ND |  | |  |
|  | Sau-30 | | 2294537-2294600  (64) |  | < | ND | ND | ND |  | |  |
|  | Sau-31 | | 2293495:2293560  (66) |  | > | ND | ND | ND |  | |  |
|  | Sau-41 | | 469933:470088  (156) |  | < | ND | ND | ND |  | |  |
|  | Sau-50 | | 1510745:1510892  (147) |  | > | ND | 87,701 | ND | 1510745-1510892 | |  |
|  | Sau-53 | | 208728:208919  (192) |  | > | ND | ND | ND |  | |  |
|  | Sau-59 | | 1057770:1057897  (127) |  | < | ND | ND | ND |  | |  |
|  | Sau-63 | 436958:437055  (98) | |  | < | 702,158 | 833,158 | ND | 436958-437055 | |  |
|  | Sau-64 | 637087:637253  (167) | |  | > | 178,731 | ND | ND | 637087-637253 | |  |
|  | Sau-66 | 763840:763919  (80) | |  | < | ND | ND | ND |  | |  |
|  | Sau-5949 | 1799353:1799411  (60) | |  | > | 1263,884 | 2288,992 | ND | 1799353-1799411 | |  |
|  | Sau-5971 | 413160:413255  (96) | |  | < | 612,792 | 2078,51 | ND | 413160-413255 | |  |
|  | Sau-6053 | 995878:996044  (170) | |  | > | ND | 210,482 | ND | 995878-996044 | |  |
|  | Sau-6072 | 2616272:2616324  (50) | |  | > | ND | ND | ND |  | |  |
| ***spr* genes** | *sprA* | 2006878-2007561  (150) | |  | < | ND | ND | ND |  | | **51** |
|  | *sprB* | 2008571/2009086  (280) | |  | > | ND | ND | ND |  | |  |
|  | *sprC* | 2010789/2011001  (140) | |  | > | ND | ND | ND |  | |  |
|  | *sprD* | 2010789/2011001  (90) | |  | < | ND | ND | ND |  | |  |
| ***WAN* genes** | WAN01CCBZ(IGR1520) |  | | SA1623-SA1624 | > | ND | 228,022 | ND | 1856485-1856726 | |  |
|  | WAN01CCFGrc(IGR1644) |  | | SA1761-SA1760 | > | ND | ND | ND |  | |  |
|  | WAN01BTFS |  | | SAP28-SAP29 | < | ND | ND | ND |  | |  |
|  | WAN01CBQ |  | | SA0434-SA0435 | > | ND | ND | ND |  | |  |
|  | WAN01CC8T |  | | SA1456-SA1455 | > | 1621,346 | 1385,673 | 2906,641 |  | |  |
|  | WAN014I74 |  | | 23srRNA-SA1968 | > | 33307,801 | 16259,734 | 52068,266 |  | |  |
|  | WAN01CC66-rc (RNaseP) |  | | SA1278-SA1277 | > |  |  | 54,33 | 1484043-1484120 | |  |
|  |  |  |  |  |  |  | 140,321 | 196,945 | 1483744-1484002 | |  |
|  | WAN014FZW |  | | SA2218-SA2219 | < | 1 | 1 | 1 |  | |  |
| ***rsa* O genes** | *rsa* OA |  | | SA0084-SA0085 | < | ND | ND | ND |  | | **53, 54** |
|  | *rsa* OB |  | | SA0347-SA0348 | < | ND | ND | ND |  | |  |
|  | *rsa* OC |  | | SA0667-SA0668 | < | ND | ND | ND |  | |  |
|  | *rsa* OD |  | | SA01393-SA1394 | > | 114,899 | 333,263 | ND | 1600191-1600375 | |  |
|  | *rsa* OE |  | | SA1579-Sa1580 | < | ND | ND | ND |  | |  |
|  | *rsa* OF |  | | SA1704-SA1705 | < | ND | ND | ND |  | |  |
|  | *rsa* OG |  | | SA2104-SA2105 | < | 625,559 | 1219,042 | 339,561 | 2367919-2368077 | |  |
|  | *rsa* OI |  | | *pro*P-SA0532 | < | ND | 271,873 | ND | 626797-627130 | |  |
|  | *rsa* OL |  | | *Csp* C-SA 0748 | < | ND | 210,482 | ND | 856954-857042 | |  |
|  |  |  |  |  |  | 817,056 | 1973,269 | ND | 857197-857300 | |  |
|  | *rsa* OM |  | | SAS023-SA0769 | > | ND | ND | 1 | 875349-875597 | |  |
|  | *rsa* OO |  | | SA0891-SA0892 | < | ND | ND | ND |  | |  |
|  | *rsa* OP |  | | SA0961-SA0962 | > | ND | 131,551 | ND | 1089149-108566 | |  |
|  | *rsa* OQ |  | | SA1198-SA1199 | > | ND | ND | ND |  | |  |
|  | *rsa* OR |  | | SA1756/57-*sak* | < | ND | ND | ND |  | |  |
|  | *rsa* OG |  | | SA2104/SA2105 | < | 625,559 | 1219,042 | 339,561 | 2367825-2368208 | |  |
|  | *rsa* OT |  | | SA2267-SA2268 | < | 2578,833 | 3244,931 | 543,297 | 2544336-2544594 | |  |
|  | *rsa* OU |  | | *pts*G-SA2327 | < | 1 | 1 | ND | 2611619-2611918 | |  |
|  | *rsa* OV |  | | SA2343-*copA* | > | ND | 122,781 | ND | 2632123-2632361 | |  |
| ***sbr* genes** | *sbr A* |  | | SA0891-SA0892 | < | ND | ND | ND |  | | **55** |
|  | *sbr B* |  | | SA1173-SA1174 | > | 204,264 | 140,321 | ND | 1336856-1336917 | |  |
|  | *sbr C* |  | | SA0586-SA0587 | > | ND | ND | 74,703 | 679912-680136 | |  |
|  |  |  |  |  |  | ND | ND | 162,989 | 680286-680594 | |  |
|  |  |  |  |  |  | ND | 87,701 | ND | 680489-680602 | |  |
|  | *sbr D* |  | | SA0040-SA0041 | < | 1 | 1 | ND |  | |  |
|  | *sbr E* |  | | SA1519-SA1520 | > | ND | ND | ND |  | |  |
| ***Teg* genes** | *Teg 1* |  | | SAS001-SA0011 | > | ND | ND | ND |  | | **54** |
|  | *Teg 76* |  | | SA0372-xprT | > | ND | ND | ND |  | |  |
|  | *Teg 35* |  | | hutH -serS | > | ND | ND | ND |  | |  |
|  | *Teg 38* |  | | SA0184-glcA | < | 1 | 1 | ND | 217075-216958 | |  |
|  | *Teg 45* |  | | gltX -cysE | > | ND | ND | ND |  | |  |
|  | *Teg 56* |  | | SA0984-pheS | > | ND | ND | ND |  | |  |
|  | *Teg 69* |  | | lytH- hisS | < |  | 184,172 |  | 1664242-1663989 | |  |
|  | *Teg 70* |  | | Tag-valS | < |  | 114,011 |  | 1696594-1696647 | |  |
|  | *Teg 73* |  | | SA1580-leuS | < | ND | ND | 1 | 1818801-1818658 | |  |
|  | *Teg 4* |  | | xylR-mecI | < | ND | ND | ND |  | |  |
|  | *Teg 19b* |  | | SAS069-truncatedSA | < | ND | 114,011 | 54,33 | 2206705-2206453 | |  |
|  | *Teg 24* |  | | SA2105-SA2104 | < | 625,559 | 1219,042 | 339,561 | 2368050-2367940 | |  |
|  | *Teg 42* |  | | SA0434-SA0435 | > | 4659,773 | 6288,15 | 2906,641 | 501430-501682 | |  |
|  | *Teg 47* |  | | SA0532-proP | < | 1 | 271,873 | 1 | 627227-626915 | |  |
|  | *Teg 60* |  | | SA1015-SA1016 | > | ND | ND | ND |  | |  |
|  | *Teg 91* |  | | SA0601-SA0600 | < | ND | ND | ND |  | |  |
|  | *Teg 21* |  | | glmM / fmtB | < | ND | ND | ND |  | |  |
|  | *Teg 26* |  | | SA2159 / SA2158 | < | ND | ND | ND |  | |  |
|  | *Teg 28* |  | | SA2238 / opuCA | < | ND | ND | ND |  | |  |
|  | *Teg 55* |  | | SA0962 / pycA | > | ND | ND | ND |  | |  |
|  | *Teg 57* |  | | uvrC / sdhC | > | ND | ND | 1 | 1127112-1127350 | |  |
|  | *Teg 61* |  | | codY / rpsB | > | ND | 1 | ND | 1245093-1245202 | |  |
|  | *Teg 72* |  | | phoR / SA1514 | < | ND | ND | ND |  | |  |
|  | 4.5S RNA | 501358-502001 | |  | > | 4659,773 | 6288,15 | 2906,641 | 501419-501712 | | **51** |
|  | tmRNA | 843706-844543 | |  | > | 3217,158 | 3972,848 | 4122,269 | 843796-844174 | |  |
|  | 6S RNA | 1660650-1660796 | |  | < | 1621,346 | 1385,673 | 2906,641 | 1660495-1660733 | |  |
|  | RNAIII | 2078477-2079445 | | SA10560-SA10570 | < | ND | 192,942 | ND | 2078724-2078847 | | **51** |
|  |  |  |  |  |  | ND | ND | 129,033 | 2078765-2079023 | |  |
|  |  |  |  |  |  | ND | 175,402 | 54,33 | 2079079-2079175 | |  |

#### Supplemental Table 5: Sequences of primers used in the study.

The T7 polymerase promoter sequence is italicized and the restriction enzyme sites are underlined.

| RNAIII(cloning in pET28a) | F: GCGGAATTCAGATCACAGAGATGTGAT |
| --- | --- |
|  | R: GCGAAGCTTAAGGCCGCGAGCTTGGGA |
| RNAIII (T7)(IVT) | F: *TAATACGACTCACTATAGGG*AGATCACAGAGATGTGAT |
|  | R: AAGGCCGCGAGCTTGGGA |
| 1-213 RNAIII(IVT) | F: *TAATACGACTCACTATAGGG*AGATCACAGAGATGTGAT |
|  | R: ACTATACGAAGATAACAAAT |
| 213-514 RNAIII(IVT) | F:*TAATACGACTCACTATAGGG*ACTAAAAGTATGAGTTATTAA |
|  | R: AAGGCCGCGAGCTTGGGA |
| 202-514 RNAIII(IVT) | F: *TAATACGACTCACTATAGGG*CTTCGTATAGTACTAAAAGT |
|  | R: AAGGCCGCGAGCTTGGGA |
| 202-317 RNAIII(IVT) | F: *TAATACGACTCACTATAGGG*CTTCGTATAGTACTAAAAGT |
|  | R: ATAGCACTGAGTCCAAGGA |
| Rsa C(IVT) | F: *TAATACGACTCACTATAGGG*GTTTACTTTGATAGGCCAGA |
|  | R: TCCCATATCGTGCGTTAAATA |
| WAN01CC66-rc (IVT) | F: *TAATACGACTCACTATAGGG*AAAAGCTCTCCATCATCTA |
|  | R: AGTGATATTTTGGGTAATCG |
| HP(IVT) | F: *TAATACGACTCACTATAGGG*TGGTAACCGCACTCGTA |
|  | R: AAACAAAATCATCTTAGCGT |
| Spinach | F: *TAATACGACTCACTATAGGG*ACGCGACTGAATGAAATGG  TGAAGGACGGGTCCACTTCGTATAGTACTAAAAGTATGAGTT  ATTAAGCCATCCCAACTTAATAACCATGTAAAATTAGCA |
|  | R: GACGCGACTAGTTACGGAGCTCACACTCTACTCAACAAATAGC  ACTGAGTCCAAGGAAACTAACTCTACTAGCAAATGTTACT  CACTTGCTAATTTTACATGGTTATTAAGTT |

| Age [years] | 67 | [4.0 - 90.0] |
| --- | --- | --- |
| Gender, male | 32 | (64 %) |
| **Site of infection** | | |
| Blood stream | 14 | (28 %) |
| UTI | 3 | (6 %) |
| Respiratory | 4 | (8 %) |
| Soft tissue | 15 | (30 %) |
| Unknown | 14 | (28 %) |
| **Origin of Bacteria** | | |
| Peripheral blood culture | 40 | (80 %) |
| Central venous blood culture | 7 | (14 %) |
| other | 3 | (6 %) |
| **Referral Department** | | |
| Cardiac | 11 | (22 %) |
| ENT | 1 | (2 %) |
| Haematology / Oncology | 8 | (16 %) |
| Nephrology | 3 | (6 %) |
| Gastro intestinal | 3 | (6 %) |
| Urology | 3 | (6 %) |
| Pneumology | 5 | (10 %) |
| Spine/cerebral | 4 | (8 %) |
| Trauma /Ortho | 5 | (10 %) |
| Skin/Soft Tissue | 4 | (8 %) |
| Unknown | 3 | (6 %) |

#### Supplemental Table 6: Patient characteristics (Bacterial isolates)

#### Supplemental Table 7: Patient characteristics (*S. aureus* sepsis patients)

| Patient No | Age  (years) | Gender | 28 day  mortality | APACHE-II  on admission | SAPS-II  on admission | Site of infection | SOFA |
| --- | --- | --- | --- | --- | --- | --- | --- |
| 1 | 53 | F | Survived | 23 | 49 | Endocarditis | 13 |
| 2 | 71 | M | Survived | 33 | 71 | Abdomen | 14 |
| 3 | 74 | M | Died | 26 | 70 | Wound | 11 |
| 4 | 60 | M | Survived | 10 | 23 | Catheter | 5 |

ICU intensive care unit, APACHE II Acute Physiology and Chronic Health Evaluation II, SAPS II Simplified Acute Physiology Score II

#### Supplemental Table 8: NMR Data (CDCl_3_, at 300 K) for staphyloxanthin (St2).^a^

| **Position** | **δC, mult.b** | **Staphyloxanthin (St2)** | | |
| --- | --- | --- | --- | --- |
|  |  | **δH, mult. (J in Hz)** | **COSY** | **HMBC** |
| 1 | 166.5, qC |  |  |  |
| 2 | 131.2, qC |  |  |  |
| 3 | 141.6, CH | 7.33, br d (11.6) | 4 | 1 |
| 4 | 122.5, CH | 6.47, dd (15.5, 11.6) | 3, |  |
| 5 | 131.7, CH | 6.66, br d (15.5) |  |  |
| 6^c^ | n.d. |  |  |  |
| 7 | 145.89, CH | 6.68, m |  |  |
| 8 | 125.48, CH | 6.52, m |  |  |
| 9 | 134.49, CH | 6.32, m |  |  |
| 10 | n.d. |  |  |  |
| 11 | 140.47, CH | 6.48, m |  |  |
| 12 | 129.08, CH | 6.61, m |  |  |
| 13 | 131.39, CH | 6.21, m |  |  |
| 14 | 126.00, CH | 5.97, m |  |  |
| 15 | n.d. |  |  |  |
| 16 | 136.94, CH | 6.38, m |  |  |
| 17 | 124.25, CH | 6.61, m |  |  |
| 18 | 135.00, CH | 6.24, m |  |  |
| 19 | n.d. |  |  |  |
| 20 | 39.8, CH_2_ | 1.98, m |  |  |
| 21 | 26.8, CH | 2.06, t (7.6) | 22 | 20, 22, 23 |
| 22 | 124.2, CH | 5.11, m | 21, 24 |  |
| 23 | 134.7, qC |  |  |  |
| 24 | 27.6, CH_3_ | 1.58, s |  |  |
| 25 | 12.8, CH_3_ | 1.96-1.98, s |  |  |
| 26 | 12.8, CH_3_ | 1.96-1.98, s |  |  |
| 27 | 12.8, CH_3_ | 1.96-1.98, s |  |  |
| 28 | 12.8, CH_3_ | 1.96-1.98, s |  |  |
| 29 | 12.8, CH_3_ | 1.96-1.98, s |  |  |
| 30 | 27.5, CH_3_ | 1.60, s | 20 | 20, 22, 23 |
| 1’ | 92.1, CH | 5.75, d, (7.4) | 2’ | 1, 3’ |
| 2’ | 72.6, CH | 5.02, t, (7.4) | 1’, 3’ |  |
| 3’ | 75.4, CH | 3.75, m | 2’, 4’ |  |
| 4’ | 70.9, CH | 3.74, m | 3’, 5’ | 3’ |
| 5’ | 76.0, CH | 3.57, m | 4’, 6’b |  |
| 6’a | 62.0, CH_2_ | 3.95, d (12.0) |  |  |
| 6’b |  | 3.85, d (12.0) | 5’ |  |
| 1’’ | 174.0, qC |  |  |  |
| 2’’ | 34.4, CH_2_ | 2.34, m |  | 1’’ |
| 3’’ | 27.5, CH_2_ | 1.57, m | 2’’ | 1’’ |
| 4’’ ‒ 13’’ | 29.4, CH_2_ | 1.25, m |  |  |
| 14’’ | 13.9, CH_3_ | 0.88 |  |  |
| 15’’ | 20.3, CH_3_ | 0.86 |  |  |

^a^-600 MHz for ^1^H NMR and 150 MHz for ^13^C NMR
^b^-numbers of attached protons were determined by analysis of 2D spectra.
^c^-underline: the assignment of NMR resonance is ambiguous due to the lack of HMBC correlations and too weak ^13^C resonances.

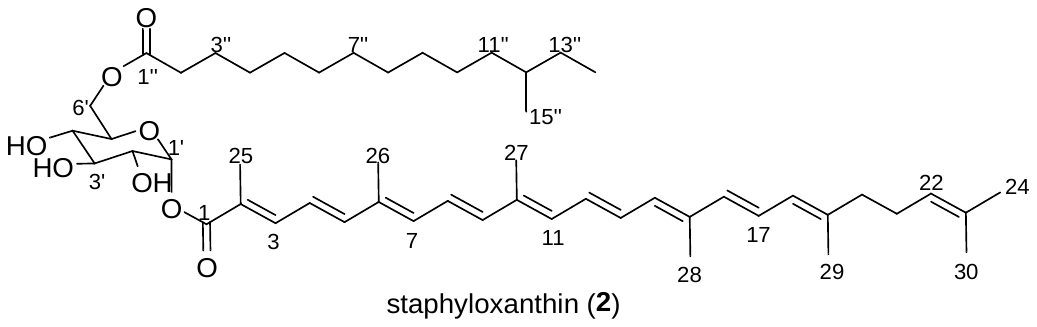

#### SUPPLEMENTARY RESULTS

#### Supplemental Results SR1

**Elucidation of 4,4’-diaponeurosporenoic acid (St1) and staphyloxanthine (St2)**

The presence of 4,4’-diaponeurosporenoic acid (St1) was confirmed by detailed analysis of UV-Vis, MS and NMR spectra. The maximum UV absorption of compound St1 (λmax 455 nm) indicated the carotenoid feature. The corresponding molecule ion at *m*/*z* 432.3020 ([M]^+•^ Δ = ‒ 0.21 ppm) deduced the molecular formula of C_30_H_40_O_2_. The ^1^H NMR spectrum of 4,4’-diaponeurosporenoic acid (St1) containing fraction indicated the signals from δ_H_ 6.2–6.6 ppm belong to the conjugated double bond from carotenoid moiety.

The structure of staphyloxanthin (St2) was assigned according to the detailed analysis of UV-Vis, MS and NMR spectra. The maximum absorption in the UV spectra at 464 nm of staphyloxanthin indicated the carotenoid feature **(Supplemental Fig. 7)**. The corresponding molecule ion at *m*/*z* 818.5674 ([M]^+•^ Δ = ‒ 2.13 ppm) deduced the molecular formula of C_51_H_78_O_8_. This conclusion was confirmed by the observation of corresponding sodium adduct at *m*/*z* 841.70 ([M+Na]^+^) in the positive mode and deprotonated ion at m/z 817.40 ([M–H]^–^) and formic acid adduct at *m*/*z* 863.75 ([M–H+FA]^–^) in Shimadzu low resolution mass spectrometer **(Supplemental Fig. 9)**. After detailed analysis of ^1^D and ^2^D NMR spectra of staphyloxanthin, the olefinic protons from δ_H_ 6.2–6.6 ppm (δ_C_ 122.5–145.9 ppm), methylene group at δ_H_ 1.98 ppm/δ_C_ 39.8 ppm and δ_H_ 2.07 ppm/δ_C_ 26.9 ppm and methyl groups at δ_H_ 1.68 ppm/δ_C_ 25.8 ppm and δ_H_ 1.97 ppm/δ_C_ 12.9 ppm suggested the presence of isoprene moiety belong to 4,4’-diaponeurosporenoic acid. This conclusion was further confirmed by the coupling constant of H-3 (d, *J* = 11.6 Hz), H-4 (dd, *J* = 15.6, 11.6 Hz) and H-5 (d, *J* = 15.6 Hz). The oxymethylene signals at δ_H_ 3.95, 3.85 ppm/δ_C_ 62.0 ppm, oxymethine at δ_H_ 5.74 ppm (d, *J* = 7.4 Hz)/δ_C_ 92.1 ppm and δ_H_ 5.02 ppm (d, *J* = 7.4 Hz)/δ_C_ 72.6 ppm, δ_H_ 3.75 ppm/δ_C_ 75.4 ppm, δ_H_ 3.74 ppm/δ_C_ 70.9 ppm, δ_H_ 3.57 ppm/δ_C_ 76.0 ppm indicated the glucose moiety. The α-configuration is deduced based on the chemical shift of anomeric carbon at C-1’ (δ_H_ 5.74 ppm/δ_C_ 92.1 ppm) and coupling constant (*J* = 7.4 Hz). Although, the sugar configuration is controversially (22) we signed according to the identical ^1^H NMR reported previously. The methylene signals at C-2’’ (δ_H_ 2.34 ppm/δ_C_ 34.4 ppm), C-3’’–C-12’’ (δ_H_ 1.25 ppm/δ_C_ 29.6 ppm), methyl group at C-14’’ (δ_H_ 0.86 ppm/δ_C_ 20.3 ppm) and C-15’’ (δ_H_ 0.88 ppm/δ_C_ 13.9 ppm) indicated the presence of branched saturated fatty acid. The low-field chemical shift of H-3 (δ_H_ 7.33 ppm/δ_C_ 141.6 ppm) and C-1 (δ_C_ 166.5 ppm) indicated the de-shielding effect based on the esterification of carbonyl group at C-1. Glycosylation at C-1’ position of glucose and C-1 of 4,4’-diaponeurosporenoic acid was deduced by the observation of HMBC correlation of H-1’ to C-1. The chemical shift of C-1’’ at δ_C_ 174.0 ppm indicated the esterification of carbonyl group **(Supplemental Table 8)**.

#### Supplemental Results SR2

**Characterization of µRNA species from MLBs**

To characterize RNA species from MLBs, µRNA isolated from MLBs (SA-MLB) were deep sequenced and compared to *S. aureus* µRNA (SA) and Δ*agr* *S. aureus* (SA-Δ*agr*) µRNA. We obtained approximately 3 million reads per sample of SA, ∆*agr* and SA-MLB RNA from each library sequenced in duplicate using Illumina dual-sided 250 base pair reads, covering the entire 15-200 base cut-off µRNA region. Sequenced reads were mapped onto reference sequences of *S. aureus* subsp. aureus N315 (accession number: NC_002745.2) and *S. aureus* subsp. aureus NCTC 8325 (accession number: NC_007795.1). These reads were further analysed using the µRNA analysis algorithms of SeqMan Pro ArrayStar and GenVision Pro followed by identification of at least 2-fold differentially abundant reads. We found 2828 different genomic locations enriched with RNA seq reads covering the entire genomic length. For this analysis we focused on µRNA reads obtained from non-coding intergenic regions and hence eliminated reads obtained from rRNA, antisense and coding regions. After elimination there were 186 genomic regions that were mapped with differentially abundant RNA seq reads on the *S. aureus* N315 genome **(Fig. 5A)**. Closer inspection revelaed many differentially abundant genomic locations covering known sRNA from *S. aureus* **(Supplemental Fig. 5B)**. We identified literature-known genomic locations for eight sRNA families comprising 106 individual sRNA from the *S. aureus* N315 genome (50) **(Supplemental Table 4)**. NGS reads were manually curated and mapped on all identified sRNA for SA, SA-MLB, and SA-Δ*agr.* Following this we calculated the percentage reads from the RNA seq data **(Supplemental Fig. 5B, 5C)**. Of eight sRNA families analysed, Rsa X17, Rsa OI, OL, OM, OP, OV, Sau-50, Sau-6053, WAN01CC8T, Teg 69, 70, 47, 91, 61 were only present in SA. However, abundant reads for Rsa A, Rsa X12, Sau-64 were only present in SA-Δ*agr*. Interestingly, reads matching with Rsa C, Sbr C, RNAIII, and WAN01CC66-rc were abundantly present in SA and SA-MLB and absent in SA-Δ*agr*. All other sRNAs were differentially abundant in reads of SA, SA-MLB and SA-Δ*agr* **(Supplemental Fig. 5B, Supplemental Table 4).**

*spr* genes such as *spr* A-G (51) were absent from our analysis of SA, SA-MLB, SA-∆*agr* **(Supplemental Table 4)**. Rsa A, C, E, F, H and I were differentially abundant in SA, SA-MLB and SA-∆*agr.* Rsa A which is σ^B^ dependent was only present in SA-∆*agr,* Rsa F which is strain specific to *S. aureus* showed relatively less presence in SA-MLB and SA-∆*agr* as compared to SA. Rsa C, which carries the small ORF as RNA III, were exclusively abundant only in SA-MLBs. Consistent with the literature, Rsa E, which is regulated by the quorum sensing system, was decreased in SA-∆*agr* (52). Rsa O genes were also differentially abundant (53, 54). Rsa OG and Rsa OT levels were low in SA-MLB as compared to SA and SA-∆*agr*. None of the Sau genes were present in SA-MLBs. Of the five σ^B^ regulated genes, *sbr* (55) only *sbr* B,C and D were found in our analysis. There was no *sbr* B and D in SA-MLBs, while *sbr* C was only expressed in SA and SA-MLBs and were completely dependent on SA-∆*agr*. Levels of WAN01CC66-rc were high in SA-MLBs as compared to SA and absent in SA-∆*agr*. Teg genes were also identified in our analysis of SA, SA-MLB and SA-∆*agr* µRNA. Teg 57 was only present in SA-MLBs but absent in SA-∆*agr* and SA. Riboswitches such as Teg 1 and Teg 76 were absent in all the three µRNA. RNAIII, one of the best characterized regulators, was found to be SA-∆*agr* dependent and exclusively present within SA-MLBs **(Supplemental Fig. 5B, Supplemental Table 4)**

Levels of RNA seq reads of sbr-C, Rsa-C, RNAIII and WAN01CC66-rc were AGR dependent and entirely present in SA and SA-MLBs **(Supplemental Fig. 5C, 5D)** making them important candidates for binding to CDN and activation of STING in human macrophages. Hence, we further analysed read coverage into genomic locations of Rsa C, RNAIII and WAN01CC66-rc **(Supplemental Fig 5D).** Interestingly, parts of sRNA locus were enriched in RNA seq reads of SA-MLBs as compared to SA and completely absent in SA-∆*agr* **(Supplemental Fig. 5D).** For example, RsaC reads (679912-680136) were exclusively present in SA-MLB whereas reads covering 680489-680602 were present in both SA and SA-MLB **(Supplemental Fig. 5D, Supplemental Table 4)**. These results indicate that SA MLBs contain distinct parts of sRNA species that are differentially enriched from the originating bacteria, indicating the possibility of specific processing and packaging of sRNA species within MLBs of *S. aureus*.
