## Supplemental figures and will be used for the link to the file on the preprint site for "Gram-positive bacteria secrete RNA aptamers to activate human STING for IL-1β release"

### Supplemental Fig. 1

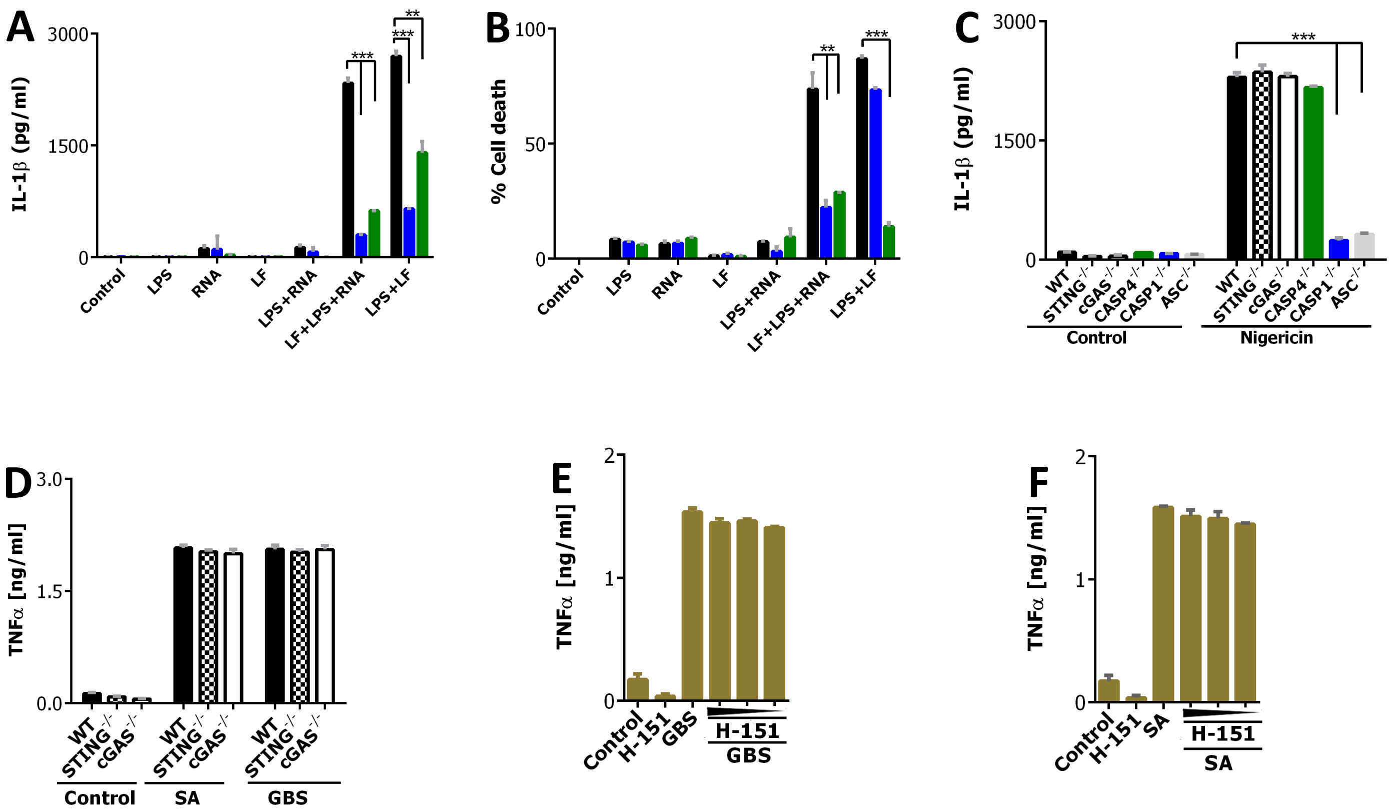

### Supplemental Fig. 2

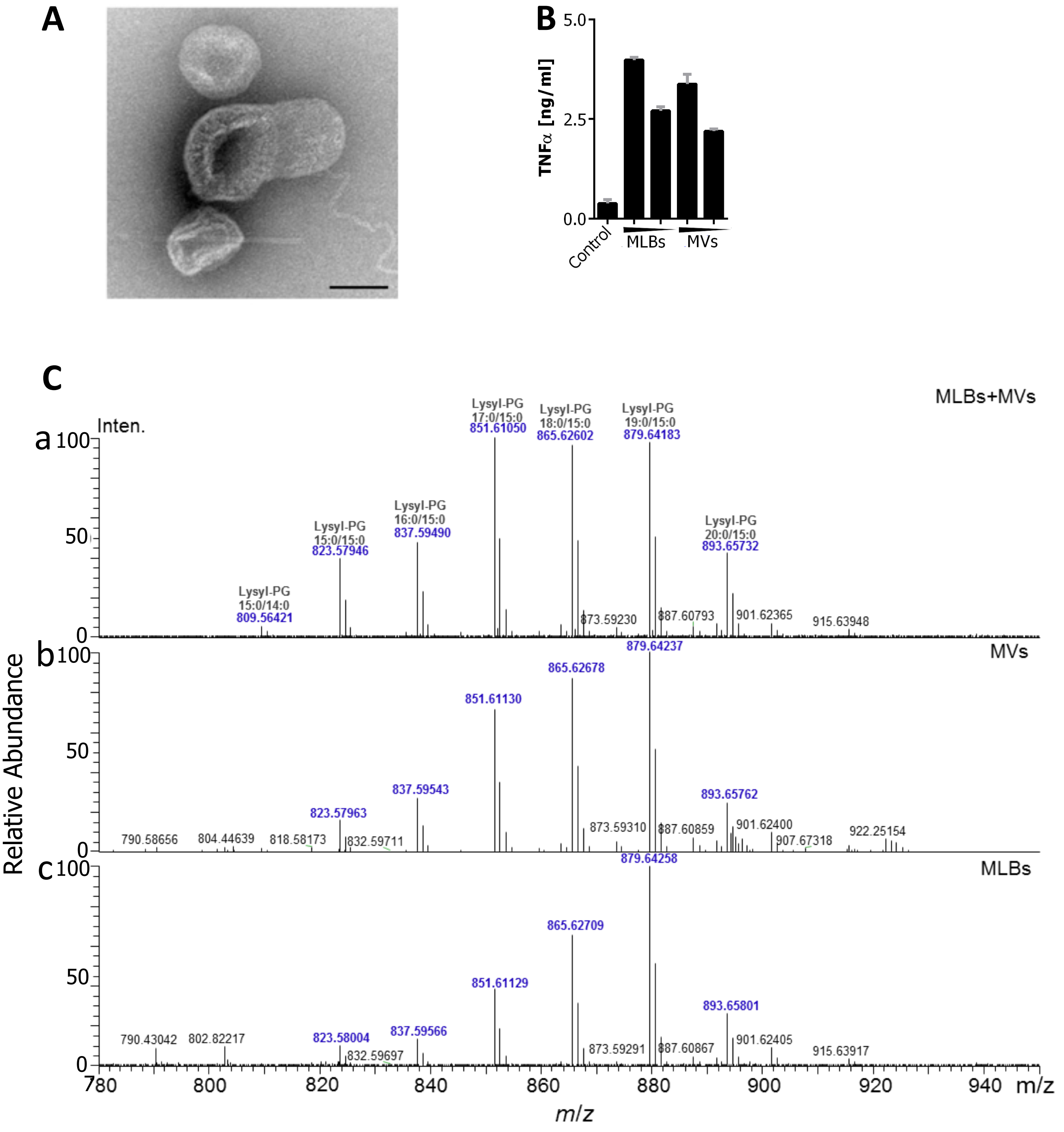

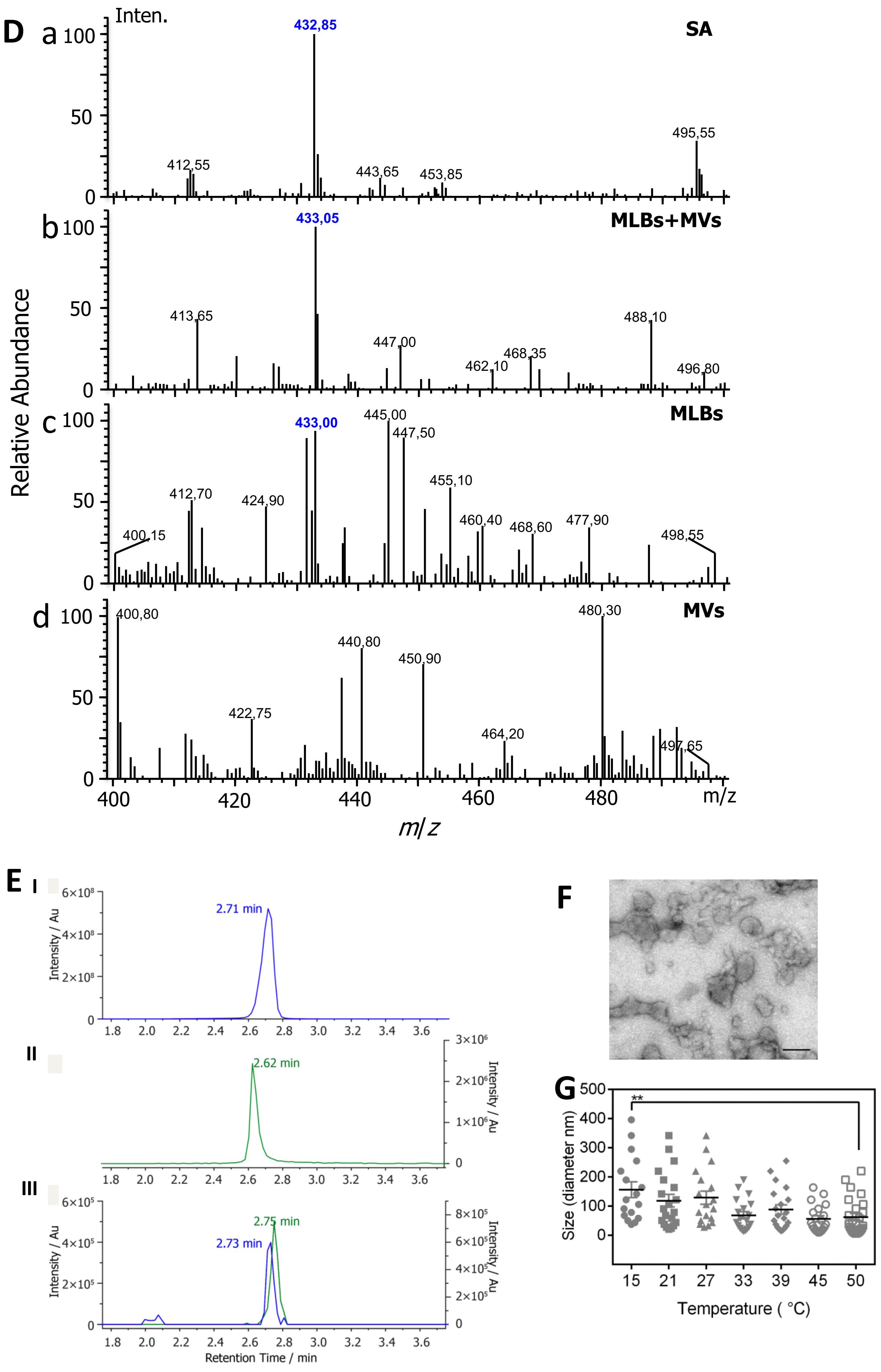

D
a
b
c
d
Inten.
Relative Abundance
SA
MLBs+MVs
MLBs
MVs
432,85
412,55
443,65
453,85
495,55
433,05
413,65
447,00
462,10
468,35
488,10
496,80
433,00
445,00
447,50
455,10
412,70
424,90
400,15
460,40
468,60
477,90
498,55
400,80
480,30
440,80
450,90
422,75
464,20
497,65
m/z
E
I
II
III
2.71 min
2.62 min
2.75 min
2.73 min
Intensity / Au
Retention Time / min
F
G
**
Size (diameter nm)
Temperature ( °C)
15
21
27
33
39
45
50

### Supplement Fig. 3

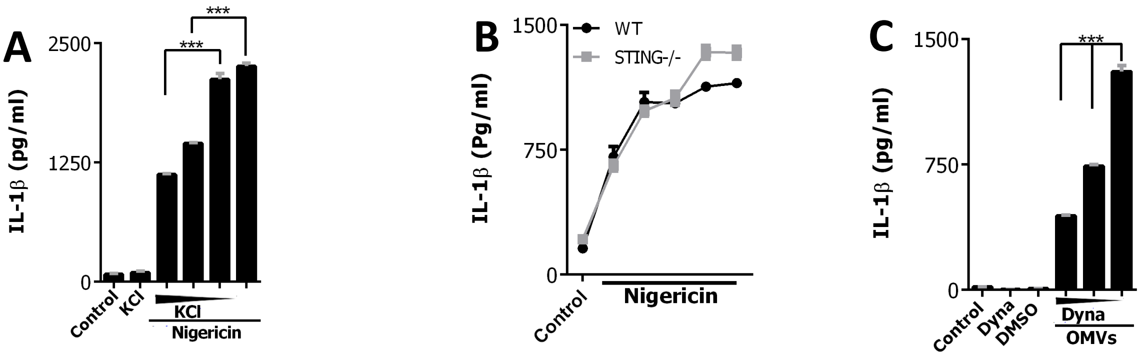


### Figure Supplement S4

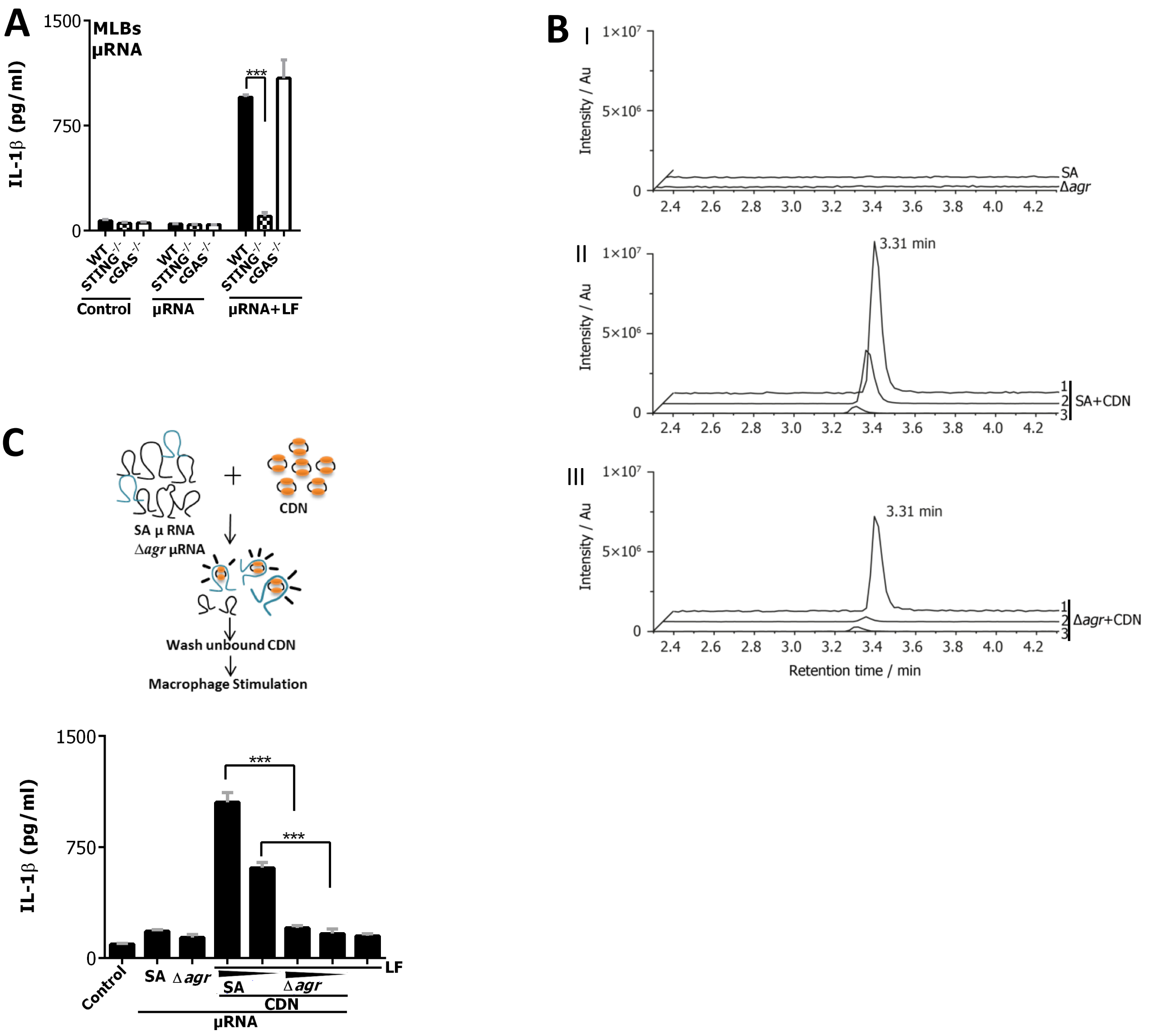

### Supplement Fig. 5

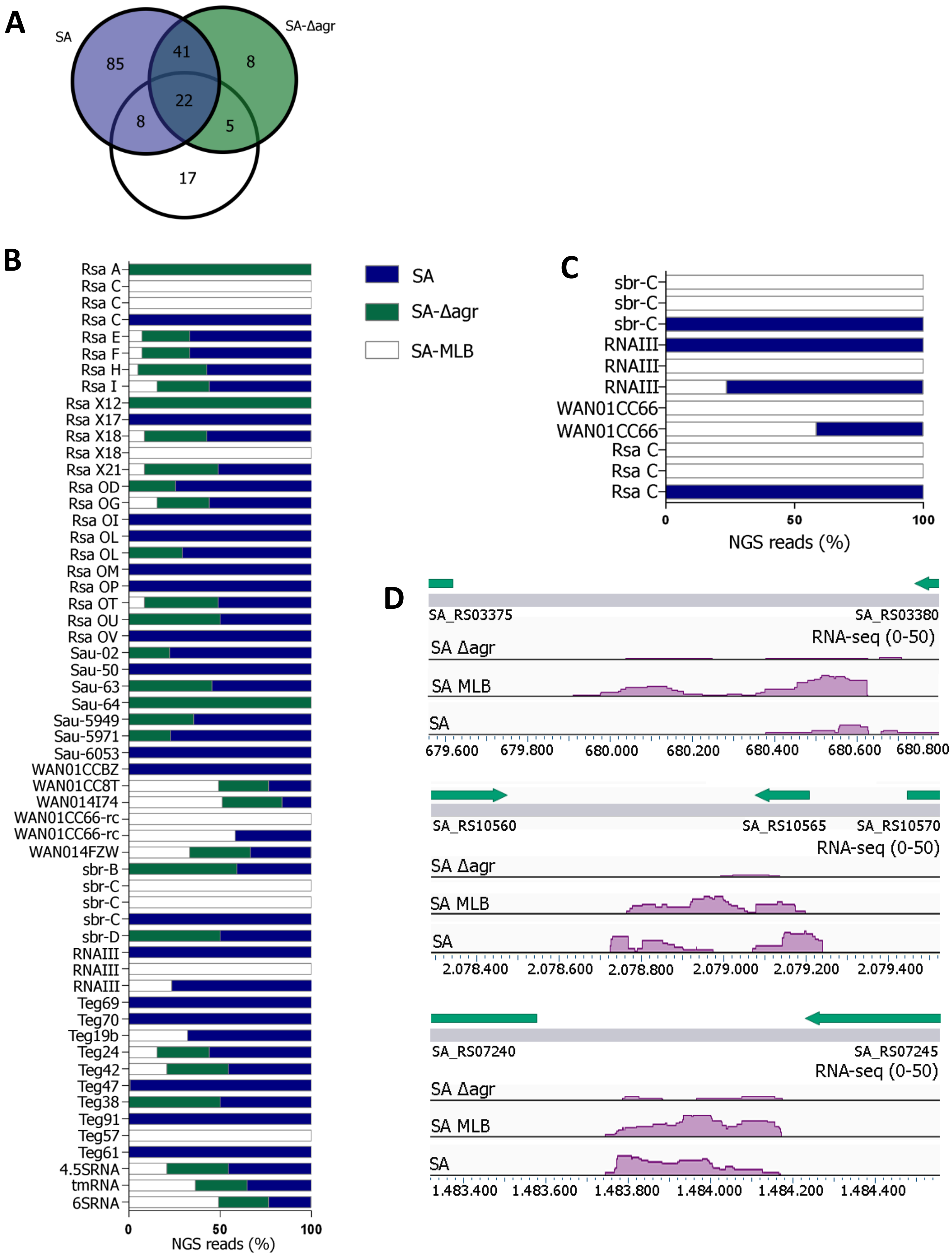

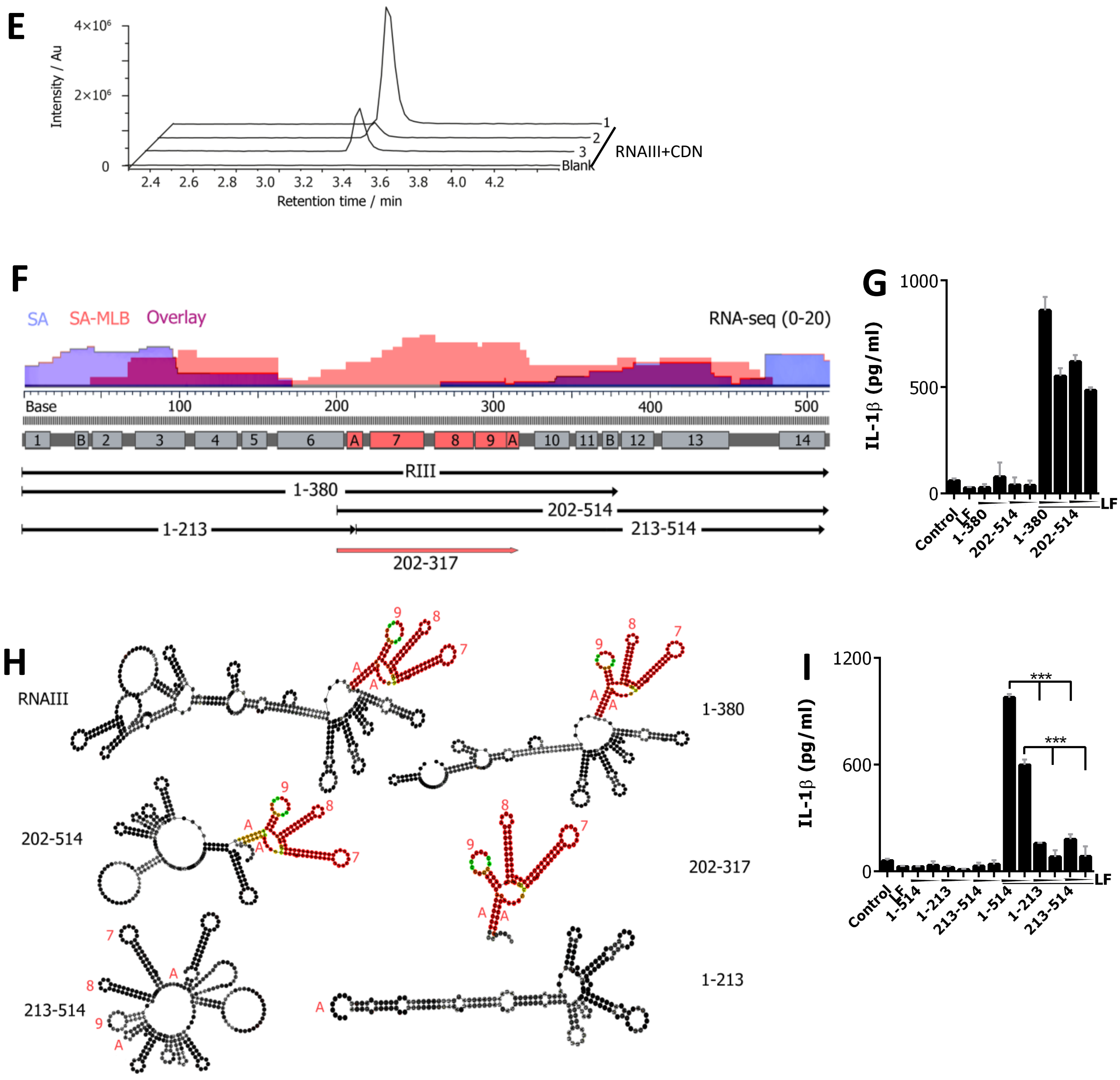
E
4×10⁶
2×10⁶
0
Intensity / Au
2.4 2.6 2.8 3.0 3.2 3.4 3.6 3.8 4.0 4.2
Retention time / min
1
2
3
Blank
RNAIII+CDN
F
SA SA-MLB Overlay
RNA-seq (0-20)
Base 100 200 300 400 500
1 B 2 3 4 5 6 A 7 8 9 A 10 11 B 12 13 14
RIII
1-380
202-514
1-213
213-514
202-317
G
IL-1β (pg/ml)
1000
500
0
Control
LF
1-380
202-514
1-380
202-514
LF
H
RNAIII
1-380
202-514
202-317
213-514
1-213
I
IL-1β (pg/ml)
1200
600
0
***
***
Control
LF
1-514
1-213
213-514
1-514
1-213
213-514
LF

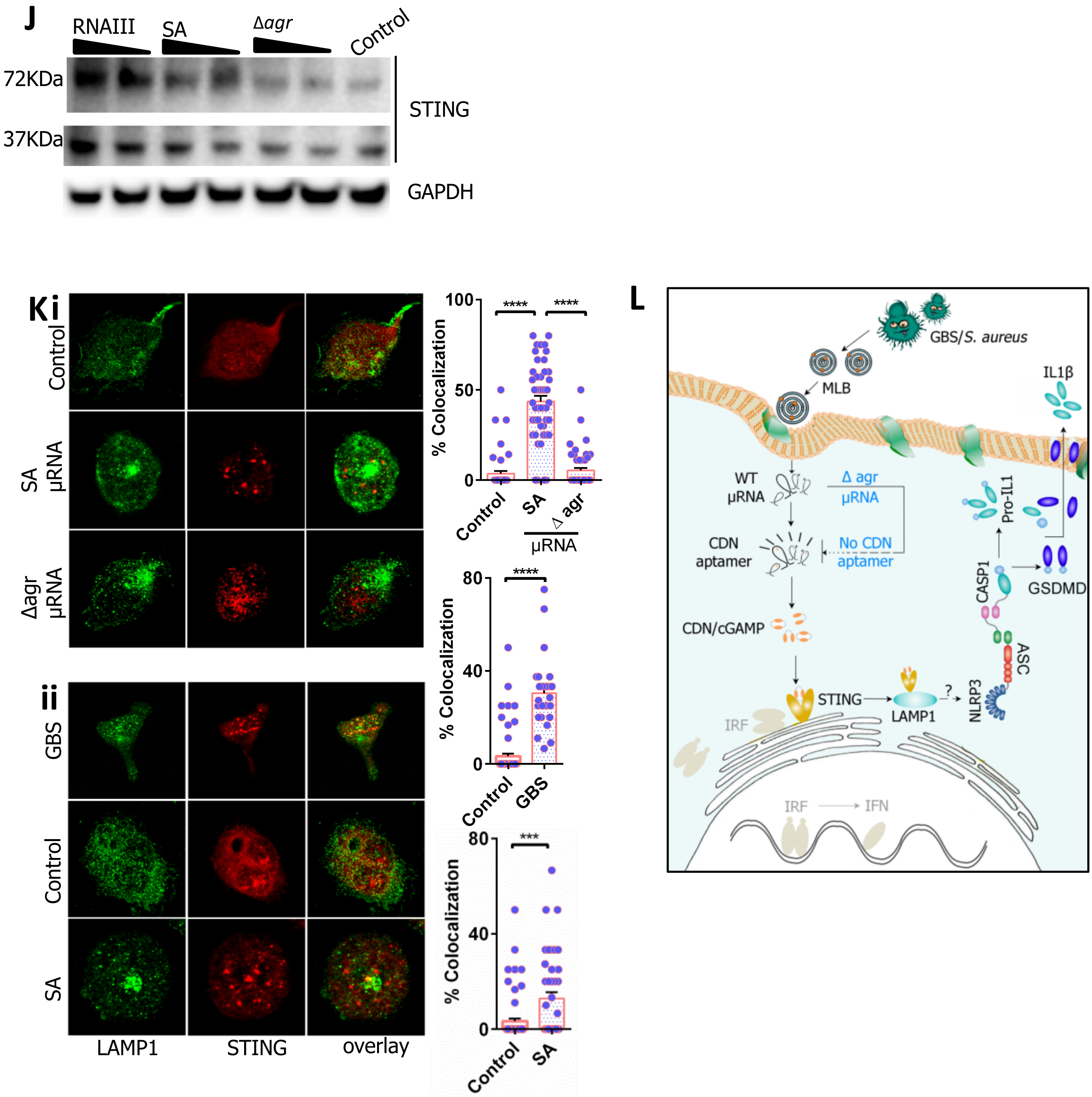
J
RNAIII
SA
Δagr
Control
72KDa
37KDa
STING
GAPDH
Ki
Control
SA µRNA
Δagr µRNA
ii
GBS
Control
SA
LAMP1
STING
overlay
% Colocalization
100
50
0
****
Control
SA
Δ agr
µRNA
80
40
GBS
***
L
GBS/S. aureus
MLB
IL1β
WT µRNA
Δ agr µRNA
CDN aptamer
No CDN aptamer
Pro-IL1
CASP1
GSDMD
CDN/cGAMP
ASC
STING
LAMP1
NLRP3
IRF
IFN

#### Supplement Fig. 6

### A

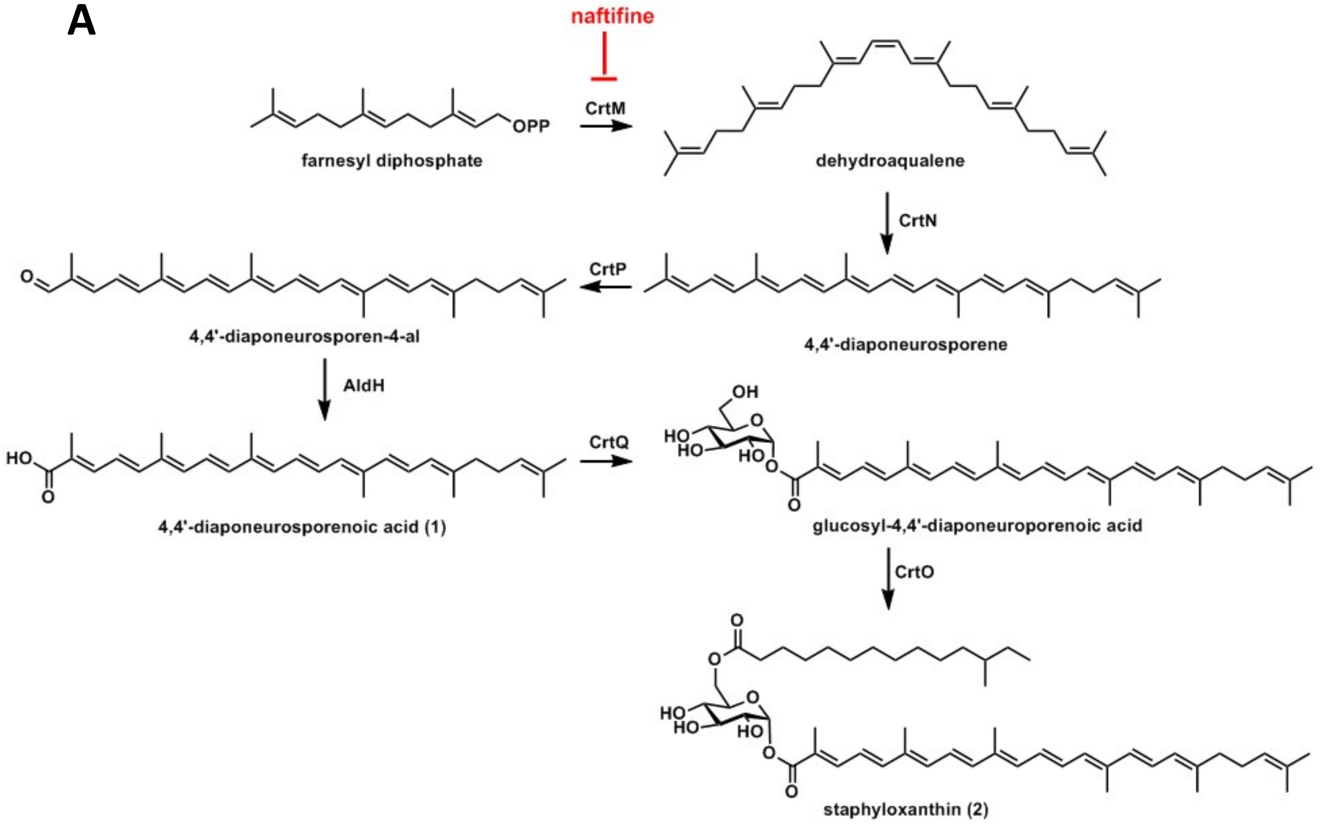

### Supplement Fig. 7

A

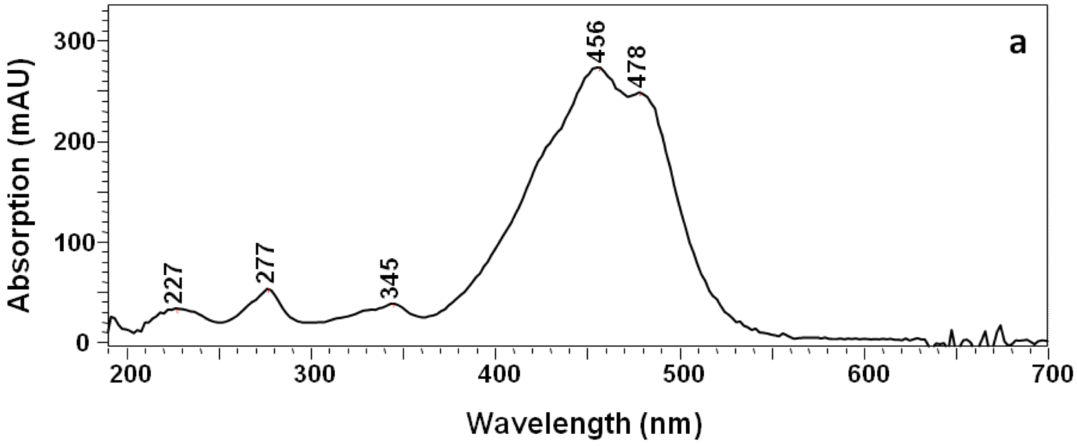


B

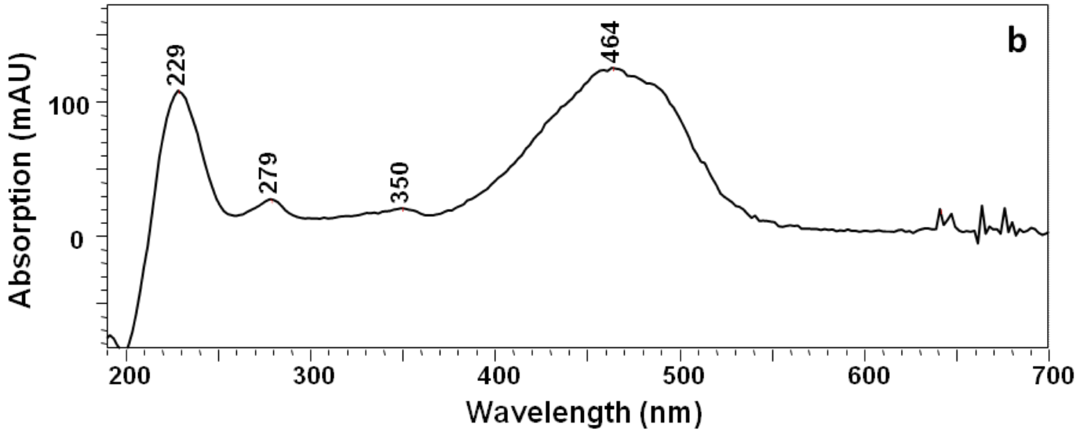


C

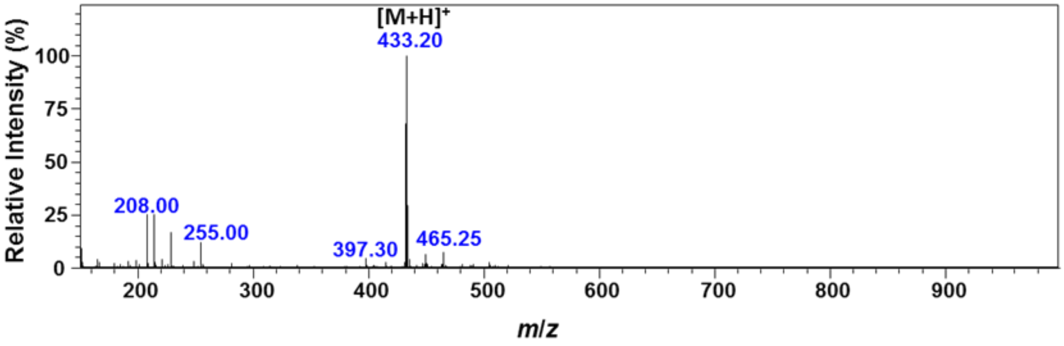

D

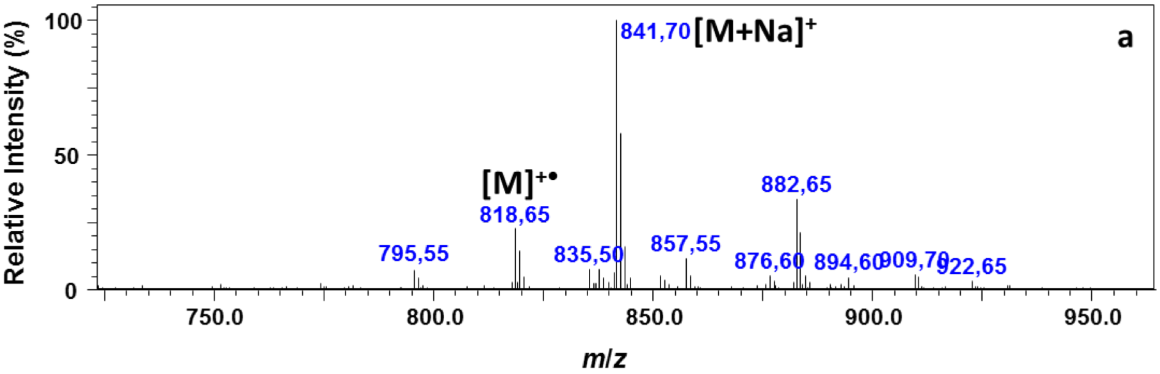
a
Relative Intensity (%)
100
50
0
841,70 [M+Na]+
[M]+•
818,65
882,65
795,55
835,50
857,55
876,60
894,60
909,70
922,65
750.0
800.0
850.0
900.0
950.0
m/z

E

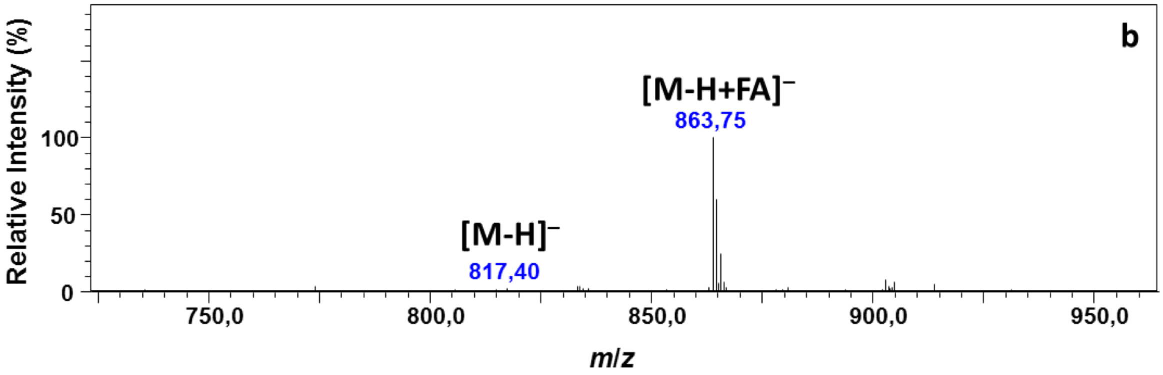
b
Relative Intensity (%)
100
50
0
[M-H+FA]−
863,75
[M-H]−
817,40
750,0
800,0
850,0
900,0
950,0
m/z

F

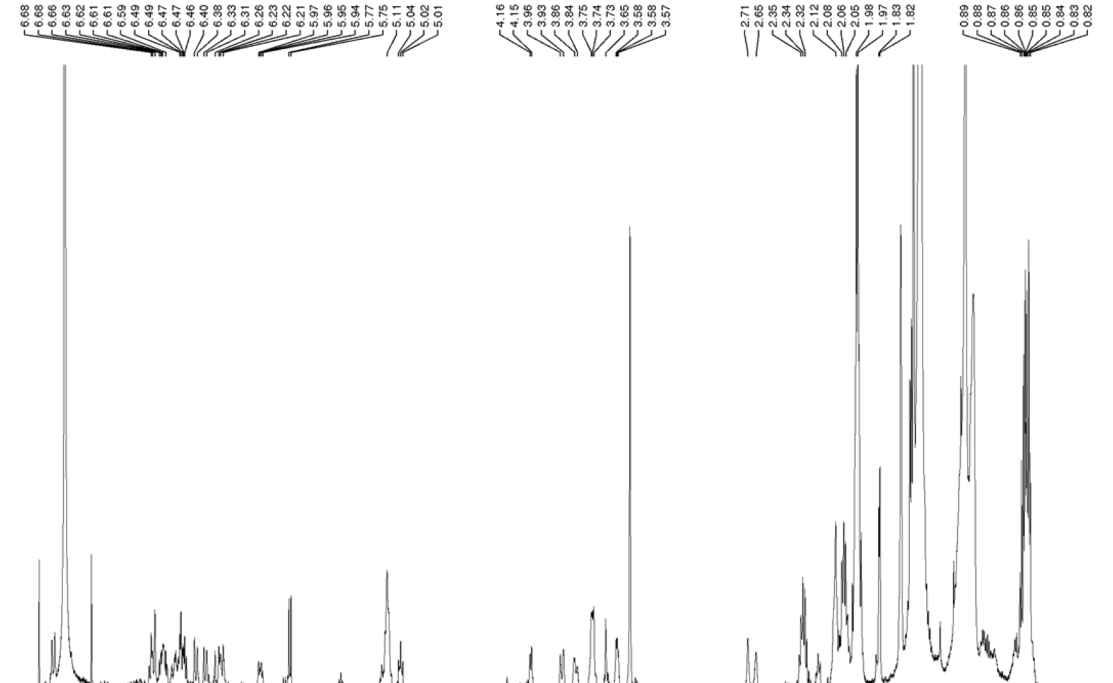
6.68 6.68 6.66 6.63 6.62 6.61 6.61 6.59 6.49 6.47 6.46 6.40 6.38 6.33 6.31 6.26 6.23 6.22 6.21 5.97 5.96 5.95 5.94 5.77 5.75 5.11 5.04 5.02 5.01
4.16 4.15 3.96 3.93 3.86 3.84 3.75 3.74 3.73 3.65 3.58 3.58 3.57
2.71 2.65 2.35 2.34 2.32 2.12 2.08 2.06 2.05 1.98 1.97 1.83 1.82
0.89 0.88 0.87 0.86 0.86 0.86 0.85 0.85 0.84 0.83 0.82
7.5
7.0
6.5
6.0
5.5
5.0
4.5
4.0
3.5
3.0
2.5
2.0
1.5
1.0
ppm

G

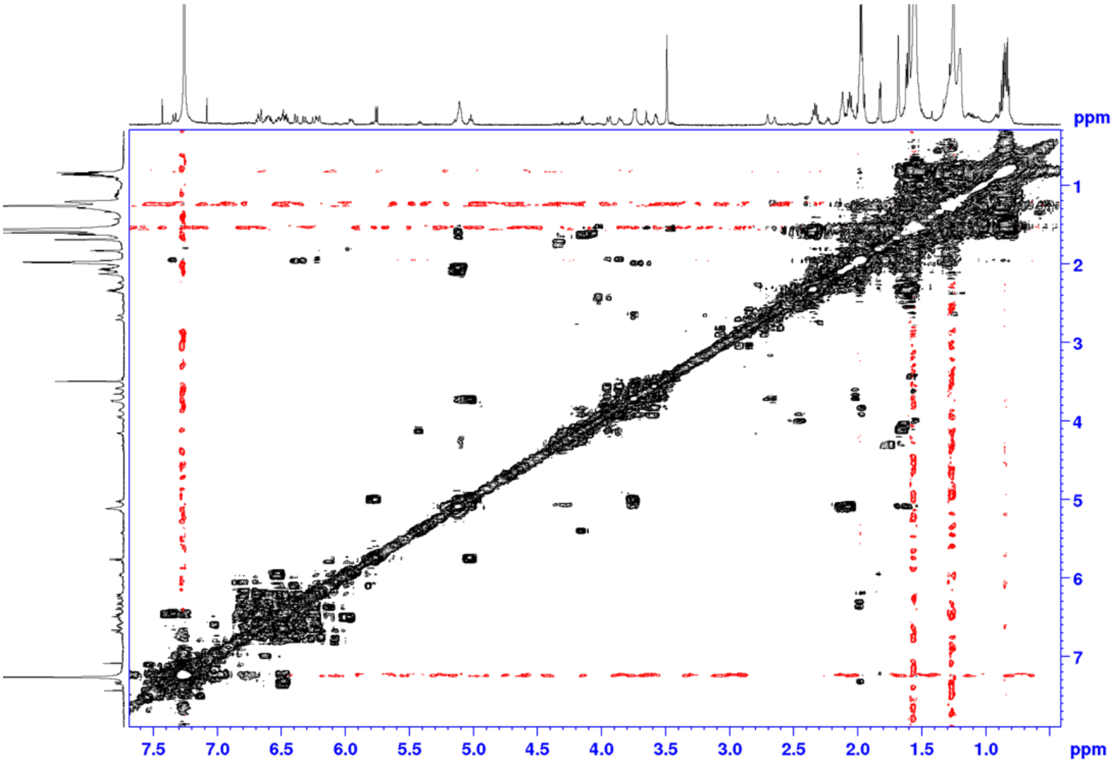
ppm
1
2
3
4
5
6
7
7.5
7.0
6.5
6.0
5.5
5.0
4.5
4.0
3.5
3.0
2.5
2.0
1.5
1.0
ppm

H

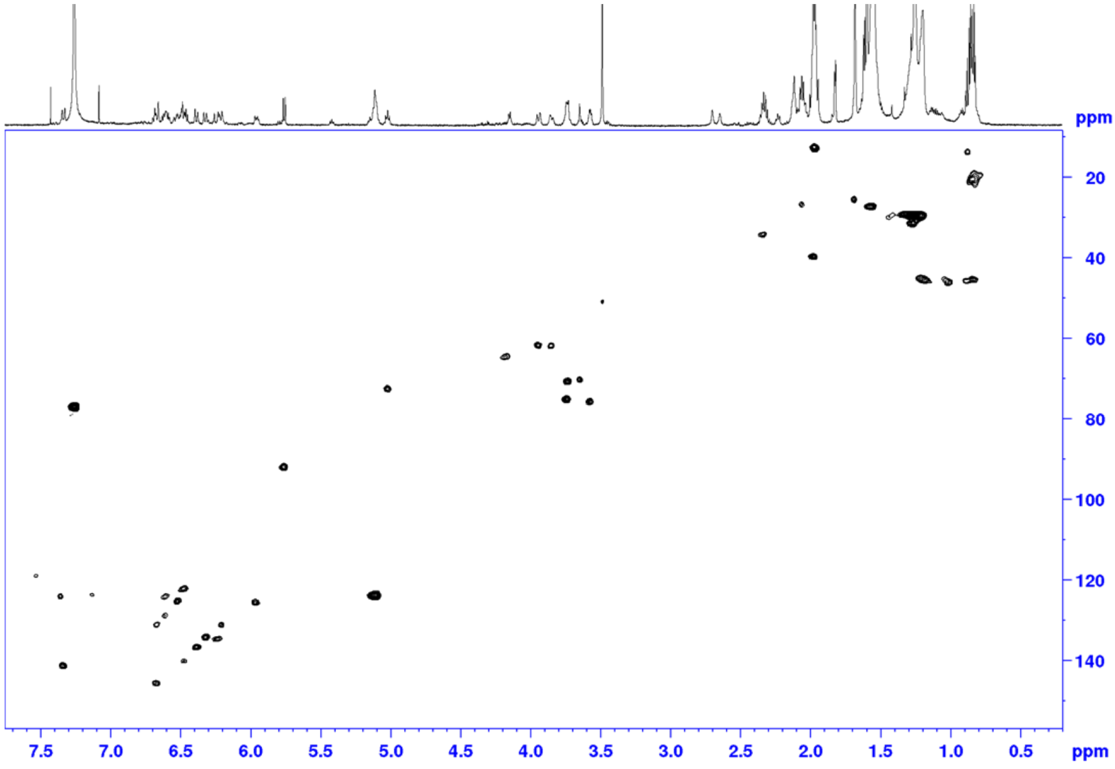
ppm
20
40
60
80
100
120
140
7.5
7.0
6.5
6.0
5.5
5.0
4.5
4.0
3.5
3.0
2.5
2.0
1.5
1.0
0.5
ppm

I

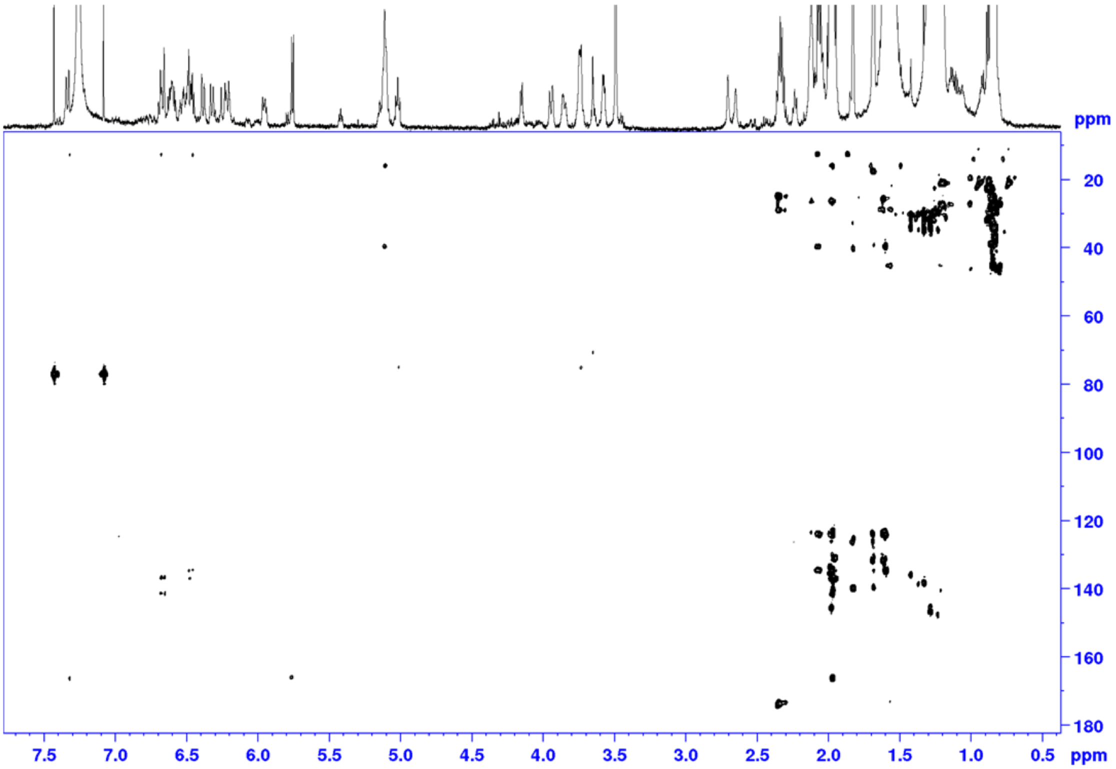


J

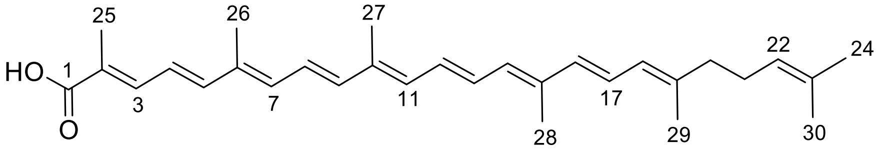


4,4'-diaponeurosporenoic acid (**1**)

K

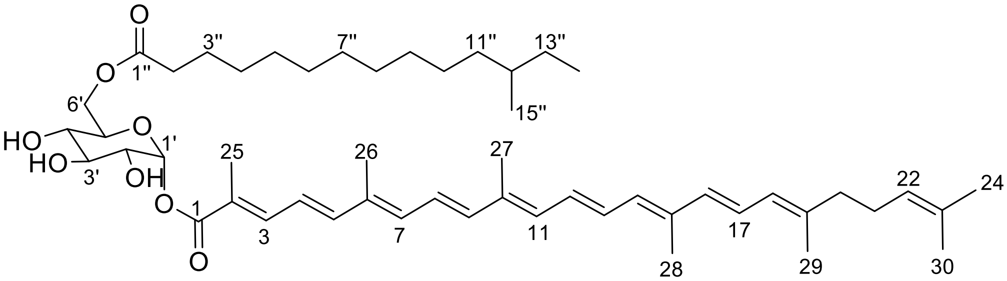


staphyloxanthin (**2**)
